# Unveiling allopolyploidization-driven genome duplications through progressive analysis of deep genome skimming data

**DOI:** 10.1101/2024.09.24.614835

**Authors:** Si-Yu Xie, Xiao-Hua Lin, Jun-Ru Wang, Dai-Kun Ma, Yu Zhang, Chao Xu, Hong Ma, Pan Li, Duo-Yuan Chen, Xin Zhong, Bin-Jie Ge, Richard G.J. Hodel, Liang Zhao, Bin-Bin Liu

## Abstract

Whole-genome duplication (WGD) events are widespread across the Web of Life (WoL). Given the prevalence of WGDs in the polyploid *Prunus* (Rosaceae), this economically- and agriculturally-important angiosperm lineage provides an excellent model for exploring this mode of reticulation. We used the polyploid *Prunus* to demonstrate a progressive strategy for analyzing Deep Genome Skimming (DGS) data in the presence of WGDs. Phylogenomic discordance analyses indicated that allopolyploidization, rather than Incomplete Lineage Sorting (ILS), played a dominant role in the origin and dynamics of polyploid *Prunus*. This study underscores how a progressive strategy to identify WGD events at different depths in a phylogenetic tree reveals the nuances of evolutionary mechanisms driving allopolyploidization. We inferred that the continued uplift of the Himalayas from the Middle to Late Miocene drove the rapid diversification of the Eastern Asia endemic *Maddenia* clade, by facilitating frequent hybridization and allopolyploidization, specifically introgression between the Himalayas-Hengduan and the Central-Eastern China clades.

## Introduction

The evolutionary history of plant lineages is shaped by mechanisms operating at deep and shallow phylogenetic levels. At deeper phylogenetic scales, processes like allopolyploidy and whole-genome duplication (WGD) have driven diversification^1–6^. Allopolyploidy, one particular case of WGD—the merging of distinct genomes from different species through hybridization and polyploidization—has been a driving force in the evolution of many plant lineages^2^. This process increases genetic diversity and leads to new species with novel traits, promoting adaptation to diverse environments^7^. At shallower phylogenetic scales, mechanisms such as introgression and horizontal gene transfer are pivotal^8–10^. Introgression, the transfer of genetic material between species through repeated hybridization and backcrossing, can rapidly introduce novel traits and enable plants to adapt to new environments or changing conditions^11^. Introgression may blur species boundaries and inhibit species delimitation^12^. Allopolyploidy and/or introgression illustrate how ancient and recent processes interact to shape the diversity and complexity of plant life on Earth.

The plum genus *Prunus* s.l. (Rosaceae), which includes the widely recognized and economically important plum trees, along with cherries, almonds, peaches, and apricots, is a remarkable example of evolutionary complexity and diversification in the plant kingdom^13–15^. *Prunus* s.l. is particularly notable for its hypothesized WGD events, especially within the polyploid racemose clade (*Prunus* subg. *Padus*: hereafter referred to as the polyploid *Prunus*)^16–18^. Multiple copies of the nuclear *At103* gene provide clear evidence of the allopolyploid origin of polyploid *Prunus*^18^, suggesting that hybridization between species, followed by chromosome doubling, has led to novel genomic combinations. However, the specific details of this network-like evolution, including the identification of maternal and paternal participants in allopolyploidy and/or hybridization, remain unresolved. Additionally, a recent phylogenomic study based on 446 single-copy nuclear (SCN) loci detected four hybridization events, highlighting extensive gene flow in the diversification of the *Maddenia* group^19^. We propose that the polyploid *Prunus* lineage has likely experienced extensive introgression events, wherein genes are transferred between species through hybridization and backcrossing. Such introgression has facilitated the exchange of beneficial traits, including disease resistance and environmental adaptability, across different species within the genus^20,21^.

Despite the importance of allopolyploidy as a significant driver of species diversification and environmental adaptation^3,5^, there remains a lack of specialized computational tools or pipelines dedicated to elucidating the reticulate evolutionary processes associated with polyploidization. Traditional approaches to estimate WGD events have relied on synteny analysis of genomic data, utilizing tools such as MCScanX^22^ or SynMap^23^. However, these methods require high-quality genome assemblies and accurate gene annotations. More recently, WGD detection has expanded to include analyses of transcriptome sequencing (RNA-Seq) data, specifically employing techniques like synonymous substitution rate (Ks) analysis between paralogous gene pairs^24^ and phylogenetic tree reconciliation between gene trees and species trees^2,25–30^. Nevertheless, acquiring comprehensive genomic and transcriptomic datasets for specific lineages remains challenging, notably due to the stringent requirements for liquid nitrogen-preserved materials for RNA-Seq. Recent advancements in target enrichment (Hyb-Seq)^31^ and Deep Genome Skimming (DGS)^32^ sequencing technologies offer promising avenues for utilizing ancient DNA extracted from museum and herbarium collections^33,34^. These techniques have rapidly gained traction within the systematics community, bridging gaps in taxon sampling. Specifically, the Tree2GD method^30^, initially designed for RNA-Seq data^28,29,35–41^, has emerged as a particularly suitable program for WGD identification within Hyb-Seq and DGS datasets.

In this study, we use the polyploid *Prunus* clade as a model: 1) to propose a general pipeline for detecting WGD events, integrating multiple methods to elucidate the evolutionary dynamics driven by polyploidization, specifically accommodating a Tree2GD approach for DGS data involving predesigned SCN loci; 2) to delineate the evolutionary processes that influence the formation of duplicated genomes and contribute to phylogenetic discordance within polyploid *Prunus*; 3) to investigate the evolutionary mechanisms responsible for the scarcity of informative sites capable of clarifying phylogenetic relationships within the *Maddenia* group.

## Results

### Assembly of plastomes and nuclear orthology inference for generating two datasets

After screening three genomes, we identified 1,806 SCN loci for use in the following phylogenomic analyses. The genomic dataset was comprised of 93 individuals with DGS data and 21 with RNA-Seq data. An analysis of all 114 samples revealed variable retrieval of SCN loci, ranging from 1,355 (accounting for 75% of the total identified loci) to the complete set of 1,806 loci (representing 100%; Supplementary Fig. 1). We derived two distinct datasets—Monophyletic Outgroup (MO) and RooTed ingroup (RT)—by employing various orthology inference methods on the non-chimeric SCN sequences. These methods generated 1,622 and 2,268 genes corresponding to the MO and RT datasets, respectively. Specifically, MO dataset alignment comprised 2,310,500 characters, and the RT dataset constituted 3,193,261 characters.

We successfully assembled 83 whole plastomes from DGS data, demonstrating the efficacy of this method for plastome assembly. However, for the remaining 10 out of the 93 DGS datasets, we obtained only the plastid coding sequences (plastid CDS) due to insufficient sequencing coverage. These 83 plastid genomes have been submitted to GenBank (Supplementary Table 1). Consistent with our previous findings^8,42,43^, RNA-Seq data were suitable only for capturing exon-related reads, a postulation further supported by our recovery heatmap (Supplementary Fig. 2).

### Nuclear/plastid backbone of *Prunus* s.l. and the gene tree conflict analyses

Phylogenetic trees were estimated employing both concatenation-(IQ-TREE2 and RAxML) and coalescent-based (ASTRAL-III) methods across three datasets, encompassing two SCN (MO and RT) and one plastid CDS datasets. This phylogenetic inference yielded nine trees (six nuclear and three plastid CDS ones), consistently revealing largely consistent topologies among the major clades (Fig. 1; Supplementary Figs. 3–11). For downstream analyses, we used the species tree estimated by ASTRAL-III based on the MO dataset for simplicity (Fig. 1; Supplementary Fig. 3).

**Figure 1.**
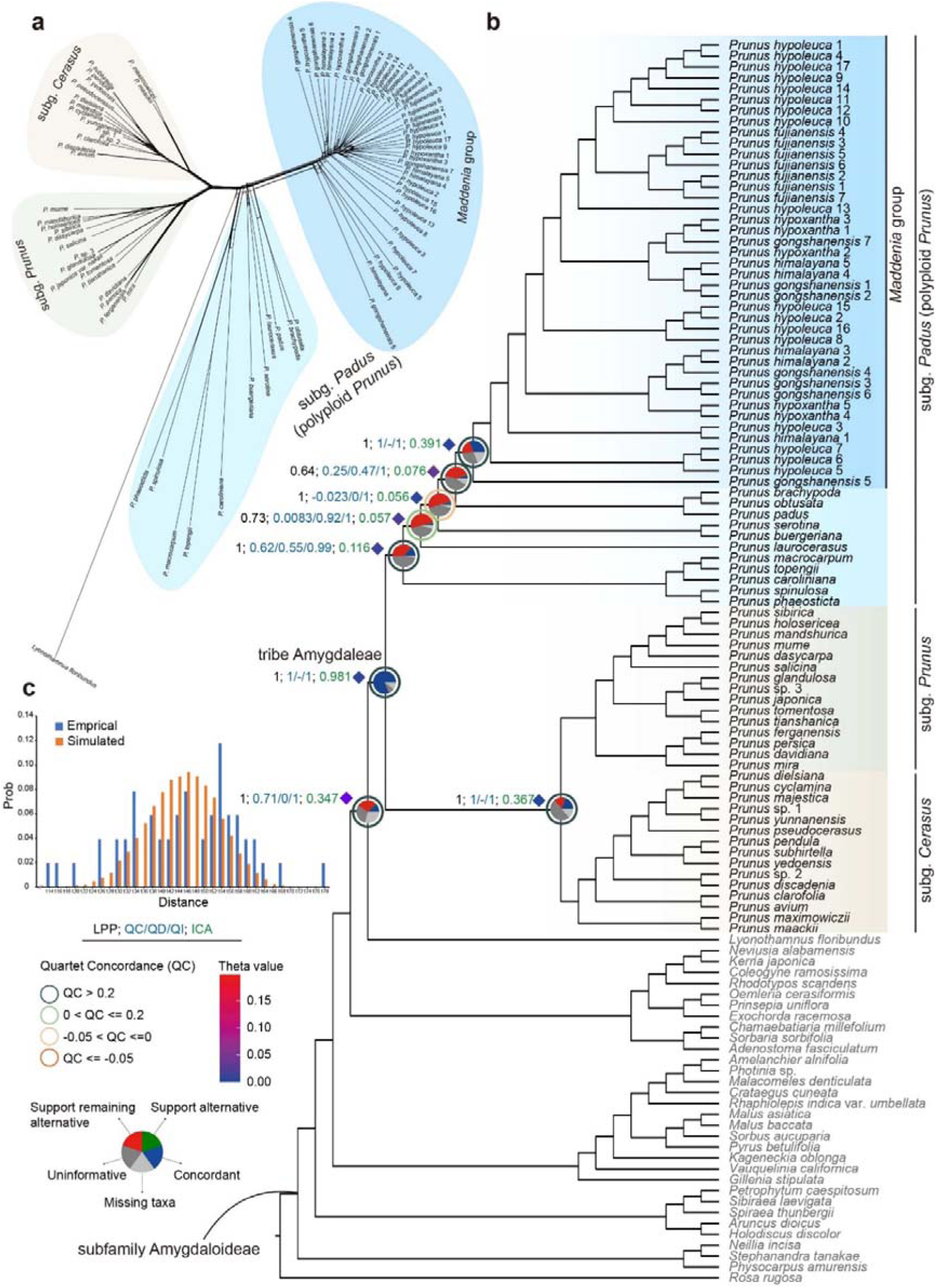
The topology of the subfamily Amygdaloideae. **a** Supernetwork inferred with SplitsTree based on SCN genes from the MO dataset in the framework of the subfamily Amygdaloideae, where parallelograms indicate incongruences among SCN genes. The *Maddenia* group is delineated in dark blue, while other species within the polyploid *Prunus* are illustrated in light blue. Additionally, *P.* subg. *Prunus* is represented in green, and *P.* subg. *Cerasus* is depicted in yellow. **b** Species tree of polyploid *Prunus* in the framework of Amygdaloideae inferred from ASTRAL-III using 1,622 nuclear MO orthologs. Pie charts on the nodes represent the following data: the proportion of gene trees that support that clade (blue), the proportion that supports the main alternative bipartition (green), the proportion that supports the remaining alternatives (red), the proportion (conflict or support) that have less than 50% BS (dark grey), and the proportion that have missing taxa (light grey) (details refer to Supplementary Fig. 12). The color of the circle around the pie chart represents the value range of QC, where QC > 0.2 is painted in dark green, 0 < QC ≤ 0.2 is painted in light green, -0.05 < QC ≤ 0 is painted in yellow, and QC ≤ -0.05 is painted in red. The three different colored support values alongside the focused nodes were presented from left to right: LPP inferred by ASTRAL-III (labeled in black; details refer to Supplementary Fig. 3), values for QC/QD/QI estimated from QS analysis (e.g., 1/-/1, labeled in blue; details refer to Supplementary Fig. 14), and ICA scores (labeled in green; details refer to Supplementary Fig. 13). Additionally, the population mutation parameter theta values inferred from the MuCCo analysis are shown above branches using colored diamond (details refer to Supplementary Fig. 26). **c** Distribution of tree-to-tree distance between empirical gene trees and the ASTRAL species tree, compared to those from the coalescent simulate.

*Prunus* s.l. was consistently identified as monophyletic across nine phylogenetic trees—six nuclear and three plastid CDS ones (SH-aLRT/ ultrafast bootstrap support (UFBoot) = 100/100; Bootstrap support (BS) = 1; Local posterior probabilities (LPP) = 1; Fig. 1; Supplementary Figs. 3–11). However, the closely related sister group indicated substantial cytonuclear discordance. The sister relationship between *Prunus* s.l. and the tribe Lyonothamneae was supported across six nuclear trees (Supplementary Figs. 3–8), although only 177 of 668 SCN gene trees were concordant with the species tree according to the *phyparts* analysis (internode certainty all (ICA) = 0.347; Supplementary Figs. 12 and 13), yet it received strong support from the Quartet Sampling (QS) analysis (0.7/0/1; Supplementary Fig. 14). Conversely, in all three plastid trees, the tribe Sorbarieae was recovered as the group most closely related to *Prunus* s.l. (Supplementary Figs. 9–11). However, this sister relationship garnered support from only three of 38 informative plastid gene trees (Supplementary Fig. 15) and displayed counter-support from the ICA value (-0.041; Supplementary Fig. 16). The QS analysis further highlighted this discrepancy, showing counter-support (-0.04/0.76/0.72; Supplementary Fig. 17) and suggesting a strong majority of quartets favored an alternative discordant quartet arrangement histories.

Consistent with the contemporary infrageneric taxonomic classification of *Prunus* s.l.^44^, three major clades were identified, each corresponding to a distinct subgenus: *P.* subg. *Prunus*, *P.* subg. *Cerasus*, and *P.* subg. *Padus* (Fig. 2a). Given the prevalence of polyploidy within *P.* subg. *Padus*, this subgenus, often characterized by distinctive racemose inflorescences, will hereafter be referred to as polyploid *Prunus*. All nine phylogenetic trees, inferred from both nuclear MO (1,622 SCN genes) and RT (2,268 SCN genes) datasets (Supplementary Figs. 3–8), along with those from plastid data (Supplementary Figs. 9–11), consistently delineated the relationships among these three subgenera (Fig. 2b). Notably, the sister relationship between *P.* subg. *Prunus* and *P.* subg. *Cerasus* was strongly supported, as evidenced by concordance in 324 of 592 SCN gene trees (ICA = 0.367; Supplementary Figs. 12 and 13) and 14 out of 24 plastid CDSs (ICA = 0.245; Supplementary Figs. 15 and 16), as well as strong QS support in the SCN and plastid (1/-/1; Supplementary Figs. 14 and 17) datasets. Then, these two subgenera are together sister to the polyploid *Prunus* with strong support (SH-aLRT/UFBoot = 100/100; BS = 1; LPP = 1; Figs. 1 and 2b; Supplementary Figs. 3–11) and concordance observed in 1,317 out of 1,321 informative SCN genes (ICA = 0.981; Supplementary Figs. 12 and 13) and 23 out of 29 plastid CDSs (ICA = 0.453; Supplementary Figs. 15 and 16) from the *phyparts* analysis, along with full QS support (1/-/1) in the SCN (Supplementary Fig. 14) and plastid (Supplementary Fig. 17) datasets.

**Figure 2.**
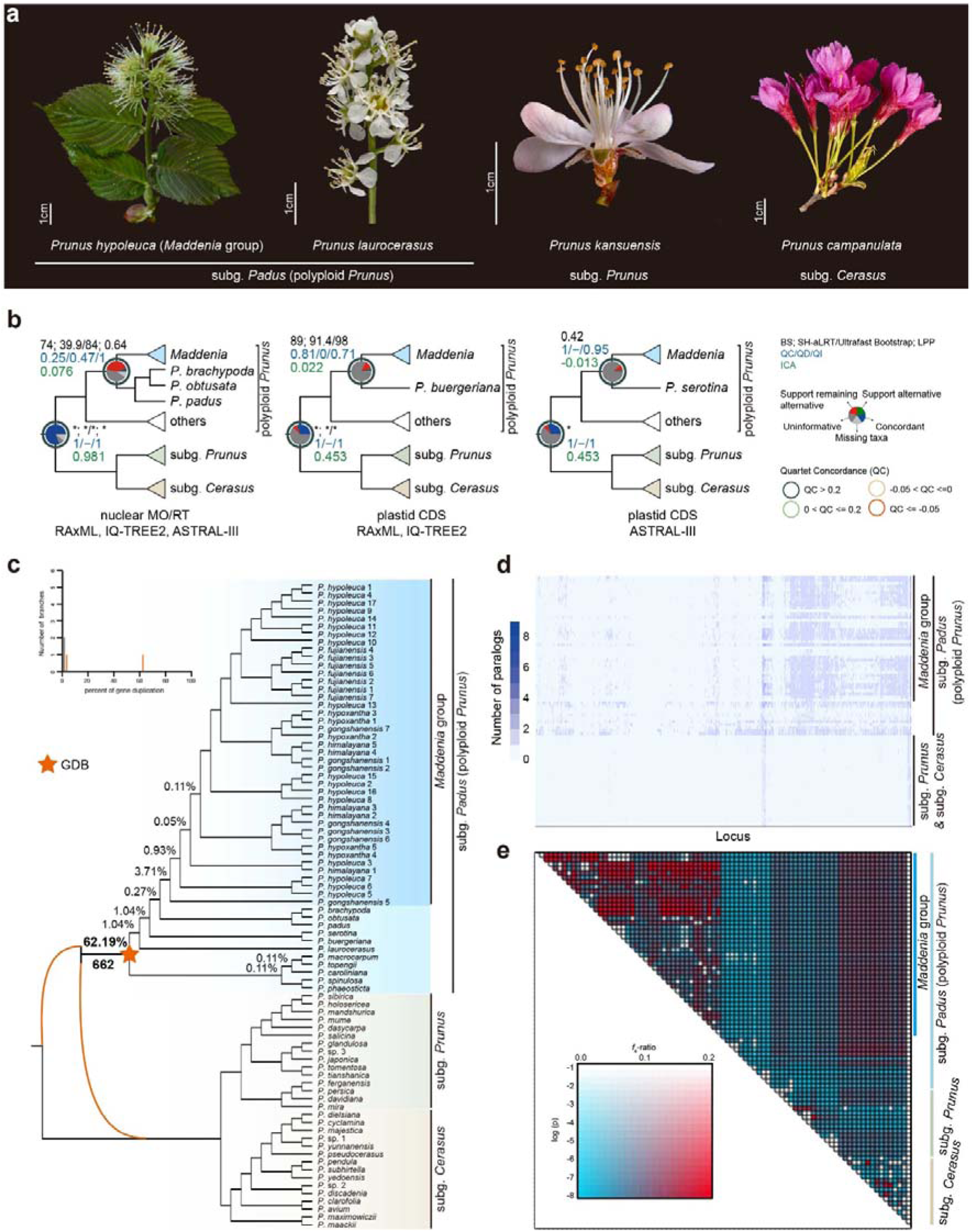
Exploring the contribution of polyploidization in the evolution of polyploid *Prunus* and its close allies. **a** Represented species of three subgenera in *Prunus* s.l., indicating the morphological diversity of inflorescence and flowers. From left to right: *Prunus* subg. *Padus* (*P. hypoleuca* of the *Maddenia* group and *P. laurocerasus*), *P.* subg. *Prunus* (*P. kansuensis*) and *P.* subg. *Cerasus* (*P. campanulata*). Scale bars = 1 cm. **b** Comparative visualization of conflicting topologies from different datasets and inference methods. Phylogenetic supports of the focal nodes from trees are presented next to the branch. The BS values from RAxML analysis (e.g., 74); the SH-aLRT support and UFBoot estimated from IQ-TREE2 (e.g., 39.9/84); the LPP from ASTRAL-LJ (e.g., 0.64) (details refer to Supplementary Figs. 3–11; labeled in black); values for QC/QD/QI estimated from QS analysis (eg., 1/-/1, labeled in blue), and ICA scores (labeled in green). Pie charts on the nodes represent the following data: the proportion of gene trees that support that clade (blue), the proportion that supports the main alternative bipartition (green), the proportion that supports the remaining alternatives (red), the proportion (conflict or support) that have less than 50% BS (dark grey), and the proportion that have missing taxa (light grey). The color of the circle around the pie chart represents the value range of QC, where QC > 0.2 is painted in dark green, 0 < QC ≤ 0.2 is painted in light green, -0.05 < QC ≤ 0 is painted in yellow, and QC ≤ -0.05 is painted in red. Notably, only the values from the MO/RT dataset are shown for MO/RT nuclear trees with the same topology recovered by alternative phylogenetic inference methods. **c** A cladogram of *Prunus* s.l. inferred from ASTRAL-III of MO ortholog trees. The upper left: Histogram of percentages of gene duplication per branch. The percentages above branches denote the proportion of duplicated genes when orthogroups from the homologs are used. Numbers below branches indicate the duplicated gene families of the node (details referring to Supplementary Fig. 29). The orange star and orange curved branches denote the allopolyploid origin of the *P.* subg. *Padus.* **d** Heatmap showing the number of paralog sequences for each gene and each sample recovered by HybPiper. Each row shows a sample, and each column is a gene. The amount of shading in each box corresponds to the number of genes recovered for that sample by the pipeline. **e** Heatmap showing statical support for gene flow between pairs of species inferred from Dsuite package. The shaded scale in boxes represents the estimated *f*_4_-ratio branch value.

The monophyly of the *Maddenia* group was consistently recovered across all six nuclear and three plastid CDS trees with maximal support (SH-aLRT/UFBoot = 100/100; BS = 1; LPP = 1; Figs. 1 and 2b; Supplementary Figs. 3–11). Despite this consensus, the phylogenetic placement of the *Maddenia* group yielded three divergent hypotheses. In the nuclear trees, it was sister to a *P.* subg. *Padus* clade from East-Central Asia (*Prunus brachypoda* + *P. obtusata* + *P. padus*) with moderate support (BS = 74; SH-aLRT/UFBoot = 39.9/84; LPP = 0.64; Fig. 2b; Supplementary Figs. 3–8). In the plastid species tree inferred by ASTRAL-III, it was sister to a New World species (*P. serotina*) with moderate support (LPP = 0.42; Fig. 2b; Supplementary Fig. 9). Alternatively, in the plastid ML trees, it was sister to an East Asian species (*P. buergeriana*) but with strong support (BS = 89; SH-aLRT/UFBoot = 91.4/98; Fig. 2b; Supplementary Figs. 10 and 11). Further conflict analysis, utilizing the *phyparts*, indicated that only 61 out of 765 informative SCN genes were concordant with the ASTRAL species tree (ICA = 0.076; Fig. 2b; Supplementary Figs. 12 and 13), with moderate QS support (0.25/0.47/1; Fig. 2b; Supplementary Fig. 14). The conflict analysis based on the ML tree topology was congruent with the ASTRAL tree, with 61 out of 765 informative SCN genes support with the topology (ICA = 0.115; Supplementary Figs. 18 and 19) and QS support (0.28/0.38/0.98; Supplementary Fig. 20). Conversely, conflict analyses based on the plastid CDS offered an alternate scenario at this node, with only two out of 14 plastid informative trees being concordant to the ML tree (ICA = 0.022; Fig. 2b; Supplementary Figs. 15 and 16) and strong QS support (0.81/0/0.71; Fig. 2b; Supplementary Fig. 17). The conflict analysis based on the species tree topology yielded similar findings to those from the ML tree, with only one of nine plastid informative genes in agreement with the species tree (ICA = -0.013; Fig. 2b; Supplementary Figs. 21 and 22) and full QS support (1/-/0.95; Fig. 2b; Supplementary Fig. 23).

In contrast to the well-defined phylogenetic relationships among the major clades within *Prunus* s.l., the species-level relationships within the *Maddenia* group exhibited topological discordance across both plastid and nuclear datasets, and between inference methods (Fig. 3; Supplementary Figs. 3–11, 24, 25). Moreover, none of the five morphologically distinguished species within the *Maddenia* group were consistently identified as monophyletic across all nine phylogenetic trees. Our conflict analysis from *phyparts* revealed persistent incongruence, as demonstrated by the widespread presence of red pies at all nodes related to the *Maddenia* group, regardless of whether the topology was based on SCN or plastid CDS datasets (Supplementary Figs. 12, 15, 18, 21).

**Figure 3.**
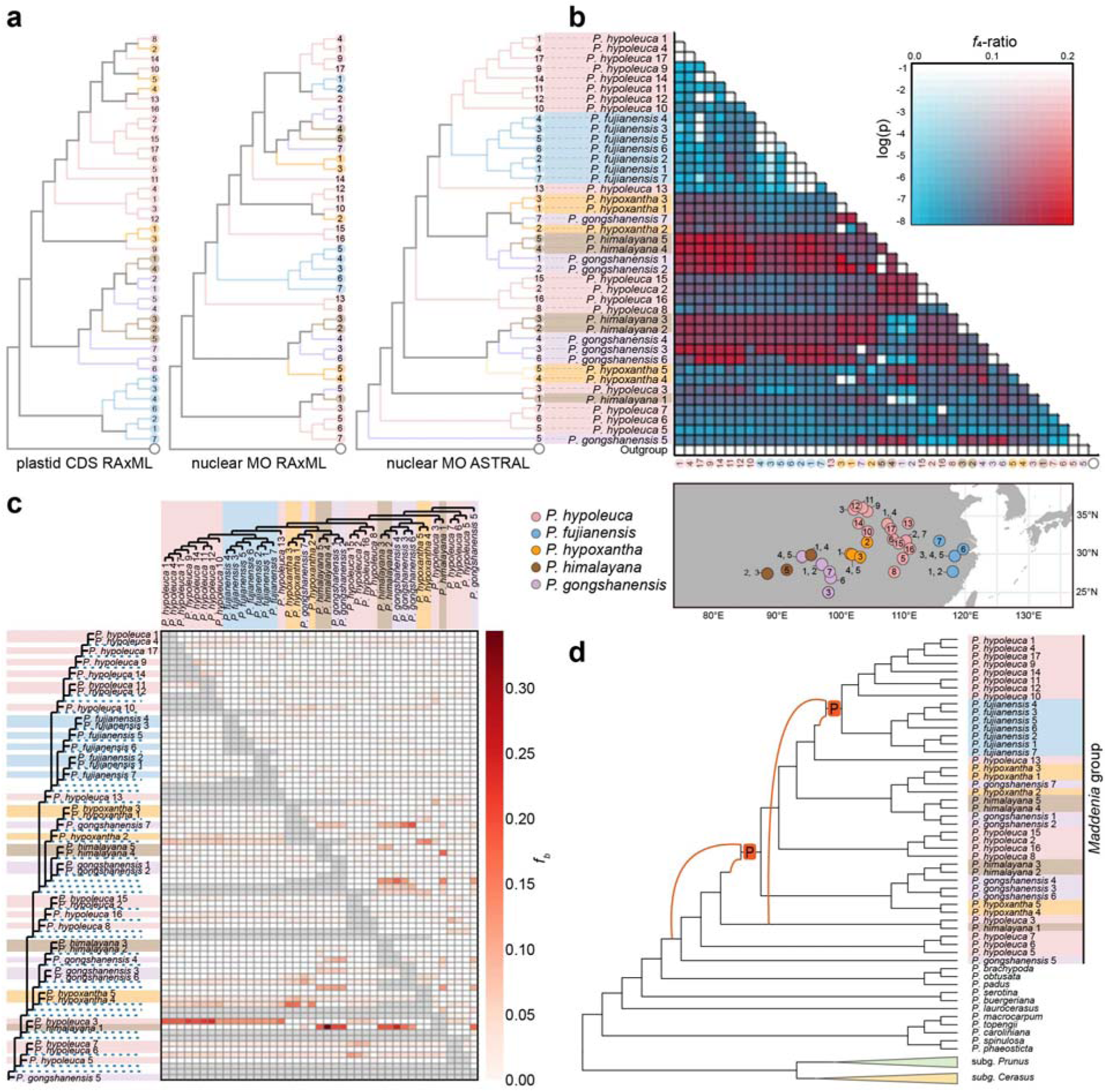
Topological structure conflict and interspecific gene flow of the *Maddenia* group. **a** Topological discordance among the 42 samples, encompassing all five species of the *Maddenia* group, based on various datasets (plastid vs. nuclear) and phylogenetic inference methods (concatenation- and coalescent-based). Three distinct topologies are shown from left to right: a ML tree inferred by RAxML from the plastid CDS dataset, an ML tree inferred by RAxML from the nuclear MO dataset, and a species tree derived by ASTRAL-III from the nuclear MO dataset. The detailed tree information refers to Supplementary Figs. 10, 4, and 3, respectively. **b** Heatmap showing statical support for gene flow between pairs of species inferred from Dsuite package. The shaded scale in boxes represents the estimated *f*_4_-ratio branch value. The map below shows sample collection sites. **c** The matrix shows inferred introgression proportions as estimated from ASTRAL-III species trees in the introgressed species pairs, and then mapped to internal branches using the *f*-branch method. The *f*-branch statistic identifies possible gene flow from the branch of the tree on the y axis to the species or population on the x-axis. **d** Summary of optimal MUL-tree inferred from GRAMPA analyses. MUL-tree from reconciliations of homologs against and ASTRAL tree inferred from MO orthologs including all taxa. Colored curved branches indicate the two possible allopolyploid events within *Maddenia* group (details referring to Supplementary Figs. 34 and 35).

Additionally, a split network for *Prunus* s.l. utilizing the nuclear MO datasets supported the division into three major clades, distinctly separating the polyploid *Prunus* from the remainder of *Prunus* s.l.. Within this framework, *P.* subg. *Prunus* and *P.* subg. *Cerasus* exhibited close phylogenetic relationships. Species within the *Maddenia* group were markedly divergent from other species in the polyploid *Prunus*, underscoring substantial phylogenetic differentiation (Fig. 1a).

### Negligible role of Incomplete Lineage Sorting in polyploid *Prunus* evolution

We employed two analytical methods to investigate the role of Incomplete Lineage Sorting (ILS) in the evolutionary history of polyploid *Prunus*. The nuclear Coalescent Simulation (CoSi) analysis, which differentiates between empirical and simulated gene-to-gene distance distributions, indicated that ILS cannot account for the observed conflicts between gene trees and the species tree (Fig. 1c). Specifically, the node encompassing polyploid *Prunus* and its sister exhibited a theta value of 0.0076 in the Mutation Calculation Based on Coalescent (MuCCo) analysis, suggesting a minimal influence of ILS on the origin of polyploid *Prunus*. Similarly, theta values for the crown of polyploid *Prunus* (theta = 0.0099), the stem of the *Maddenia* group (theta = 0.035), and the crown of the *Maddenia* group (theta = 0.0049) were also notably low (Fig. 1b; Supplementary Fig. 26), further supporting the limited impact of ILS. In summary, both the CoSi (Fig. 1c) and MuCCo analysis (theta value < 0.1; Fig. 1b; Supplementary Fig. 26) corroborate the conclusion that ILS has negligible influence on the evolutionary dynamics of polyploid *Prunus*.

### The dominant role of allopolyploidization in the origin and dynamics of polyploid *Prunus*

According to the literature, the base chromosome number for members of *Prunus* s.l. is *x* = 8^45^, with variation in ploidy levels among these species. Most species within *P.* subg. *Prunus* and *P.* subg. *Cerasus* have chromosome counts of 2n = 2*x* = 16, whereas the polyploid *Prunus* typically exhibit higher chromosome levels (e.g., 2n = 4*x* = 32 for most species, 2n = 22*x* = 176 for *P. laurocerasus*)^46^. Our flow cytometric analysis revealed diverse chromosome counts in the *Maddenia* group, identifying 2n = 4*x* = 32 in *P. fujianensis* 1, *P. himalayana* 4, *P. hypoleuca* 2, and *P. hypoxantha* 1, and 2n = 8*x* =64 in *P. gongshanensis* 7 (Supplementary Fig. 27).

Analysis of paralogs from homologous gene trees revealed that polyploid *Prunus* exhibits a significantly higher number of paralogs than other species within *Prunus* s.l. (Fig. 2d; Supplementary Fig. 28). This increased number of paralogs suggests potential gene duplication (GD) events throughout the evolutionary history of polyploid *Prunus*. We subsequently mapped these homologous gene trees onto the MO-based species tree of *Prunus* s.l., estimating the proportion of GD at each node (Fig. 2c). Our findings indicated that one node within *Prunus* s.l. showed an elevated proportion of GD compared to others, marked with an orange star (Fig. 2c). The node corresponding to the most recent common ancestor (MRCA) of polyploid *Prunus* presented the highest duplication proportion, with 62.19% of genes indicating support for the duplication event (Fig. 2c).

As an alternative approach, we used the Tree2GD to analyze 229,705 genes from polyploid *Prunus* genomes using the DGS dataset. Reconciliation of 36,488 gene trees with the species tree identified 662 GDs in the clade of *P.* subg. *Padus* (Fig. 2c; Supplementary Fig. 29). Among these GDs, 63% retained both duplicates (ABAB type), providing further evidence for WGDs (Fig. 2c; Supplementary Fig. 29).

We employed GRAMPA to examine the mode of WGD event suggested by our GD and Tree2GD analysis. The results confirmed that the WGD event associated with polyploid *Prunus* was of an allopolyploid origin, with the optimal multi-labeled trees (MUL-trees) achieving the best score (i.e., Lowest Reconciliation Score (LRS) of 474,967). This polyploid *Prunus* likely originated from an ancient hybridization between the MRCA of a combined clade (*P.* subg. *Prunus* + *P.* subg.*Cerasus*) and the MRCA of *Prunus* s.l. (Fig. 2c; Supplementary Fig. 30).

Further exploration of introgression in *Prunus* s.l. revealed two distinct evolutionary patterns (Supplementary Fig. 31). The polyploid *Prunus* exhibited significant gene flow with a combined clade, specifically *P.* subg. *Prunus* and *P.* subg. *Cerasus*, with notable interactions particularly within the *P.* subg. *Cerasus* (Fig. 2e). However, within the focused *Maddenia* clade, while significant genetic introgression was observed, there was virtually no detectable gene flow between the *Maddenia* clade and any other clades within the polyploid *Prunus* (Fig. 2e; Supplementary Fig. 31).

### Hybridization and polyploidization in the *Maddenia* group

The tree topologies derived from plastid CDS and SCN datasets for the *Maddenia* group showed considerable discordance. In the plastid CDS and whole plastome trees, the *Maddenia* group split into three major clades, roughly corresponding to their geographical distribution (Fig. 3a; Supplementary Figs. 9–11, 24, 25). In contrast, the six SCN trees depicted significantly conflicting phylogenetic relationships with varying levels of support (strong, moderate, or low) across the *Maddenia* group nodes (Fig. 1b; Supplementary Figs. 3–8), despite using up to 1,622 (MO dataset) and 2,268 (RT dataset) SCN genes. None of the five recognized species within these datasets consistently formed monophyletic groups, except for *Prunus fujianensis*, which was monophyletic in species trees from the nuclear MO and RT datasets (Fig. 3a; Supplementary Figs. 3 and 6). Specifically, *P. fujianensis* appeared polyphyletic in the SCN gene trees but was monophyletic in the CDS and SCN ASTRAL trees, where it was nested within *P. hypoleuca*. Meanwhile, the other four *Maddenia* species—*P. himalayana*, *P. gongshanensis*, *P. hypoleuca*, and *P. hypoxantha*—were all polyphyletic in the SCN phylogeny (Fig. 3a; Supplementary Figs. 3–8).

Our introgression analysis using Dsuite identified two major clades that extensively introgressed with other samples or clades within the *Maddenia* group (Fig. 3b; Supplementary Fig. 32). One clade consisted of four samples (two each from *P. gongshanensis* 1 & 2 and *P. himalayana* 4 & 5), and another included five samples (two from *P. himalayana* 2 & 3 and three from *P. gongshanensis* 3, 4, & 6). Although these two clades overlapped geographically in the Himalayas and Hengduan Mountains, weak or no introgression was detected between them. In contrast, gene flow has been observed between the Himalaya-Hengduan region and East and Central China.

The *f*-branch statistic affirmed the extensive reticulate evolution at both deep and shallow phylogenetic levels (Fig. 3c; Supplementary Fig. 33). The nuclear ASTRAL tree showed clearer signals of gene flow among the basal lineage, particularly at higher *f*-branch values. In the *Maddenia* group, we observed widespread interspecific and intraspecific gene flow. Notably, gene flow between *P. himalayana* and *P. gongshanensis*, as well as between *P. hypoleuca* and *P. fujianensis*, indicated interspecific hybridization. Additionally, our analysis detected intraspecific hybridization within samples from *P. fujianensis*, *P. gongshanensis*, *P. himalayana*, *P. hypoleuca*, and *P. hypoxantha*, highlighting complex hybridization patterns in this group.

To explore variations in ploidy and genome size within the *Maddenia* group, we performed flow cytometric analysis on several selected samples. Our findings revealed that the C value of *P. hypoleuca* is approximately 519 Mb, using rice as the reference. In contrast, the C value for *P. gongshanensis* is about 1,282 Mb, using maize as the reference, nearly twice that of other species examined (Supplementary Fig. 27).

To explore the drivers of the differentiation in the *Maddenia* group, we followed the methods proposed by Morales-Briones et al.^47^ to disentangle potential polyploidy events using GRAMPA^48^. Our results revealed two allopolyploid events within the *Maddenia* group (Fig. 3d; Supplementary Figs. 34 and 35). The monophyletic group *P. fujianensis* and its sister clade, which included some samples of *P. hypoleuca*, were the descendent lineage of the lineage “*P. himalayana* 1 and *P. hypoleuca* 3” and the *P. hypoleuca* 13 (Fig. 3d; Supplementary Fig. 34). The clade consisting of almost all samples, except *P. himalayana* 1, *P. hypoleuca* 3, 5, 6, & 7, and P. *gongshanensis* 5, was the descendent lineage of the lineage “*P. himalayana* 1 and *P. hypoleuca* 3” and the MRCA of the *Maddenia* group (Fig. 3d; Supplementary Fig. 35).

### Diversification and biogeographic analyses

The historical biogeographic analyses of the polyploid *Prunus* and the *Maddenia* group were conducted using both SCN genes and plastid CDS datasets. Our findings indicate that *Prunus* s.l. originated approximately 74.97 million years ago (Mya), with a 95% highest posterior density (HPD) interval of 83.98–64.47 Mya, according to the nuclear MO dataset (Supplementary Figs. 36 and 37). Similarly, estimates based on the plastid CDS dataset suggest an origin at 71.62 Mya, with a 95% HPD of 86.84–52.36 Mya (Supplementary Figs. 38 and 39). These estimated times were subsequently used as secondary calibration nodes in the diversification analysis of polyploid *Prunus*.

In the nuclear MO dataset, polyploid *Prunus* originated in East Asia in the Late Eocene, approximately 35.25 Mya (95% HPD: 37.07–33.81 Mya), and subsequently dispersed westward to West Asia and eastward to the New World around 34.02 Mya (95% HPD: 36.05–32.33 Mya) (Fig. 4a, c; Supplementary Figs. 40–42). We inferred that the *Maddenia* group originated in Central China in the Early Oligocene, around 31.57 Mya (95% HPD: 33.77–29.56 Mya), and diversified in the Late Oligocene, approximately 26.18 Mya (95% HPD: 28.90–23.54 Mya; Supplementary Figs. 40, 41, 43). The MRCA of the *Maddenia* group was likely distributed in Central China, subsequently dispersing to Southeast and Southwest China through multiple dispersal and vicariance events (Supplementary Fig. 43). The divergence times estimated for polyploid *Prunus* using the plastid CDS dataset were similar to those inferred from the SCN genes dataset but slightly younger for the *Maddenia* group. The ancestral area for this lineage was estimated to cover an expansive region that includes central and southwest China (Fig. 4a, b; Supplementary Figs. 44–47).

**Figure 4.**
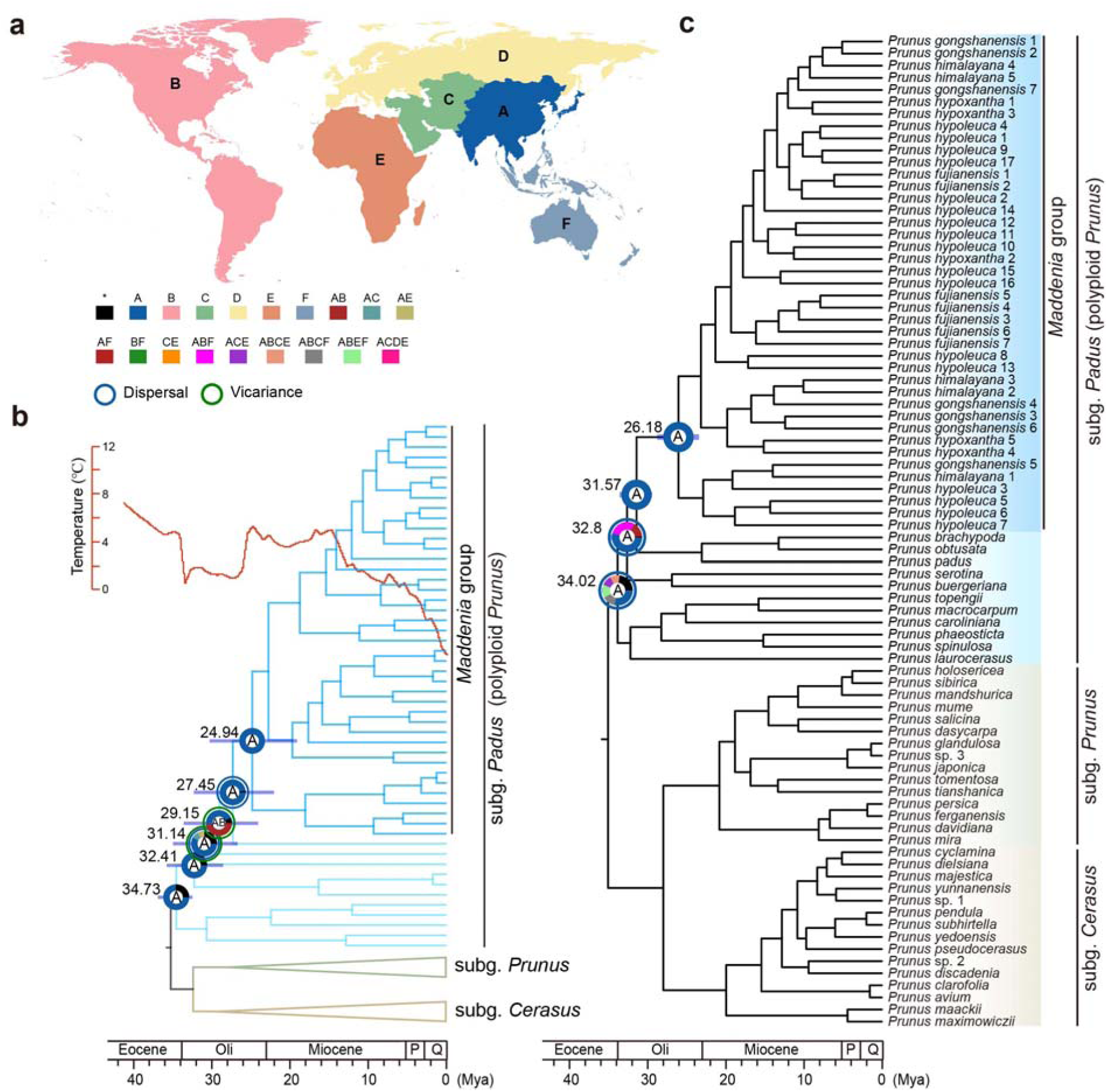
Divergence time estimation and geographical range evolution of the polyploid *Prunus*. **a** Inset map outlines the six distribution areas used for geographical analysis (details referring to Supplementary Figs. 33–38). (A) East Asia, (B) North & South America, (C) West Asia, (D) Europe, (E) Africa, and (F) Australasia. **b** Dated chronogram of the *Prunus* inferred from PAML based on the plastid CDS dataset. Focal nodes feature 95% confidence of estimated divergence times and ancestral geographical ranges. The red line represents the global temperature changes from Westerhold et al.^125^. **c** Dated chronogram of the *Prunus* s.l. inferred from PAML based on the nuclear MO dataset. Focal nodes feature 95% confidence of estimated divergence times and ancestral geographical ranges.

The EBD analyses based on the SCN and plastid CDS datasets revealed distinct diversification patterns. Our nuclear EBD analysis showed a pattern of increased diversification, characterized by a speciation rate of approximately 0.3, from 20 to 7 Mya, followed by a sharp decline after the Late Miocene (around 7 Mya). Then, the relative extinction rate began to increase after 7 Mya (Fig. 5a). In contrast, the plastid CDS-based EBD analysis indicated a distinct pattern, with a high net diversification rate (> 0.2) from about 10 to 4 Mya. Since 20 Mya, the speciation rate has shown a consistent trend, with a significant rise starting around 10 Mya, paralleling the trend in the net diversification rate of the lineage. The extinction rate gradually decreased, and the relative extinction rate declined but experienced a sudden rise around 2 Mya (Fig. 5b).

**Figure 5.**
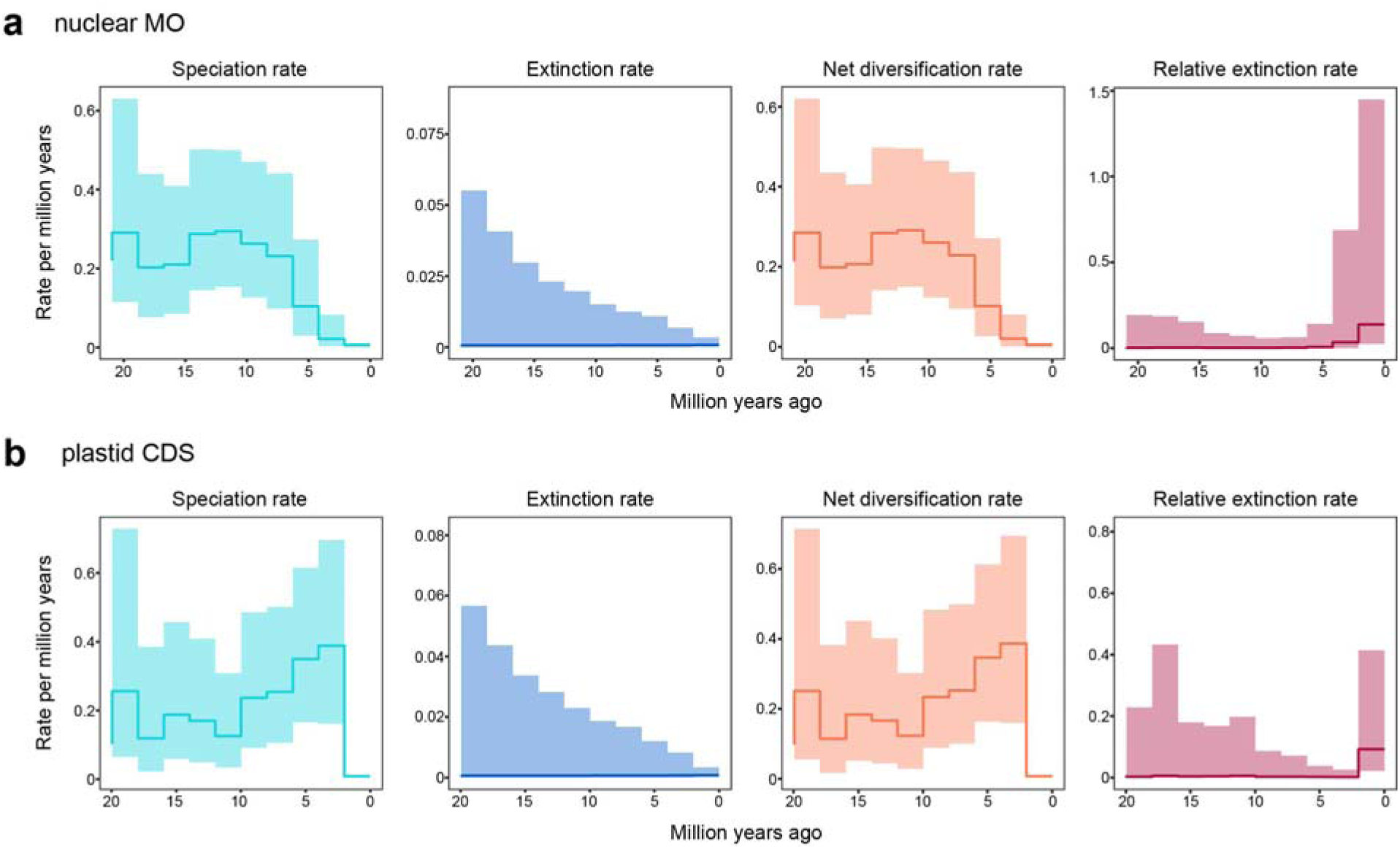
Speciation rates, extinction rates, net diversification rates and changes in speciation rates among major lineages within the *Maddenia* group over time base on Revbayes analysis. **a** Using nuclear MO dataset. **b** Using plastid CDS dataset. Shaded regions show the 90% credible intervals for the rate.

## Discussion

### Allopolyploidization drives the origin of the polyploid *Prunus*

The monophyly of polyploid *Prunus* has been debated, with support for both monophyly^16,19,49^ and paraphyly^14,50–52^. Our study strongly supports monophyly across nine phylogenies (six nuclear and three plastid CDS ones; Figs. 1 and 2b, Supplementary Figs. 3–11), consistent with recent RNA-Seq-^16^ and restriction site-associated DNA sequencing (RAD-Seq)-based^19^ studies. Earlier conflicting results likely stem from limited markers and sampling issues, compounded by introgression events between polyploid *Prunus* and the other two subgenera (Supplementary Fig. 31)^43^.

We used polyploid *Prunus* to demonstrate a progressive strategy for analyzing DGS data in the presence of genome doubling. Our findings indicate allopolyploidization, rather than ILS, as the main driver in the origin and dynamics of polyploid *Prunus*. Our conflict analyses showed that nuclear SCN genes (1,317 of 1,321; ICA = 0.981; Supplementary Figs. 12 and 13) and plastid CDS (23 of 29; ICA = 0.453; Supplementary Figs. 15 and 16) generally support a concordant topology (Fig. 2b). However, successive WGD and polyploidization analyses indicate an alternative scenario: polyploid *Prunus* likely originated through a WGD event, with 62.19% of duplicated genes, followed by an allopolyploidy event (Fig. 2c, d). Previous studies using multiple copies of *At103* genes also support this allopolyploid origin^18^. Further introgression analysis revealed extensive gene flow between the polyploid *Prunus* clade and a combined clade of *P.* subg. *Prunus* and *P.* subg. *Cerasus*, with greater introgression from *P.* subg. *Cerasus* (Supplementary Fig. 31). These findings suggest that the MRCA of *Prunus* s.l. likely acted as the paternal parent, while the MRCA of *P.* subg. *Prunus* and *P.* subg. *Cerasus* served as the maternal parent. These MRCAs may represent extinct or unsampled “ghost species” that transferred genetic material to current lineages^53,54^.

We propose that these ancestral groups hybridized in East Asia during the Late Eocene, with subsequent WGD facilitating their adaptation to climate changes, including the cooling trend of the Eocene-Oligocene transition (EOT)^55^. Conflict analyses of SCN and plastid gene trees consistently show uneven genetic contributions, likely due to recurrent backcrossing with the maternal parent. The orogeny of the Himalayas and Hengduan Mountains likely shaped the East Asian monsoon and Yangtze River system^56^, creating new environmental niches that promoted frequent backcrossing. This upheaval may also have led to the extinction of the paternal parent, a scenario similar to that observed in the origin of *Weniomeles*^42^.

### Rampant introgression drives the diversification of the *Maddenia* group

Using the Step-by-Step Exclusion (SSE)^43^ approach, we evaluated the impact of reticulate evolution on the diversification of the *Maddenia* group. Divergence in the distance distribution from CoSi analysis suggests that ILS is not the primary cause of phylogenetic conflict (Fig. 1c). Low theta values (theta < 0.1) throughout the nodes from MuCCo analysis further discount ILS as a factor in origin and diversification of *Maddenia* (Fig. 1b; Supplementary Fig. 26). We propose that hybridization-driven genetic introgression significantly influenced its diversification, consistent with previous findings that widespread hybridization eroded reproductive barriers among species^19^. Our analyses detected extensive reticulate evolution at multiple phylogenetic levels, with nearly all species exhibiting gene flow with those from Central China (Fig. 3b, Supplementary Fig. 32). This supports the hypothesis that the MRCA of *Maddenia* group originated in Central China and underwent rapid radiation during the Miocene (Fig. 4; Supplementary Figs. 40, 41, 43–45, 47). Biogeographic analyses reveal multiple dispersal and vicariance events, spreading the lineage southwest and southeast over time (Supplementary Figs. 43 and 47). The *f*-branch results showed minimal differences in gene flow among branches (Fig. 3c; Supplementary Fig. 33), suggesting uniform genetic exchange ratios^57^. We infer that species within the *Maddenia* group likely followed a similar evolutionary path, where hybridization- and polyploidization-driven introgression facilitated rapid radiation during the Miocene. GRAMPA analyses showed that nearly all samples participated in polyploidy events as either parents or offspring (Fig. 3d; Supplementary Figs. 34 and 35), underscoring the significant role of allopolyploidization in its diversification. We also observed ploidy variations within the group, ranging from tetraploid to octoploid (Supplementary Fig. 27). Similar allopolyploidization events, often involving changes in ploidy or chromosome numbers, have been documented in other taxa, including Didymocarpinae (Gesneriaceae)^58^, *Rosa* (Rosaceae)^59^, and Campanuleae (Campanulaceae)^6^, highlighting allopolyploidization as a key driver of plant diversification.

The Himalayan uplift has greatly impacted species diversity across various lineages. Alpine bamboos in the Hengduan Mountains^60^ and *Rhododendron* (Ericaceae)^61^ have shown increased speciation linked to mountain uplift and Miocene tectonic activity. Our biogeographic analysis suggests that the *Maddenia* group originated in Central China and diversified during the Late Oligocene warming (Fig. 4; Supplementary Figs. 40, 41, 43–45, 47). From the Late Eocene to Miocene, the uplift of the Himalayas, driven by the Indian-Asian plate collision^62,63^, likely promoted rapid radiation in the *Maddenia* group, evidenced by the increased speciation and net diversification rates from 20 to 7 Mya (Fig. 5). Around 11 Mya in the Late Miocene, intensified monsoon circulation and erosion counteracted further uplift, slowing speciation rates of the *Maddenia* group (Fig. 5). Orogenesis-induced reticulate evolution have led to widespread interspecific gene introgression.

### Whole-genome duplication analysis in deep genome skimming data

Polyploidy and WGD are widespread in plant evolution^2,25,64–66^, with all modern angiosperms having experienced at least one WGD^25^. While chromosome ploidy levels have been used to detect WGDs^67^, estimating ancient events requires genomic data due to gene loss, mutations, and diploidization over time^68–70^. Advances in sequencing and bioinformatics have streamlined WGD analysis but also introduced new challenges.

Gene tree/species tree reconciliation accurately identifies gene duplications across lineages. It maps each node in the homolog gene tree to the respective node in the species tree, identifying duplications if nodes align with their direct descendants^47,48,71^. Nodes with many duplications suggest a polyploidy origin^72^. However, this approach leverages robust phylogenetic relationships and genomic data to count duplicated genes precisely. Gene tree/species tree reconciliation is commonly used to investigate WGDs in RNA-Seq and Hyb-Seq datasets. However, its application to DGS datasets is challenged by the need for sufficient paralogs, dense taxon sampling, and robust phylogeny.

Genome-wide duplication events typically generate thousands of paralogous genes^24,64^, often having various evolutionary changes. These changes can include partial degradation from accumulated mutations, reducing their functional capacity to that of their single-copy ancestral counterparts. Alternatively, duplicated genes may become nonfunctional due to degenerative mutations or be ultimately lost^24,68^. Genomic-scale datasets like DGS offer a unique advantage by capturing a wider array of paralogs, including silent genes that are often undetectable in transcriptomic data. This comprehensive approach minimizes inaccuracies and biases from missing data, enabling a more complete evaluation of all genes.

Dense taxon sampling across major clades is essential for accurately detecting WGDs. The DGS method performs well at utilizing degraded or ancient DNA from herbarium/museum specimens, enabling the inclusion of rare and extinct species in phylogenetic analyses, thus bridging sampling gaps^73–75^. In contrast, RNA-Seq relies on fresh or flash-frozen materials, limiting its ability to achieve broad taxon sampling^36,76^.

Analyzing WGDs requires a robust phylogenetic framework, especially when using tree-based methods. Stiller et al.^77^ highlighted that the number of loci impacts the inferred tree more than taxon sampling. While RNA-Seq and Hyb-Seq are commonly used in phylogenetic studies^31,41,78–80^, assembling orthologs can be challenging due to data variations resulting from different tissues, time, individuals, growth stages, and environmental conditions^76^. Furthermore, generating thousands of orthologous loci using these methods is often prohibitively expensive, even for well-funded labs. In contrast, the DGS method is more flexible, cost-effective, and avoids complex experimental procedures compared to RNA-Seq and Hyb-Seq^32^. By increasing sequencing depth, DGS efficiently captures large numbers of orthologous loci, proving its effectiveness as a reliable tool for phylogenetic research^6,8,32,42,43^.

Numerous bioinformatics tools exist for RNA-Seq and Hyb-Seq data^2,30,47,71,81^, but software designed explicitly for whole genomic data has lagged behind genomic sequencing advances. We assessed existing tools and adapted scripts to improve compatibility with whole genomic DGS data. Our strategy investigates allopolyploidization-driven WGDs in polyploid *Prunus*, a lineage with frequent WGDs^16–18^, through three steps: counting gene duplications, mapping them onto the species tree, and distinguishing between allo- and auto-polyploidy events (Fig. 6). We conducted parallel analyses to detect potential WGDs across the phylogeny, utilizing deeply sequenced DGS data to identify paralogs and applying gene tree/species tree reconciliation^47^ to determine WGD events. Simultaneously, we employed DGS-Tree2GD scripts as an alternative approach. Our results highlight the capability of DGS data to assemble numerous nuclear genes, enabling comprehensive analysis of gene duplication bursts (GDB). Clustering of gene duplications at each node of the species tree provides strong evidence of potential polyploidization or WGD events^72^. GRAMPA analysis further differentiates between autopolyploidy and allopolyploidy, two types of WGD events^48^. By integrating fossil records and species distributions, our biogeographic analyses confirm the timing and locations of these polyploidy events^8,42^, identifying polyploid lineages and their parental contributors. This workflow significantly enhances the efficiency and utility of DGS data (Fig. 6).

**Figure 6.**
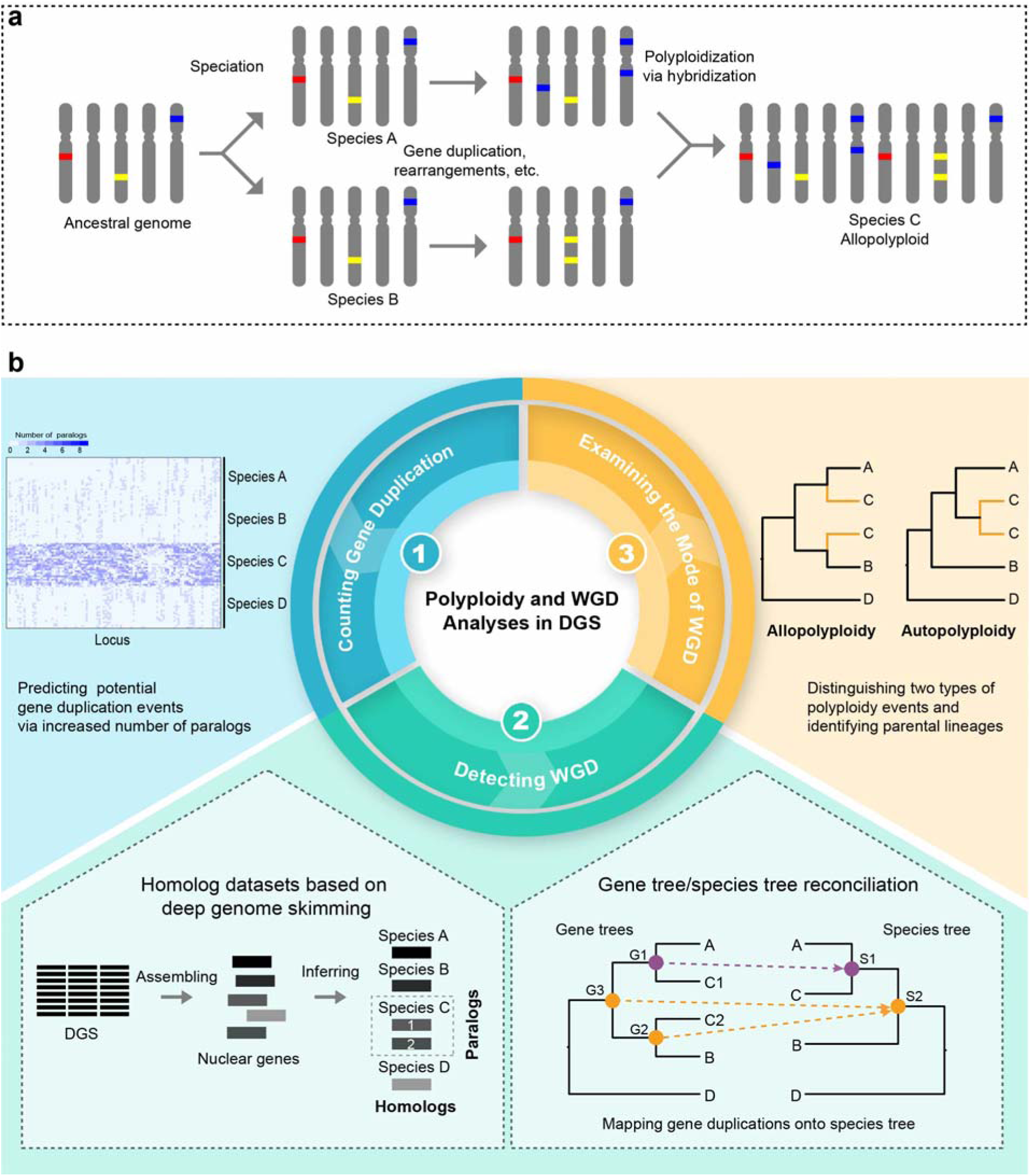
Workflow of progressive analyses for unraveling allopolyploidization-driven genome duplications from Deep Genome Skimming (DGS) Data. **a** Figure illustrates the speciation of the allopolyploid species C through 1074 genome duplication and introgression between species A and B (Adapted from Fig. 2 in Glover et al.^126^). **b** A three-step procedure for analyzing WGD events: **1)** Paralog analysis to count gene duplications, highlighting a significant increase in paralogs in species C compared to species A and B, indicating a potential WGD event. **2)** Detecting WGD using two complementary methods: inferring a robust species tree from the nuclear dataset and mapping gene trees to this species tree for identifying gene duplication events. The left part of the illustration details the process for homolog inference and distinguishing orthologs from homologs from DGS data, while the right part outlines the strategy for mapping duplicated species onto corresponding nodes in the species tree. For instance, Node G1, including tips A and C1, maps to Node S1 in the species tree, while Nodes G2 and G3, containing tips A/C1/C2/B and C2/B respectively, and the MRCA of species A, B, and C, defined as Node S2, are mapped to Node S2, which is also identified as a duplication node. **3)** Examining the mode of WGD: for lineages with potential WGD events, GRAMPA analysis is performed to determine whether they involve allopolyploidy or autopolyploidy, with further confirmation of maternal and paternal parents in cases of allopolyploidy through comparison of nuclear and plastid phylogenies.

## Methods

### Taxon sampling, DNA extraction, and sequencing

This investigation examines the hypothesis of an allopolyploid origin of the polyploid *Prunus*, initially posited by Zhao et al.^18^. Building on a comprehensive phylogenetic framework^17^ for the genus across diverse geographical regions, we follow this backbone and incorporate all major clades for the present study. We particularly focus on the *Maddenia* group, where our taxon sampling was extensive within five focal species. This included seven individuals for *Prunus fujianensis* and *P. gongshanensis*, five for *P. himalayana* and *P. hypoxantha*, and 17 for *P. hypoleuca*, totaling 41 samples. These samples comprehensively represent the broad phylogenetic and geographic diversity observed within the *Maddenia* group^19^. To circumvent the limitation of fossil records in *Prunus* s.l., our sampling strategy was broadened to include 30 more species within the subfamily Amygdaloideae, and we designated *Rosa rugosa* from the subfamily Rosoideae as the outgroup based on the nuclear topology^29^. Consequently, this study assembled a collection of 114 samples, comprising 89 datasets from DGS and 25 from RNA-Seq, thereby ensuring a rich dataset for elucidating allopolyploid evolution in *Prunus* s.l. (Supplementary Table 1).

Genomic DNAs were extracted from both silica-gel dried leaves and herbarium specimens utilizing a modified CTAB protocol^82,83^ within the laboratory of the Institute of Botany, Chinese Academy of Sciences (IBCAS). After a stringent quality control process, DNA libraries were prepared using the NEBNext^®^ Ultra^™^ II DNA Library Prep Kit and then sequenced on the Illumina MiSeq platform (Illumina, San Diego, CA, USA) at the Kunming Institute of Botany, Chinese Academy of Sciences (KUN) or BGISEQ-500 platform (BGI, Shenzhen) of Frasergen (Wuhan, China) for Next-Generation Sequencing (NGS). All resultant raw data have been archived in the National Center for Biotechnology Information (NCBI) Sequence Read Archive (SRA) under the designated BioProject (PRJNA1031385), while the corresponding voucher specimens have been deposited in the herbarium of Northwest A&F University (WUK) and the China National Herbarium (PE). A comprehensive catalog of all 114 samples is in Supplementary Table 1.

### Plastome assembly, annotation, and alignment

The raw data were initially processed to trim low-quality reads and remove adapters using Trimmomatic v. 0.39^84^. The resulting clean reads were then quality-checked with FastQC v. 0.12.1, available at [https://www.bioinformatics.babraham.ac.uk/projects/fastqc/]. Following quality assessment, the plastid genomes were assembled using GetOrganelle v. 1.7.7.0^85^ and subsequently annotated using the Plastid Genome Annotator (PGA)^86^. All plastomes assembled during this study were submitted to GenBank (see the accession numbers in Supplementary Table 1).

In this study, we employed two alternative strategies for extracting plastid coding sequences (plastid CDS) across different data types to generate the “plastid CDS” dataset. Geneious Prime v. 2023.2^87^ was utilized to extract the plastid CDS from the assembled complete plastomes. Concurrently, we employed HybPiper v. 2.1.6^88^ to assemble the plastid CDS from RNA-seq data, with the closely related CDSs downloaded from GenBank used as references. The plastid CDSs were aligned using MAFFT v. 7.520^89^ and trimmed using trimAl v. 1.4.1^90^, in which poorly aligned regions were removed. Then those sequences were concatenated with AMAS v. 1.0^91^. Given the reduced site variation within the *Maddenia* group, we generated a “*Maddenia*-plastome” dataset specifically to infer the maternal phylogeny of *Maddenia*. All the aligned matrixes were deposited in the Dryad [DOI: https://datadryad.org/stash/share/zrSRWvefAjRapx7TElIyJAh0uKcmofE41va3HnVH4o8].

### Single-copy nuclear loci development and assembly

We isolated the coding sequences of three genomes within *Prunus* s.l., i.e., *P. dulcis* (accession number: GCA902201215.1), *P. armeniaca* (accession number: GCA903112645.1), and *P. persica* (accession number: GCA000346465.2). This analysis was conducted utilizing MarkerMiner v. 1.0^92^, which was used to select SCN loci that could serve as phylogenetic markers, with “Fvesca” (*Fragaria vesca*) as the reference. The parameter “minTranscriptLen” was set to a minimum length of 600 bp with the remaining parameters retained at their default values, balancing the need for specificity in our search criteria with the broad-based, standardized approaches inherent to the software. The reference genes available on Dryad [DOI: https://datadryad.org/stash/share/zrSRWvefAjRapx7TElIyJAh0uKcmofE41va3HnVH4o8].

We implemented the HybPiper v. 2.1.6 pipeline^88^ to assemble SCN loci from the clean reads. The specific assignment of reads to their corresponding target loci was facilitated by BWA v. 0.7.17^93^. Following this, SPAdes v. 3.15.5^94^ was utilized to execute *de novo* assembly of the reads individually. This was done with a coverage cutoff value set to 5 to ensure that only reads with sufficient depth were assembled, thus enhancing the reliability of the assembled sequences. We used commands to statistics on sequence length (Supplementary Table 2) and facilitate the visualization of the assembly results with a heatmap (Supplementary Fig. 1). We also discerned and annotated putative paralogous genes—those genes that arise from duplication events within the genome—within each sample. Identifying such paralogs is essential to avoid potential confounding factors in subsequent analyses. The resulting sequences were subsequently used for orthology inference.

### Orthology inference and dataset generation

The identification of potential paralogs is critical for ensuring the accuracy of phylogenetic inferences. In this study, we implemented an automated orthology inference approach for nuclear genes assembled from DGS reads to generate two SCN datasets, “Monophyletic Outgroup” (MO) dataset and “RooTed ingroup” (RT) dataset with *Rosa rugosa* designated as the outgroup. This approach initially conceptualized for RNA-Seq data analysis by Yang & Smith^71^, underwent subsequent modifications for adapting the target enrichment datasets, such as Hyb-Seq^4^. Our PhyloAI team has executed further refinements to these scripts, ensuring their compatibility with DGS datasets. This enhancement has been substantiated through its successful application in recent studies, such as the taxonomic research on *Malus*^8^ and *Pyrus*^42,43^. For detailed descriptions of these procedures, refer to the Supplementary Methods.

### Integrative phylogenetic strategies: concatenation- and coalescent-based analyses

In this study, we used concatenation- and coalescent-based methods across plastid CDS and two SCN loci datasets for accurate phylogenetic inference. The selection of optimal partitioning schemes and evolutionary models was conducted using PartitionFinder2^95^, guided by the corrected Akaike information criterion model (AICc), and incorporated linked branch lengths for model testing. To accommodate the varying complexities and sizes of the datasets, we implemented two distinct search strategies: the ‘greedy’ algorithm^96^ for its efficiency in larger datasets and ‘rcluster’^97^ for its capability to manage computational demands effectively. The determined partitioning schemes and evolutionary models were applied for the subsequent phylogenetic inference.

ML trees were inferred utilizing IQ-TREE2 v. 2.2.2.7^98^, engaging 1,000 replicates each for ultrafast bootstrap support assessment^99^ and the SH-aLRT test^100^ to evaluate tree topology reliability. Furthermore, RAxML v. 8.2.13^101^ was utilized as a supplementary tool for ML tree inference, employing the best-fit substitution model. This was accompanied by the generation of 200 bootstrap replicates for the SCN datasets and 100 replicates for both the “plastid CDS” and “*Maddenia*-plastome” datasets.

Coalescent-based species trees were inferred using ASTRAL-III^102^, with input gene trees generated from the plastid CDS and SCN loci datasets, in order to infer species relationships while accommodating gene tree discordance. This integrative strategy ensures capturing maternal lineage via plastid data and the patterns of biparental inheritance through SCN sequences. All the resultant trees are available through the Dryad Digital Repository [DOI: https://datadryad.org/stash/share/zrSRWvefAjRapx7TElIyJAh0uKcmofE41va3HnVH4o8].

### Inference of global split networks

Increasing evidence suggests that web-like patterns of evolution instead of simple tree-like ones can explain how different groups are related, especially when complex processes like hybridization and polyploidy play a role. Given the frequent observation of reticulate signals within *Prunus* s.l., particularly among polyploid groups, we utilized SplitsTree v. 4.19.2^103^ to estimate a split network from aligned SCN sequences derived from the MO dataset.

### Detecting and visualizing nuclear gene tree discordance

To elucidate the discrepancies between gene trees and species trees, we utilized *phyparts* v. 0.0.1^104^ to perform an in-depth analysis of conflicting and concordant bipartitions. This analysis facilitated the calculation of ICA scores, thereby providing a quantitative measure of the extent of discordance at each phylogenetic node. These analytical outcomes were visualized by implementing the script “phypartspiecharts_missing_uninformative.py”, accessible via (https://bitbucket.org/dfmoralesb/target_enrichment_orthology/src/master/phypartspiecharts_missing_uninformative.py).

Furthermore, QS analysis^105^ was employed to assess branch support within the species tree, which can easily distinguish significant topological discrepancies from instances of insufficient branch information. The QS analysis, conducted with 100 replicates and a log-likelihood cutoff of 2, yielded four pivotal metrics, which could offer a comprehensive understanding of each node in the species tree. The results of the QS analysis were visualized utilizing the plot_QC_ggtree.R script (https://github.com/ShuiyinLIU/QS_visualization).

### Incomplete Lineage Sorting analyses

In this investigation, we employed two distinct methods to explore the influence of ILS as a potential explanation for the observed discordance between gene trees and the species tree. The CoSi analysis could provide a holistic view of the impact and prevalence of ILS across the entire phylogeny. To complement this approach, we also conducted the MuCCo analysis, an approach for assessing the influence of ILS at individual nodes within the phylogenetic tree, thereby offering insight into the specific contributions of ILS to the observed phylogenetic incongruences. In this research, we adapted and refined the scripts for these two methods to accommodate our DGS data. The enhanced scripts, along with comprehensive procedural documentation for both methods, are now publicly accessible on GitHub (https://github.com/PhyloAI/Coalescent-Simulation-Analysis; https://github.com/PhyloAI/Mutation-Calculation-based-on-Coalescent-Model).

In the CoSi analysis, subtrees were extracted from gene trees, followed by the inference of a species tree employing ASTRAL-III with these subtrees. Subsequently, utilizing the “sim.coaltree.sp” function within the R package Phybase v. 2.0^106^, 10,000 gene trees were simulated based on the species tree generated in the preceding step, employing a coalescent model. The topological variance between empirical gene trees and the simulated counterparts was estimated using DendroPy v. 4.5.2^107^, to assess the effect of ILS on the conflicts among gene trees.

In the MuCCo analysis, we further explored the contribution of ILS by calculating the population mutation parameter “theta”^108^. Initially, the branch length information of the species tree was converted from coalescent units to mutation units, utilizing alignments matrix and a species tree as a fixed topology in RAxML v. 8.2.13^101^. Subsequently, the theta of each internal branch was calculated by dividing the branch length in mutation units estimated from RAxML by the length in coalescent units obtained from the ASTRAL species tree. A theta value exceeding 0.1 indicates a substantial degree of ILS.

### SNP calling and geneflow analyses

The genome of *Prunus mume* (accession number: GCF000346735.1) was served as the reference genome for Single Nucleotide Polymorphism (SNP) calling. We used BWA v. 0.7.17^93^ to facilitate the reads mapping to the reference genome, whereas SAMtools v. 1.19^109^ was instrumental in determining the relative position of these reads. Additionally, Picard v. 3.1.1 (http://broadinstitute.github.io/picard/) was employed to mark duplicates within the sorted BAM files. For the detection of genetic variation and the combination of genomic variant call format (gVCF) files into a singular VCF file, we applied the “HaplotypeCaller” and “CombineGVCFs” functions, respectively, as implemented in GATK4 v. 4.5.0.0^110^. The filtration of variants from the combined VCF file was conducted using the functions, i.e., “VariantFiltration”, “SelectVariants”, and “VCFtools”^111^.

To elucidate the dynamics of gene flow among closely related species within the polyploid *Prunus* lineage, particularly within the *Maddenia* group, we employed the Dsuite package^57^ to calculate the *f*_4_-ratio statistics. Utilizing the combined VCF files and an ASTRAL-derived phylogenetic tree as inputs, the “Dtrios” command calculates *f*_4_-ratio statistics for all possible trios of species. We applied the Benjamini-Hochberg (BH) correction method using the “p.adjust” (https://rdocumentation.org/packages/stats/versions/3.6.2/topics/p.adjust) function in the R package, thereby controlling the false discovery rate. Subsequent visualization of the adjusted data was accomplished via the Ruby script “plot_d.rb” (https://github.com/mmatschiner/tutorials/tree/master/analysis_of_introgression_with_snp_data). Additionally, the heuristic approach “Fbranch” was utilized to analyze the gene flow of specific lineages on the ASTRAL tree related to the species under study, with the visualization of these results by the script “dtools.py”.

### Flow cytometric analysis

To explore the diversity of ploidy/genome size variations among species within polyploid *Prunus*, we selected several representative samples of the *Maddenia* group, which has been posited to have undergone multiple allopolyploidization events. This study applied a recently established method for DNA flow cytometry^112^, preparing samples combined with 250 μL nuclear isolation buffer PVPK12-mGB_2_. Nuclei suspensions were filtered and then stained with Propidium Iodide (PI) under dark conditions for 15 minutes and treated with RNase to prevent staining of double-stranded RNA. The analyses were performed on a LSRFortessa flow cytometer (BD, USA), with a minimum of 5000 nuclei analyzed in total. Data acquisition was done through Cell Quest Pro software, and the coefficient of variation (CV) along with the debris factor (DF) were evaluated using Flow JO software^112,113^.

### Multiple methods for whole-genome duplications analyses

Species within the polyploid *Prunus* exhibited a notable increase in paralog numbers compared to the diploid counterparts, indicating the potential WGD events in the origin of the polyploid *Prunus*. To identify and analyze these WGD events, we employed the “wgd_map” pipeline^4,114^. The process started with extracting rooted orthogroups from the final homolog trees, utilizing the script “extract_clades.py”. Subsequently, the “map_dups.py” script was applied to map these orthogroups onto the species tree. This step could document the proportion of gene duplications at the nodes corresponding to the MRCA of the involved clades.

We also adapted the scripts of Tree2GD to accommodate DGS data (https://github.com/PhyloAI/DGS-Tree2GD). All samples of the polyploid *Prunus* samples along with an outgroup, *Lyonothamnus floribundus*, were used for this analysis. We extracted 29,705 genes from the *Prunus mume* genome (accession number: GCF000346735.1) and assembled these nuclear coding sequences (nuclear CDS) using HybPiper v. 2.1.7^88^, creating a new dataset for WGD analysis. The nuclear CDS were translated using TransDecoder v. 5.7.1 (Haas, BJ. https://github.com/TransDecoder/TransDecoder) with default settings. Diamond v. 2.1.9^115^ was then used to build an index and perform all-against-all sequence alignment for each pair of species, using the “more-sensitive” model and filtering results with an e-value greater than 1e-10. Homologous sequences were predicted using phyloMCL v. 2.0^116^, with species less than four discarded, resulting in the retention of 59,212 gene clusters. We used the dolloparsimony script^30^ to map the genes in each gene family to the species tree, calculating the total number of gene families at each node and counting newly obtained and lost gene families. Multiple CDS alignments for each gene cluster were performed using MUSCLE v. 5.1^117^ with “--cds2tree”. Each phylogenetic gene tree was inferred using IQ-TREE2 v. 2.3.4^98^ with the “GTR” model and 1,000 bootstraps. To detect retained gene duplicates (GDs) on lineages, gene trees were reconciled with a reference species tree by Tree2GD v. 1.0.40^30^. Nodes with GD values > 300, and a percentage of GD with both lineages A and B (both taxa A and B having two copies, ABAB type of GD) > 50% were proposed as WGD events.

### Allopolyploidy analyses

In our WGD analyses, we observed that the nodes corresponding to the MRCAs of the polyploid *Prunus* and *Maddenia* group displayed a significant frequency of gene duplication, which strongly suggests potential WGD events at these two nodes. We performed GRAMPA v. 1.4.0^48^ to investigate the polyploidy modes (allopolyploid or autopolyploid) associated with inferred WGD events. The parameter “h1” was used to explore the mode of polyploidization (auto- vs. allo-) and to identify the potential parental lineages involved. Additionally, given the uncertainty about the presence of allo-, auto-, or non-polyploid events in the *Maddenia* group, we tested all possible nodes comprehensively followed the strategies of Morales-Briones et al.^4^

### Historical biogeographic analysis

To elucidate the spatiotemporal dynamics within the polyploid *Prunus*, with a particular emphasis on the *Maddenia* group, we estimated divergence times using ML trees derived from nuclear and plastid CDS datasets. Because of the distinct evolutionary rates among various lineages, uneven taxon sampling and distribution of calibrating fossils across the phylogeny can introduce potential errors in molecular dating analysis. We adopted a two-step approach to achieve accurate molecular dating for the polyploid *Prunus*. Initially, we sampled all major clades within the subfamily Amygdaloideae and selected eight fossil records (F1–F8) and one secondary calibration node (C1) throughout the phylogeny (detailed in Supplementary Methods and Supplementary Table 3), to ensure precise divergence time estimations for the polyploid *Prunus*. Subsequently, for the divergence time estimations of the *Maddenia* group, we used the estimated dating times from the stem node of polyploid *Prunus* as the secondary calibration point, alongside one fossil record of *P. wutuensis* (F2).

Considering the computational burden in dating analysis, we used the approximate likelihood calculation applied by the program MCMCTREE in PAML v. 4.10.7^118^. We opted for the “independent rates” clock model and the “HKY85” model for nucleotide substitution and configured the parameters as follows: initiating with a burn-in period of 1,000,000 iterations, sampling frequency was set to every 10 iterations and the total number of samples collected was specified as 500,000. The assessment of stationarity and convergence across these runs was conducted, ensuring that the effective sample size (ESS) for all parameters is larger than a minimum threshold of 200.

In the second step, we utilized the tribe Lyonothamneae as the outgroup in the SCN genes dataset and the tribe Sorbarieae as the outgroup in the plastid CDS dataset. The estimated stem ages of *Prunus*—83.98–64.47 Mya in the SCN dataset and 86.84–52.36 Mya in the plastid CDS dataset— served as the secondary calibration points. To ensure methodological consistency, we applied the same parameters as those used in the first step.

To estimate the ancestral geographic origin of the polyploid *Prunus* lineage, we employed the BioGeoBEARS v. 1.1.1^119^ implemented in the Reconstruct Ancestral State in Phylogenies (RASP) v. 4.1^120^. We defined six distinct biogeographic regions based on the distribution patterns in the polyploid *Prunus*, and they are as follows: (A) East Asia, (B) North & South America, (C) West Asia, (D) Europe, (E) Africa, and (F) Australasia. Furthermore, to investigate the origin and dispersal of the *Maddenia* group, we defined three geographic areas, including (A) Southeast China (Fujian, Zhejiang, Anhui, and Jiangxi); (B) Southwest China (Xizang and Yunnan); (C) Central China (Gansu, Qinghai, Shaanxi, Hubei, Hunan, Henan, Chongqing, and Sichuan). In these analyses, the “DIVALIKE” model was selected due to its ability to accommodate vicariance, dispersal, and extinction events.

### Diversification rates analysis

RevBayes v. 1.2.1^121^ was used to estimate the speciation and extinction rate dynamics over time within the *Maddenia* group, grounded in the episodic birth-death (EBD) model^122,123^. The analyses continued until the ESS value of the estimated model parameters exceeded a threshold of 200 after burn-in. The magnitude and timing of rate changes were then visualized using the R package RevGadgets v1.0.0^124^. These analyses utilized both the nuclear MO and plastid CDS datasets to explore how divergent phylogenies might affect the assessment of diversification rates.

## Supporting information

Supplemental Figures and their legends, legends of Supplemental Tables 1-3, Supplemental Methods

## Data availability

All the DGS data are deposited in the NCBI SRA under the BioProject PRJNA1031385, and the detailed information for each sample is referred to Supplementary Table 1. Sequence alignments, phylogenetic trees, and other data files generated in this study are deposited at Dryad Digital Repository [https://datadryad.org/stash/share/zrSRWvefAjRapx7TElIyJAh0uKcmofE41va3HnVH4o8].

## Code availability

All newly generated and adapted codes for this study are available on GitHub at https://github.com/PhyloAI.

## Acknowledgements

In this study, computational analyses were conducted on the PhyloAI supercomputer, a high-performance computing resource owned by Bin-Bin Liu (https://doi.org/10.12282/PhyloAIHPC). Concurrently, molecular experiments were performed at the Plant DNA and Molecular Identification Platform (PDMIP) at IBCAS and the laboratory at Northwest A&F University. We also extend our gratitude to all members of the PhyloAI team. Financial support for this work was provided by the National Natural Science Foundation of China (32170381 and 31770200 to LZ, 32270216 and 32000163 to BBL), the Youth Innovation Promotion Association CAS (2023086 to BBL), and the Biological Taxonomy Scientist Position at the Chinese Academy of Sciences (CAS-TAX-24-013 to BBL). The study was supported by National Wild Plant Germplasm Resource Center for Chenshan (ZWGX2402 to XZ).

## Author contributions

B.B.L., L.Z., and R.G.J.H. conceived and designed the study, with B.B.L. leading and supervising the project. S.Y.X. and X.H.L. drafted the manuscript. L.Z., S.Y.X., J.R.W., and C.X. were responsible for collecting the materials. S.Y.X., D.K.M. and Y.Z. conducted the experiments. S.Y.X., X.H.L., and J.R.W. performed the phylogenomic analyses. P.L., X.Z., and B.J.G. contributed DGS data of two samples (*Prunus gongshanensis* 1 & 2) and one sample of silica-gel dried leaves (*Prunus himalayana* 4), while H.M. supplied all the RNA-Seq data used in this study. R.G.J.H. and D.Y.C. offered advice on structuring the paper. All authors participated in writing and interpreting the results and approved the final manuscript.

## Additional information

### Supplementary information

The online version contains supplementary material available at http://doi.org/…

## References

1. Jiao, Y. N. Double the genome, double the fun: genome fuplications in angiosperms. Mol. Plant. 11, 357–358 (2018).

2. One Thousand Plant Transcriptomes Initiative. One thousand plant transcriptomes and the phylogenomics of green plants. Nature 574, 679–685 (2019).

3. Rothfels, C. J. Polyploid phylogenetics. New Phytol. 230, 66–72 (2021).

4. Morales-Briones, D. F. et al. Analysis of paralogs in target enrichment data pinpoints multiple ancient polyploidy events in *Alchemilla* s.l. (Rosaceae). Syst. Biol. 71, 190–207 (2022).

5. Stull, G. W., Pham, K. K., Soltis, P. S. & Soltis, D. E. Deep reticulation: the long legacy of hybridization in vascular plant evolution. Plant J. 114, 743–766 (2023).

6. Xu, C. et al. Dense sampling of taxa and genomes untangles the phylogenetic backbone of a non-model plant lineage rife with deep hybridization and allopolyploidy. Preprint at 10.1101/2023.10.21.563444 (2023).

7. You, Y. C. et al. Transition of survival strategies under global climate shifts in the grape family. Nat. Plants 10, 1100–1111 (2024).

8. Liu, B. B. et al. Phylogenomic conflict analyses in the apple genus *Malus* s.l. reveal widespread hybridization and allopolyploidy driving diversification, with insights into the complex biogeographic history in the Northern Hemisphere. J. Integr. Plant Biol. 64, 1020–1043 (2022).

9. Yu, J. R. et al. Integrated phylogenomic analyses unveil reticulate evolution in *Parthenocissus* (Vitaceae), highlighting speciation dynamics in the Himalayan-Hengduan Mountains. New Phytol. 238, 888–903 (2023).

10. Yu, J. R. et al. Distinct hybridization modes in wide- and narrow-ranged lineages of *Causonis* (Vitaceae). BMC Biol. 21, 209 (2023).

11. Wan, J. N. et al. The rise of baobab trees in Madagascar. Nature 629, 1091–1099 (2024).

12. Hendriks, K. P. et al. Global Brassicaceae phylogeny based on filtering of 1,000-gene dataset. Curr. Biol. 33, 4052–4068.e1-e6 (2023).

13. Lu, L. D. et al. Rosaceae. in Flora of China Vol. 9 (eds. Wu, Z. Y., Raven, P. H. & Hong, D. Y.) 46–434 (Science Press; Missouri Botanical Graden Press, Beijing; St. Louis, 2003).

14. Chin, S.-W., Shaw, J., Haberle, R., Wen, J. & Potter, D. Diversification of almonds, peaches, plums and cherries - molecular systematics and biogeographic history of *Prunus* (Rosaceae). Mol. Phylogenet. Evol. 76, 34–48 (2014).

15. Phipps, J. B. & Brouillet, L. Rosaceae. in Flora of North America North of Mexico Vol. 9 18–662 (Oxford University Press, New York and Oxford, 2014).

16. Hodel, R. G. J., Zimmer, E. & Wen, J. A phylogenomic approach resolves the backbone of *Prunus* (Rosaceae) and identifies signals of hybridization and allopolyploidy. Mol. Phylogenet. Evol. 160, 107118 (2021).

17. Hodel, R. G. J. et al. A phylogenomic approach, combined with morphological characters gleaned via machine learning, uncovers the hybrid origin and biogeographic diversification of the plum genus. Preprint at 10.1101/2023.09.13.557598 (2023).

18. Zhao, L. et al. Multiple events of allopolyploidy in the evolution of the racemose lineages in *Prunus* (Rosaceae) based on integrated evidence from nuclear and plastid data. PLoS ONE 11, e0157123 (2016).

19. Su, N. et al. On the species delimitation of the *Maddenia* group of *Prunus* (Rosaceae): evidence from plastome and nuclear sequences and morphology. Front. Plant Sci. 12, 743643 (2021).

20. Wu, D. D. et al. Pervasive introgression facilitated domestication and adaptation in the *Bos* species complex. *Nat*. Ecol. Evol. 2, 1139–1145 (2018).

21. Li, N. et al. DNA methylation repatterning accompanying hybridization, whole genome doubling and homoeolog exchange in nascent segmental rice allotetraploids. New Phytol. 223, 979–992 (2019).

22. Wang, Y. P. et al. MCScanX: a toolkit for detection and evolutionary analysis of gene synteny and collinearity. Nucleic Acids Res. 40, e49 (2012).

23. Haug-Baltzell, A., Stephens, S. A., Davey, S., Scheidegger, C. E. & Lyons, E. SynMap2 and SynMap3D: web-based whole-genome synteny browsers. Bioinformatics 33, 2197–2198 (2017).

24. Lynch, M. & Conery, J. S. The evolutionary fate and consequences of duplicate genes. Science 290, 1151–1155 (2000).

25. Jiao, Y. N. et al. Ancestral polyploidy in seed plants and angiosperms. Nature 473, 97–100 (2011).

26. Li, Z. et al. Early genome duplications in conifers and other seed plants. Sci. Adv. 1, e1501084 (2015).

27. Yang, Y. et al. Dissecting molecular evolution in the highly diverse plant clade Caryophyllales using transcriptome sequencing. Mol. Biol. Evol. 32, 2001–2014 (2015).

28. Huang, C.-H. et al. Resolution of Brassicaceae phylogeny using nuclear genes uncovers nested radiations and supports convergent morphological evolution. Mol. Biol. Evol. 33, 394–412 (2016).

29. Xiang, Y. Z. et al. Evolution of Rosaceae fruit types based on nuclear phylogeny in the context of geological times and genome duplication. Mol. Biol. Evol. 34, 262–281 (2017).

30. Chen, D. Y., Zhang, T. K., Chen, Y. M., Ma, H. & Qi, J. Tree2GD: a phylogenomic method to detect large-scale gene duplication events. Bioinformatics 38, 5317–5321 (2022).

31. Weitemier, K. et al. Hyb-Seq: combining target enrichment and genome skimming for plant phylogenomics. Appl. Plant Sci. 2, apps.1400042 (2014).

32. Liu, B. B. et al. Capturing single-copy nuclear genes, organellar genomes, and nuclear ribosomal DNA from deep genome skimming data for plant phylogenetics: a case study in Vitaceae. J. Syst. Evol. 59, 1124–1138 (2021).

33. Forrest, L. L. et al. The limits of Hyb-Seq for herbarium specimens: impact of preservation techniques. Front. Ecol. Evol. 7, 439 (2019).

34. Cooper, B. J. et al. Target enrichment and extensive population sampling help untangle the recent, rapid radiation of *Oenothera* sect. *Calylophus*. Syst. Biol. 72, 249–263 (2023).

35. Ren, R. et al. Widespread whole genome duplications contribute to genome complexity and species diversity in angiosperms. Mol. Plant. 11, 414–428 (2018).

36. Guo, J. et al. Phylotranscriptomics in Cucurbitaceae reveal multiple whole-genome duplications and key morphological and molecular innovations. Mol. Plant. 13, 1117–1133 (2020).

37. Huang, C.-H., Qi, X. P., Chen, D. Y., Qi, J. & Ma, H. Recurrent genome duplication events likely contributed to both the ancient and recent rise of ferns. J. Integr. Plant Biol. 62, 433–455 (2020).

38. Zhao, Y. Y. et al. Nuclear phylotranscriptomics and phylogenomics support numerous polyploidization events and hypotheses for the evolution of rhizobial nitrogen-fixing symbiosis in Fabaceae. Mol. Plant. 14, 748–773 (2021).

39. Zhang, L. et al. Phylotranscriptomics resolves the phylogeny of Pooideae and uncovers factors for their adaptive evolution. Mol. Biol. Evol. 39, msac026 (2022).

40. Zhang, L. et al. Phylogenomics insights into gene evolution, rapid species diversification, and morphological innovation of the apple tribe (Maleae, Rosaceae). New Phytol. 240, 2102–2120 (2023).

41. Zhang, Q. et al. Phylotranscriptomic analyses reveal deep gene tree discordance in *Camellia* (Theaceae). Mol. Phylogenet. Evol. 188, 107912 (2023).

42. Jin, Z. T. et al. Nightmare or delight: taxonomic circumscription meets reticulate evolution in the phylogenomic era. Mol. Phylogenet. Evol. 189, 107914 (2023).

43. Jin, Z. T. et al. Unraveling the Web of Life: incomplete lineage sorting and hybridization as primary mechanisms over polyploidization in the evolutionary dynamics of pear species. Preprint at 10.1101/2024.07.29.605463 (2024).

44. Shen, X. et al. Evolution of cherries (*Prunus* Subgenus *Cerasus*) based on chloroplast genomes. Int. J. Plant Sci. 24, 15612 (2023).

45. Tan, Q. P. et al. Chromosome-level genome assemblies of five *Prunus* species and genome-wide association studies for key agronomic traits in peach. Hortic. Res. 8, 213 (2021).

46. Watkins, R. Cherry, plum, peach, apricot and almond: *Prunus* spp. (Rosaceae). in Evolution of Crop Plants (ed. Simmons, N. W.) 242–247 (Longman, London, 1976).

47. Morales-Briones, D. F. et al. Disentangling sources of gene tree discordance in phylogenomic data sets: testing ancient hybridizations in Amaranthaceae s.l. Syst. Biol. 70, 219–235 (2021).

48. Thomas, G. W. C., Ather, S. H. & Hahn, M. W. Gene-tree reconciliation with MUL-trees to resolve polyploidy events. Syst. Biol. 66, 1007–1018 (2017).

49. Shi, W. T., Wen, J. & Lutz, S. Pollen morphology of the *Maddenia* clade of *Prunus* and its taxonomic and phylogenetic implications. J. Syst. Evol. 51, 164–183 (2013).

50. Lee, S. & Wen, J. A phylogenetic analysis of *Prunus* and the Amygdaloideae (Rosaceae) using ITS sequences of nuclear ribosomal DNA. Am. J. Bot. 88, 150–160 (2001).

51. Wen, J. et al. Phylogenetic inferences in *Prunus* (Rosaceae) using chloroplast *ndh*F and nuclear ribosomal ITS sequences. J. Syst. Evol. 46, 322–332 (2008).

52. Liu, X. L. et al. Polyphyly of the *Padus* group of *Prunus* (Rosaceae) and the evolution of biogeographic disjunctions between eastern Asia and eastern North America. J. Plant Res. 126, 351–361 (2013).

53. Zhang, R. G., Shang, H. Y., Jia, K. H. & Ma, Y. P. Subgenome phasing for complex allopolyploidy: case-based benchmarking and recommendations. Brief. Bioinform. 25, bbad513 (2024).

54. Durvasula, A. & Sankararaman, S. Recovering signals of ghost archaic introgression in African populations. Sci. Adv. 6, eaax5097 (2020).

55. Zachos, J., Pagani, M., Sloan, L., Thomas, E. & Billups, K. Trends, rhythms, and aberrations in global climate 65 Ma to present. Science 292, 686–693 (2001).

56. He, Z. Y. et al. New constraints on the late Oligocene-Miocene thermo-tectonic evolution of the southeastern Tibetan Plateau from low-temperature thermochronology. Tectonics 42, e2023TC007881 (2023).

57. Malinsky, M., Matschiner, M. & Svardal, H. Dsuite - Fast *D*-statistics and related admixture evidence from VCF files. Mol. Ecol. Resour. 21, 584–595 (2021).

58. Yang, L. H. et al. Phylogenomic analyses reveal an allopolyploid origin of core Didymocarpinae (Gesneriaceae) followed by rapid radiation. Syst. Biol. 72, 1064–1083 (2023).

59. Debray, K. et al. Unveiling the patterns of reticulated evolutionary processes with phylogenomics: hybridization and polyploidy in the genus *Rosa*. Syst. Biol. 71, 547–569 (2022).

60. Ye, X. Y. et al. Rapid diversification of alpine bamboos associated with the uplift of the Hengduan Mountain. J. Biogeogr. 46, 2678–2689 (2019).

61. Mo, Z. Q. et al. Resolution, conflict and rate shifts: insights from a densely sampled plastome phylogeny for *Rhododendron* (Ericaceae). Ann. Bot. 130, 687–701 (2022).

62. Zhu, W. L. et al. Subduction evolution controlled Himalayan orogenesis: implications from 3-D subduction modeling. Appl. Sci. 12, 7413 (2022).

63. Ding, L. et al. Timing and mechanisms of Tibetan Plateau uplift. Nat. Rev. Earth Environ. 3, 652–667 (2022).

64. Cui, L. Y. et al. Widespread genome duplications throughout the history of flowering plants. Genome Res. 16, 738–749 (2006).

65. Jiao, Y. N. et al. A genome triplication associated with early diversification of the core eudicots. Genome Biol. 13, R3 (2012).

66. Soltis, P. S. & Soltis, D. E. Ancient WGD events as drivers of key innovations in angiosperms. Curr. Opin. Plant Biol. 30, 159–165 (2016).

67. Masterson, J. Stomatal size in fossil plants: evidence for polyploidy in majority of angiosperms. Science 264, 421–424 (1994).

68. Van de Peer, Y., Maere, S. & Meyer, A. The evolutionary significance of ancient genome duplications. Nat. Rev. Genet. 10, 725–732 (2009).

69. Wendel, J. F. The wondrous cycles of polyploidy in plants. Am. J. Bot. 102, 1753–1756 (2015).

70. Cheng, F. et al. Gene retention, fractionation and subgenome differences in polyploid plants. Nat. Plants 4, 258–268 (2018).

71. Yang, Y. & Smith, S. A. Orthology inference in nonmodel organisms using transcriptomes and low-coverage genomes: improving accuracy and matrix occupancy for phylogenomics. Mol. Biol. Evol. 31, 3081–3092 (2014).

72. Cannon, S. B. et al. Multiple polyploidy events in the early radiation of nodulating and nonnodulating legumes. Mol. Biol. Evol. 32, 193–210 (2015).

73. Liu, B. B. et al. Phylogenomic analyses of the *Photinia* complex support the recognition of a new genus *Phippsiomeles* and the resurrection of a redefined *Stranvaesia* in Maleae (Rosaceae). J. Syst. Evol. 57, 678–694 (2019).

74. Särkinen, T., Staats, M., Richardson, J. E., Cowan, R. S. & Bakker, F. T. How to open the treasure chest? Optimising DNA extraction from herbarium specimens. PLoS ONE 7, e43808 (2012).

75. Saeidi, S., McKain, M. R. & Kellogg, E. A. Robust DNA isolation and high-throughput sequencing library construction for herbarium specimens. J. Vis. Exp. e56837 (2018

76. Guo, C. et al. Phylogenomics and the flowering plant tree of life. J. Integr. Plant Biol. 65, 299– 323 (2023).

77. Stiller, J. et al. Complexity of avian evolution revealed by family-level genomes. Nature 629, 851–860 (2024).

78. Lemmon, A. R., Emme, S. A. & Lemmon, E. M. Anchored hybrid enrichment for massively high-throughput phylogenomics. Syst. Biol. 61, 727–744 (2012).

79. Mandel, J. R. et al. A target enrichment method for gathering phylogenetic information from hundreds of loci: an example from the Compositae. Appl. Plant Sci. 2, 1300085 (2014).

80. Dodsworth, S. et al. Hyb-Seq for flowering plant systematics. Trends Plant Sci. 24, 887–891 (2019).

81. Li, Z. et al. Multiple large-scale gene and genome duplications during the evolution of hexapods. Proc. Natl. Acad. Sci. USA 115, 4713–4718 (2018).

82. Doyle, J. J. & Doyle, J. L. A rapid DNA isolation procedure for small quantities of fresh leaf tissue. Phytochem. Bull. 19, 11–15 (1987).

83. Li, J. L., Wang, S., Yu, J., Wang, L. & Zhou, S. L. A modified CTAB protocol for plant DNA extraction. Chin. Bull. Bot. 48, 72–78 (2013).

84. Bolger, A. M., Lohse, M. & Usadel, B. Trimmomatic: a flexible trimmer for Illumina sequence data. Bioinformatics 30, 2114–2120 (2014).

85. Jin, J. J. et al. GetOrganelle: a fast and versatile toolkit for accurate de novo assembly of organelle genomes. Genome Biol. 21, 241 (2020).

86. Qu, X. J., Moore, M. J., Li, D. Z. & Yi, T. S. PGA: a software package for rapid, accurate, and flexible batch annotation of plastomes. Plant Methods 15, 50 (2019).

87. Kearse, M. et al. Geneious Basic: an integrated and extendable desktop software platform for the organization and analysis of sequence data. Bioinformatics 28, 1647–1649 (2012).

88. Johnson, M. G. et al. HybPiper: extracting coding sequence and introns for phylogenetics from high-throughput sequencing reads using target enrichment. Appl. Plant Sci. 4, 1600016 (2016).

89. Nakamura, T., Yamada, K. D., Tomii, K. & Katoh, K. Parallelization of MAFFT for large-scale multiple sequence alignments. Bioinformatics 34, 2490–2492 (2018).

90. Capella-Gutiérrez, S., Silla-Martínez, J. M. & Gabaldón, T. trimAl: a tool for automated alignment trimming in large-scale phylogenetic analyses. Bioinformatics 25, 1972–1973 (2009).

91. Borowiec, M. L. AMAS: a fast tool for alignment manipulation and computing of summary statistics. PeerJ 4, e1660 (2016).

92. Chamala, S. et al. MarkerMiner 1.0: A new application for phylogenetic marker development using angiosperm transcriptomes. Appl. Plant Sci. 3, 1400115 (2015).

93. Li, H. & Durbin, R. Fast and accurate short read alignment with Burrows-Wheeler transform. Bioinformatics 25, 1754–1760 (2009).

94. Prjibelski, A., Antipov, D., Meleshko, D., Lapidus, A. & Korobeynikov, A. Using SPAdes de novo assembler. Current Protocols in Bioinformatics 70, e102 (2020).

95. Lanfear, R., Frandsen, P. B., Wright, A. M., Senfeld, T. & Calcott, B. PartitionFinder 2: new methods for selecting partitioned models of evolution for molecular and morphological phylogenetic analyses. Mol. Biol. Evol. 34, 772–773 (2017).

96. Lanfear, R., Calcott, B., Ho, S. Y. W. & Guindon, S. Partitionfinder: combined selection of partitioning schemes and substitution models for phylogenetic analyses. Mol. Biol. Evol. 29, 1695–1701 (2012).

97. Lanfear, R., Calcott, B., Kainer, D., Mayer, C. & Stamatakis, A. Selecting optimal partitioning schemes for phylogenomic datasets. BMC Evol. Biol. 14, 82 (2014).

98. Minh, B. Q. et al. IQ-TREE 2: new models and efficient methods for phylogenetic inference in the genomic era. Mol. Biol. Evol. 37, 1530–1534 (2020).

99. Minh, B. Q., Nguyen, M. A. T. & von Haeseler, A. Ultrafast approximation for phylogenetic bootstrap. Mol. Biol. Evol. 30, 1188–1195 (2013).

100. Anisimova, M. & Gascuel, O. Approximate likelihood-ratio test for branches: a fast, accurate, and powerful alternative. Syst. Biol. 55, 539–552 (2006).

101. Stamatakis, A. RAxML version 8: a tool for phylogenetic analysis and post-analysis of large phylogenies. Bioinformatics 30, 1312–1313 (2014).

102. Zhang, C., Rabiee, M., Sayyari, E. & Mirarab, S. ASTRAL-III: polynomial time species tree reconstruction from partially resolved gene trees. BMC Bioinforma. 19, 153 (2018).

103. Huson, D. H. & Bryant, D. Application of phylogenetic networks in evolutionary studies. Mol. Biol. Evol. 23, 254–267 (2006).

104. Smith, S. A., Moore, M. J., Brown, J. W. & Yang, Y. Analysis of phylogenomic datasets reveals conflict, concordance, and gene duplications with examples from animals and plants. BMC Evol. Biol. 15, 150 (2015).

105. Pease, J. B., Brown, J. W., Walker, J. F., Hinchliff, C. E. & Smith, S. A. Quartet Sampling distinguishes lack of support from conflicting support in the green plant tree of life. Am. J. Bot. 105, 385–403 (2018).

106. Liu, L. & Yu, L. L. Phybase: an R package for species tree analysis. Bioinformatics 26, 962–963 (2010).

107. Sukumaran, J. & Holder, M. T. DendroPy: a Python library for phylogenetic computing. Bioinformatics 26, 1569–1571 (2010).

108. Cai, L. M. et al. The perfect storm: gene tree estimation error, incomplete lineage sorting, and ancient gene flow explain the most recalcitrant ancient angiosperm clade, Malpighiales. Syst. Biol. 70, 491–507 (2021).

109. Danecek, P. et al. Twelve years of SAMtools and BCFtools. GigaScience 10, giab008 (2021).

110. Van der Auwera, G. A. & O’Connor, B. D. Genomics in the Cloud: Using Docker, GATK, and WDL in Terra. (O’Reilly Media, Inc., Sebastopol, CA, 2020).

111. Danecek, P. et al. The variant call format and VCFtools. Bioinformatics 27, 2156–2158 (2011).

112. Zhang, J. D. & Feng, M. A plant sample optimal pretreatment for flow cytometric analysis. Chin. Bull. Bot. 58, 285–297 (2023).

113. Loureiro, J., Rodriguez, E., Dolezel, J. & Santos, C. Comparison of four nuclear isolation buffers for plant DNA flow cytometry. Ann. Bot. 98, 679–689 (2006).

114. Yang, Y. et al. Improved transcriptome sampling pinpoints 26 ancient and more recent polyploidy events in Caryophyllales, including two allopolyploidy events. New Phytol. 217, 855–870 (2018).

115. Buchfink, B., Reuter, K. & Drost, H.-G. Sensitive protein alignments at tree-of-life scale using DIAMOND. Nat. Methods 18, 366–368 (2021).

116. Zhou, S. Y., Chen, Y. M., Guo, C. C. & Qi, J. PhyloMCL: accurate clustering of hierarchical orthogroups guided by phylogenetic relationship and inference of polyploidy events. Methods Ecol. Evol. 11, 943–954 (2020).

117. Edgar, R. C. MUSCLE: a multiple sequence alignment method with reduced time and space complexity. BMC Bioinforma. 5, 113 (2004).

118. Yang, Z. H. PAML 4: phylogenetic analysis by maximum likelihood. Mol. Biol. Evol. 24, 1586– 1591 (2007).

119. Matzke, N. J. BioGeoBEARS: BioGeography with Bayesian (and Likelihood) evolutionary analysis with R scripts. (2018).

120. Yu, Y., Blair, C. & He, X. J. RASP 4: ancestral state reconstruction tool for multiple genes and characters. Mol. Biol. Evol. 37, 604–606 (2020).

121. Höhna, S. et al. RevBayes: Bayesian phylogenetic inference using graphical models and an interactive model-specification language. Syst. Biol. 65, 726–736 (2016).

122. Stadler, T. Mammalian phylogeny reveals recent diversification rate shifts. Proc. Natl. Acad. Sci. USA 108, 6187–6192 (2011).

123. Höhna, S. The time-dependent reconstructed evolutionary process with a key-role for mass-extinction events. J. Theor. Biol. 380, 321–331 (2015).

124. Tribble, C. M. et al. RevGadgets: an R package for visualizing Bayesian phylogenetic analyses from RevBayes. Methods Ecol. Evol. 13, 314–323 (2022).

