## Supplemental Figures and their legends, legends of Supplemental Tables 1-3, Supplemental Methods for "Unveiling allopolyploidization-driven genome duplications through progressive analysis of deep genome skimming data"

Supplementary Fig. 1

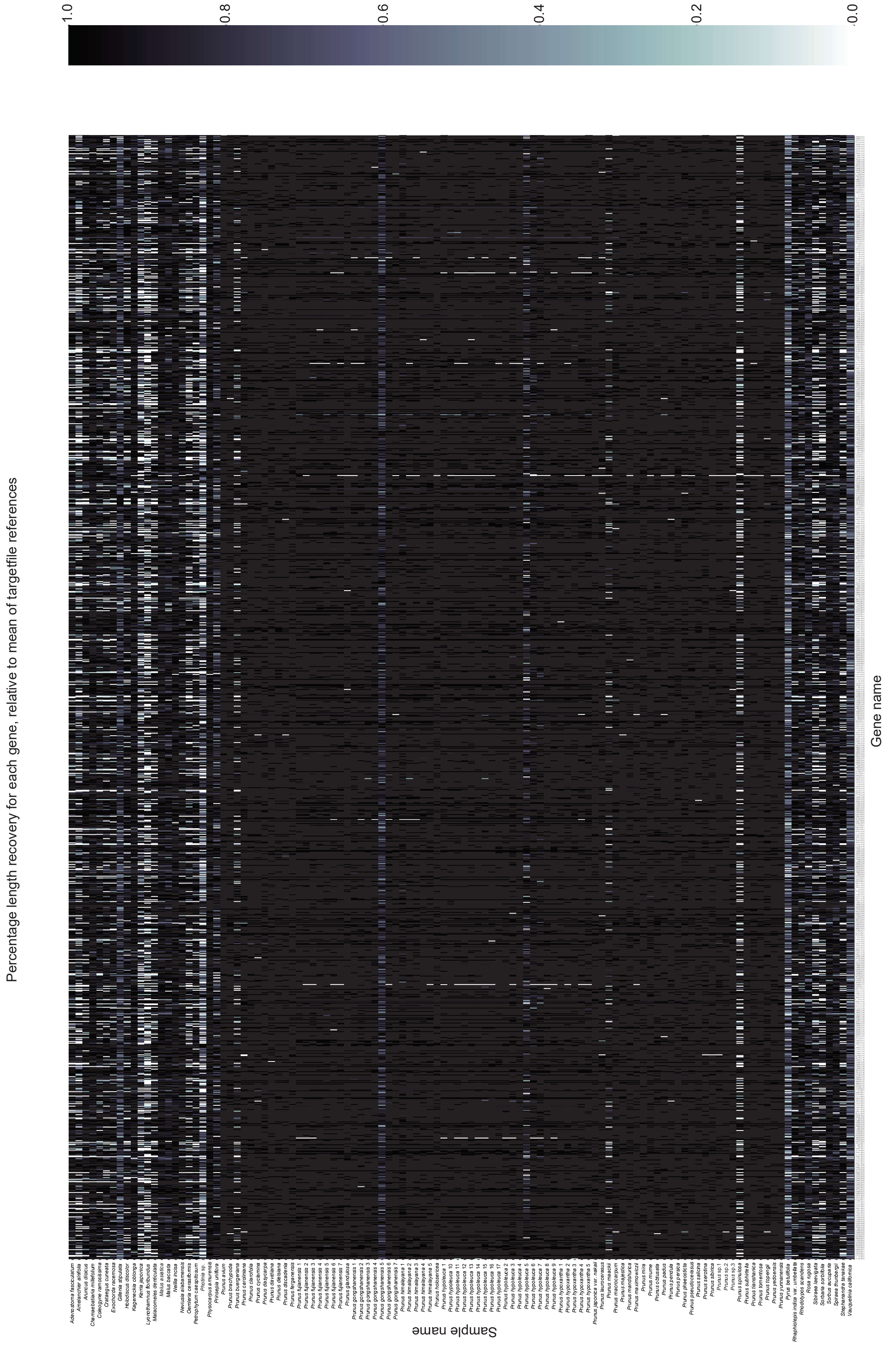

Supplementary Fig. 2

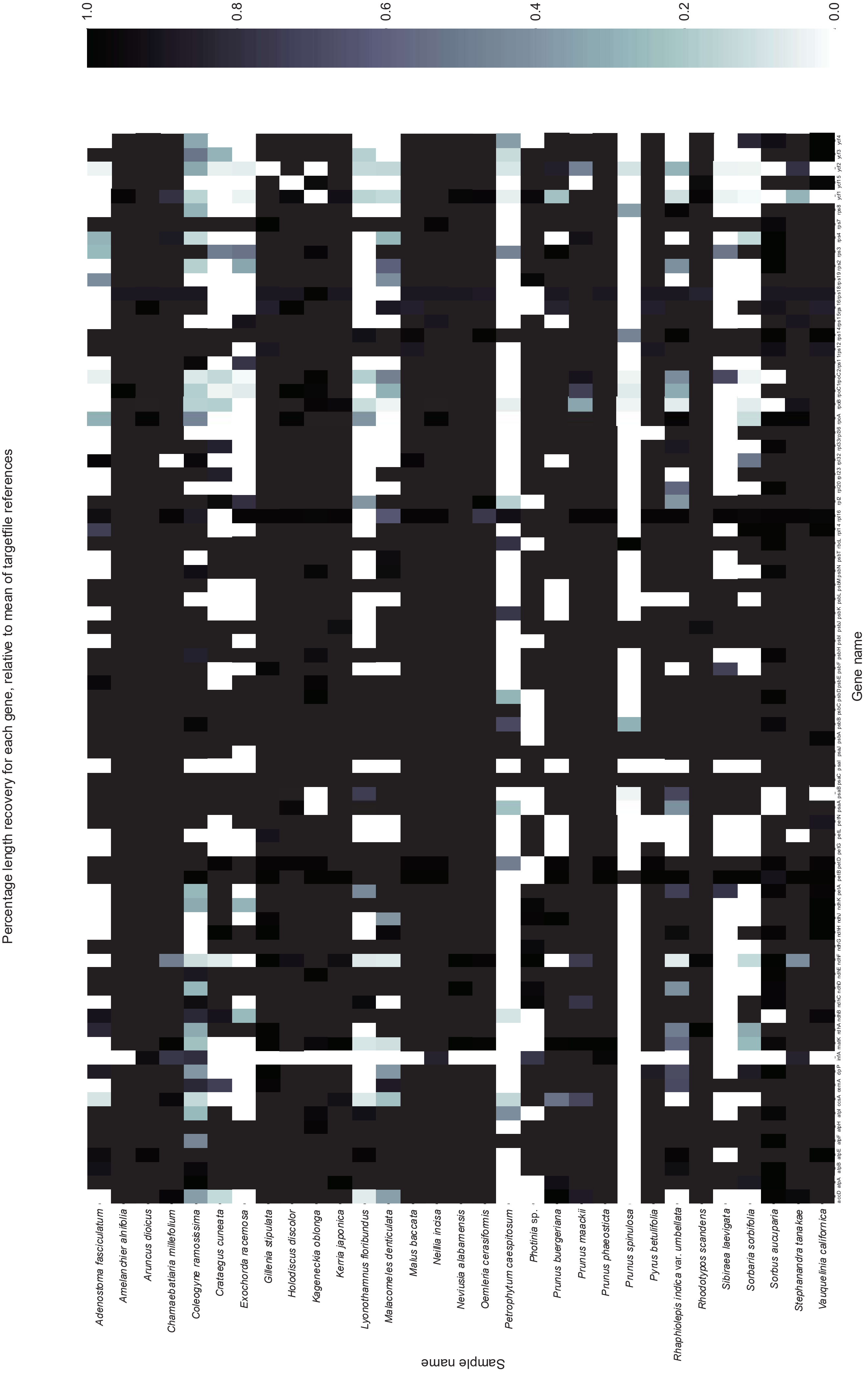

Supplementary Fig. 3

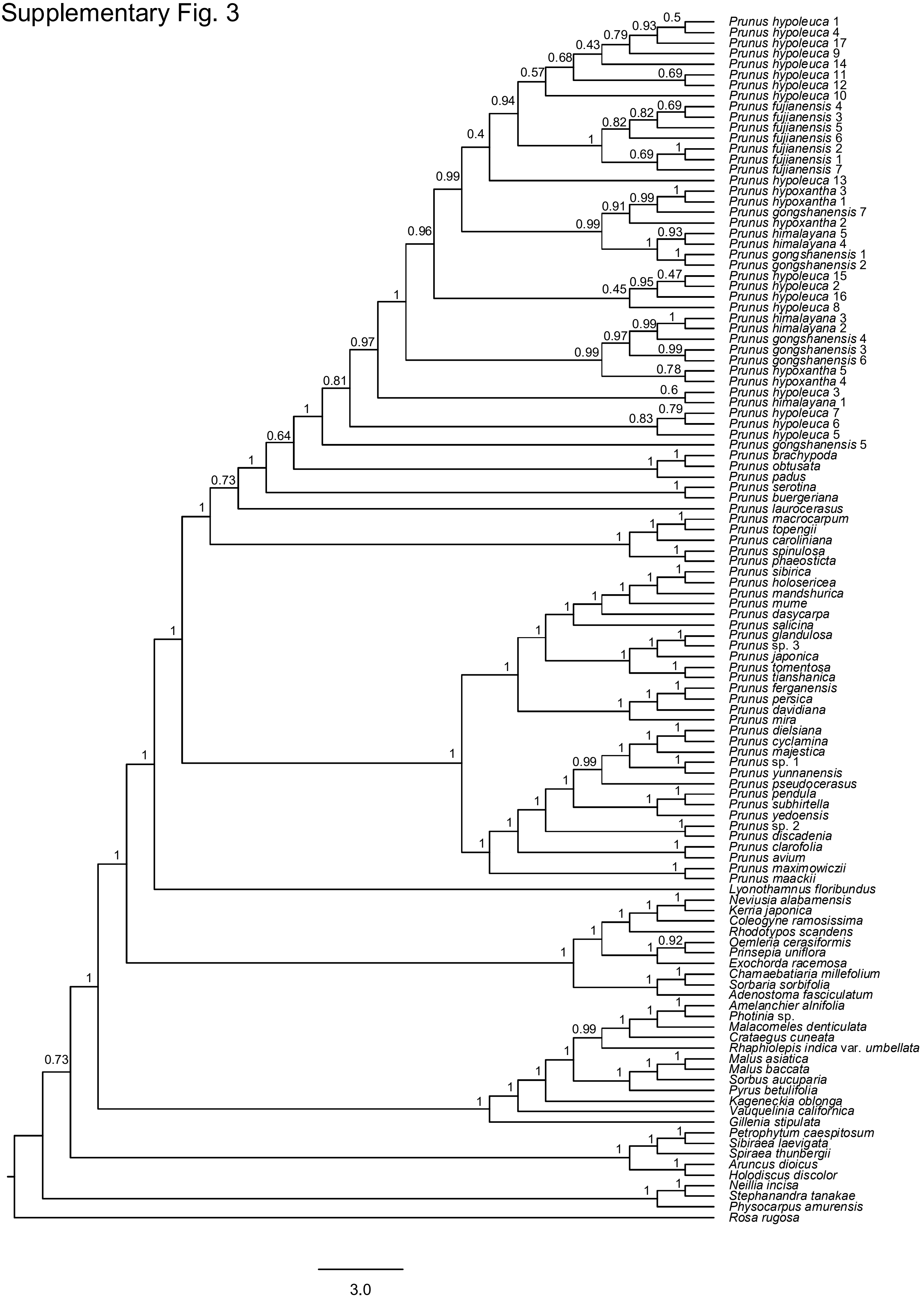

Supplementary Fig. 4

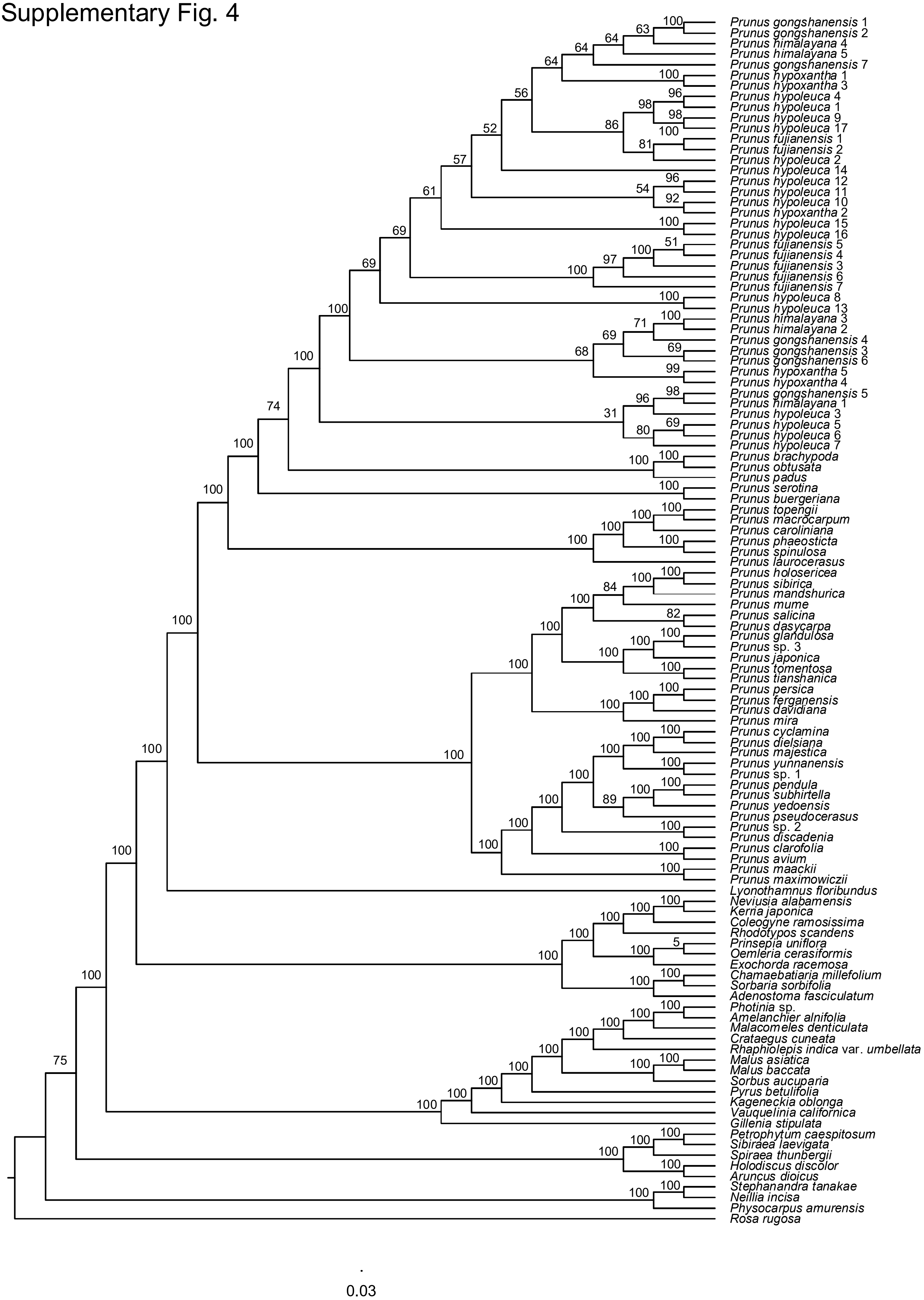

Supplementary Fig. 5

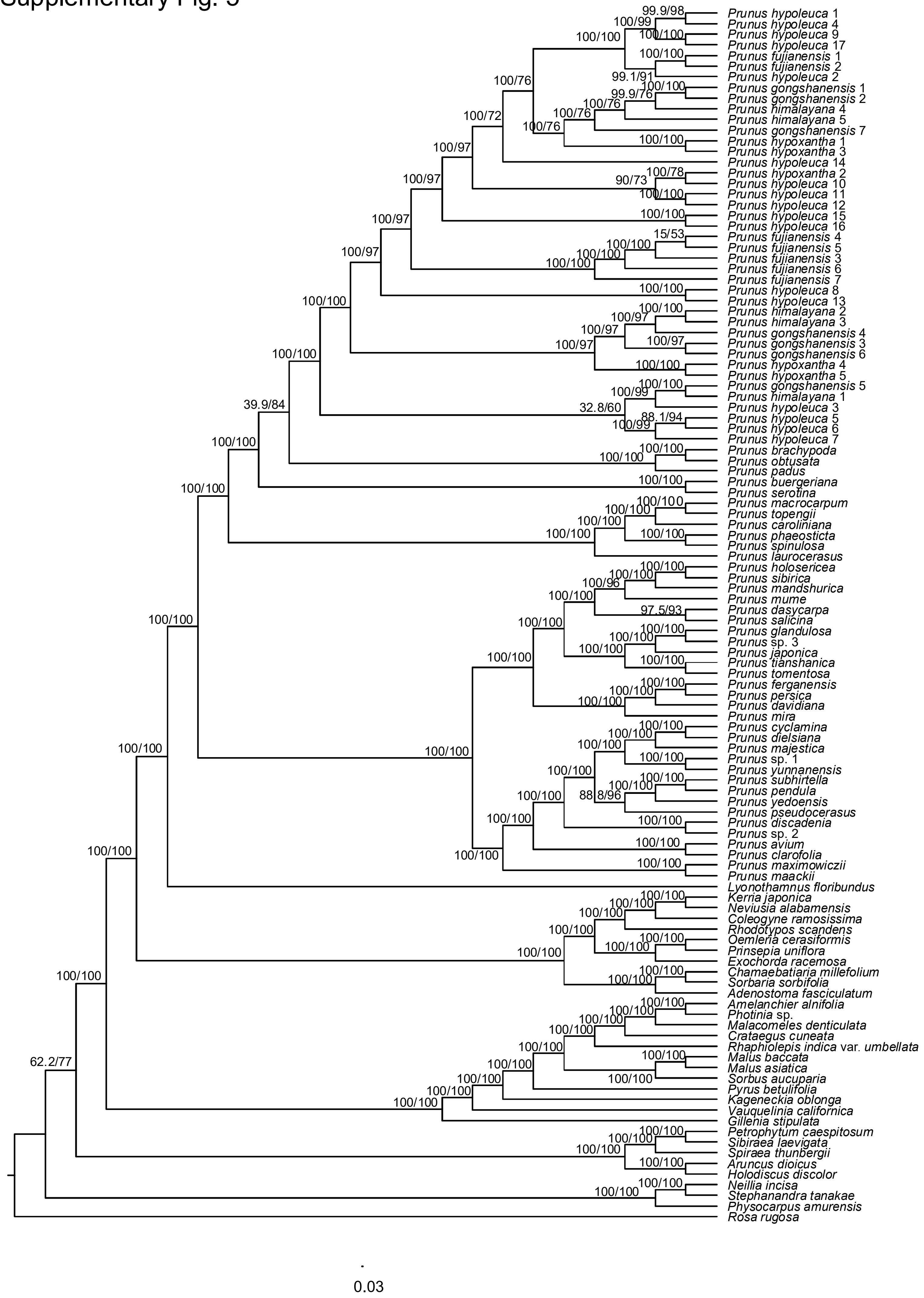

Supplementary Fig. 6

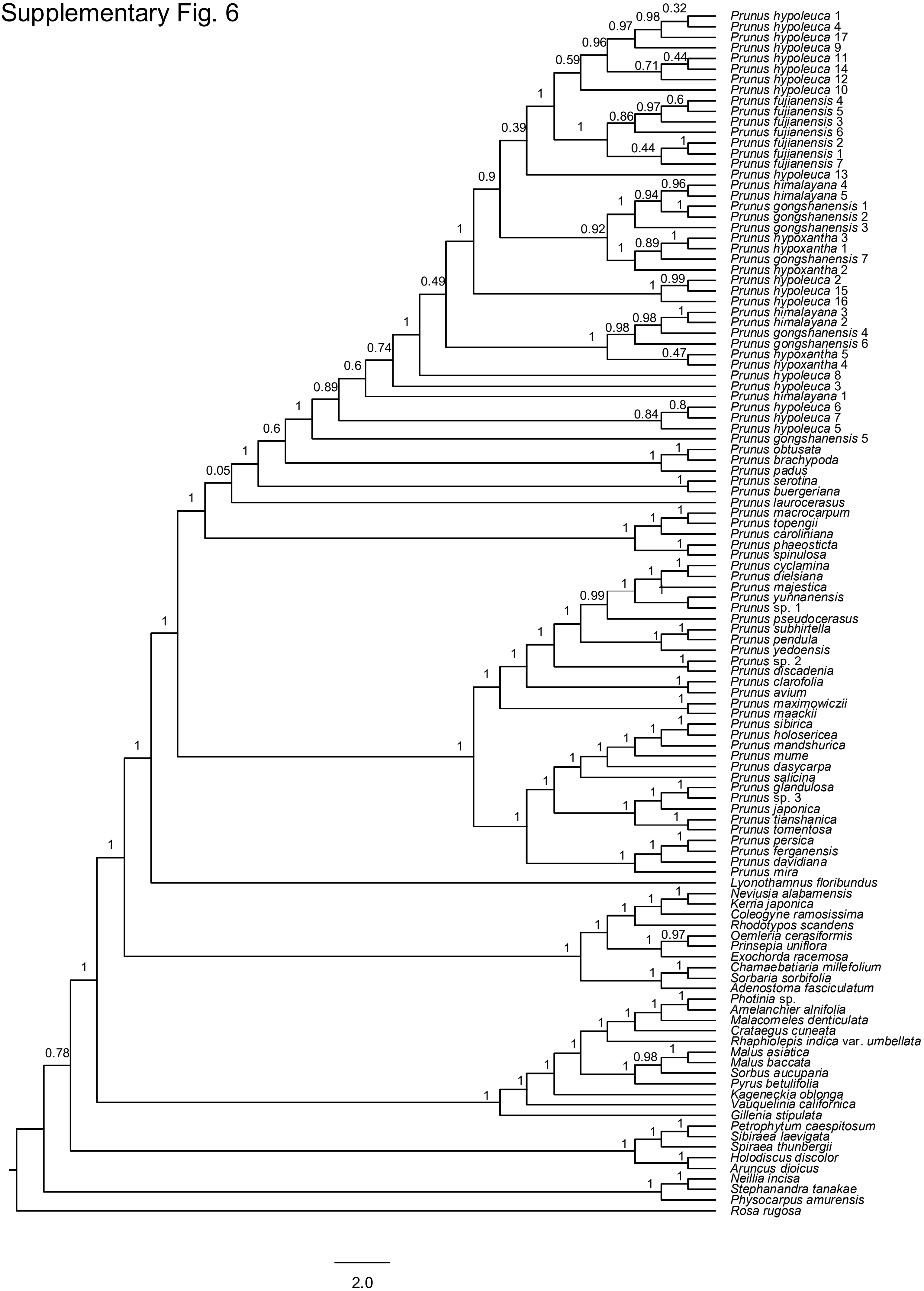

Supplementary Fig. 7

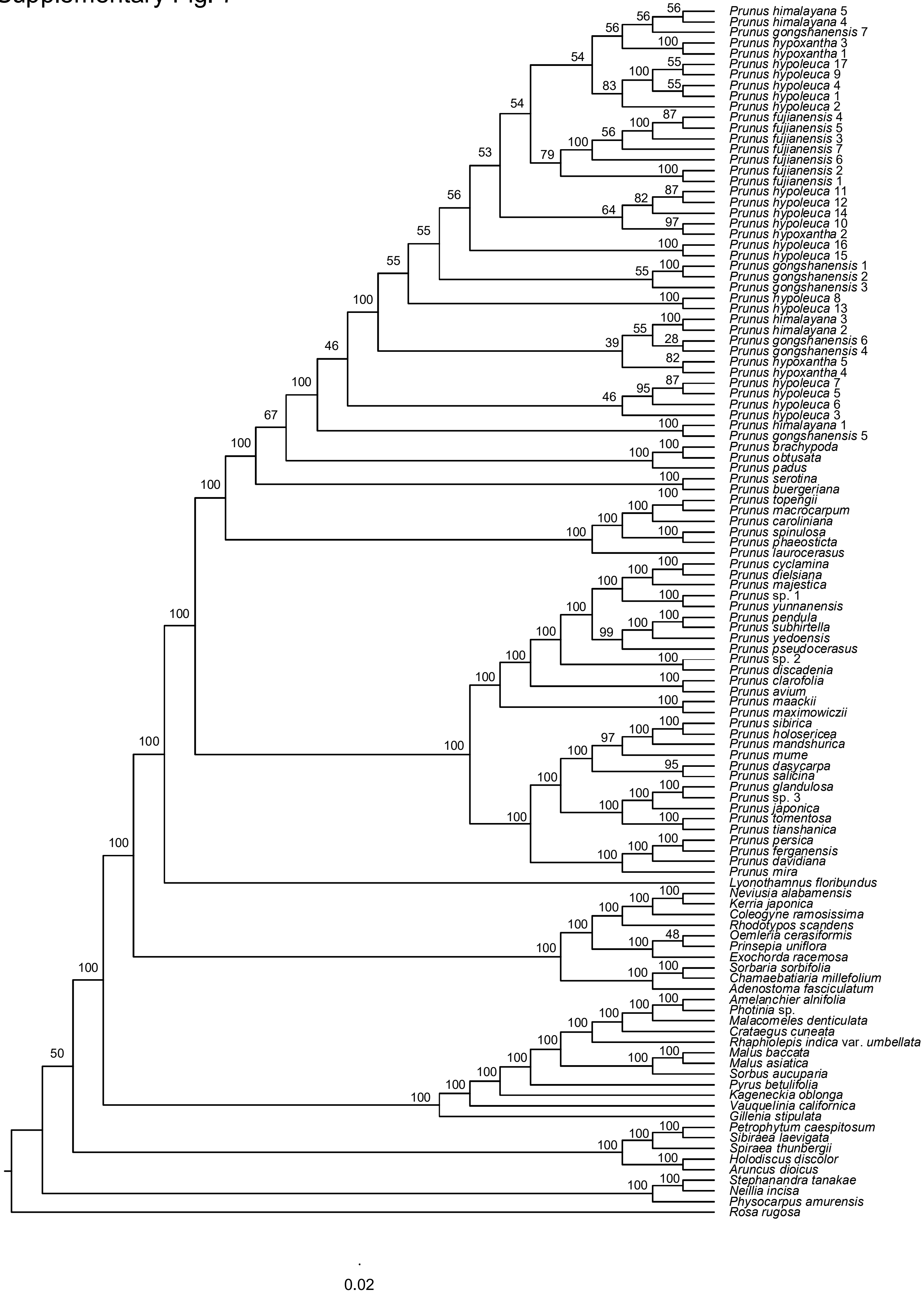

Supplementary Fig. 8

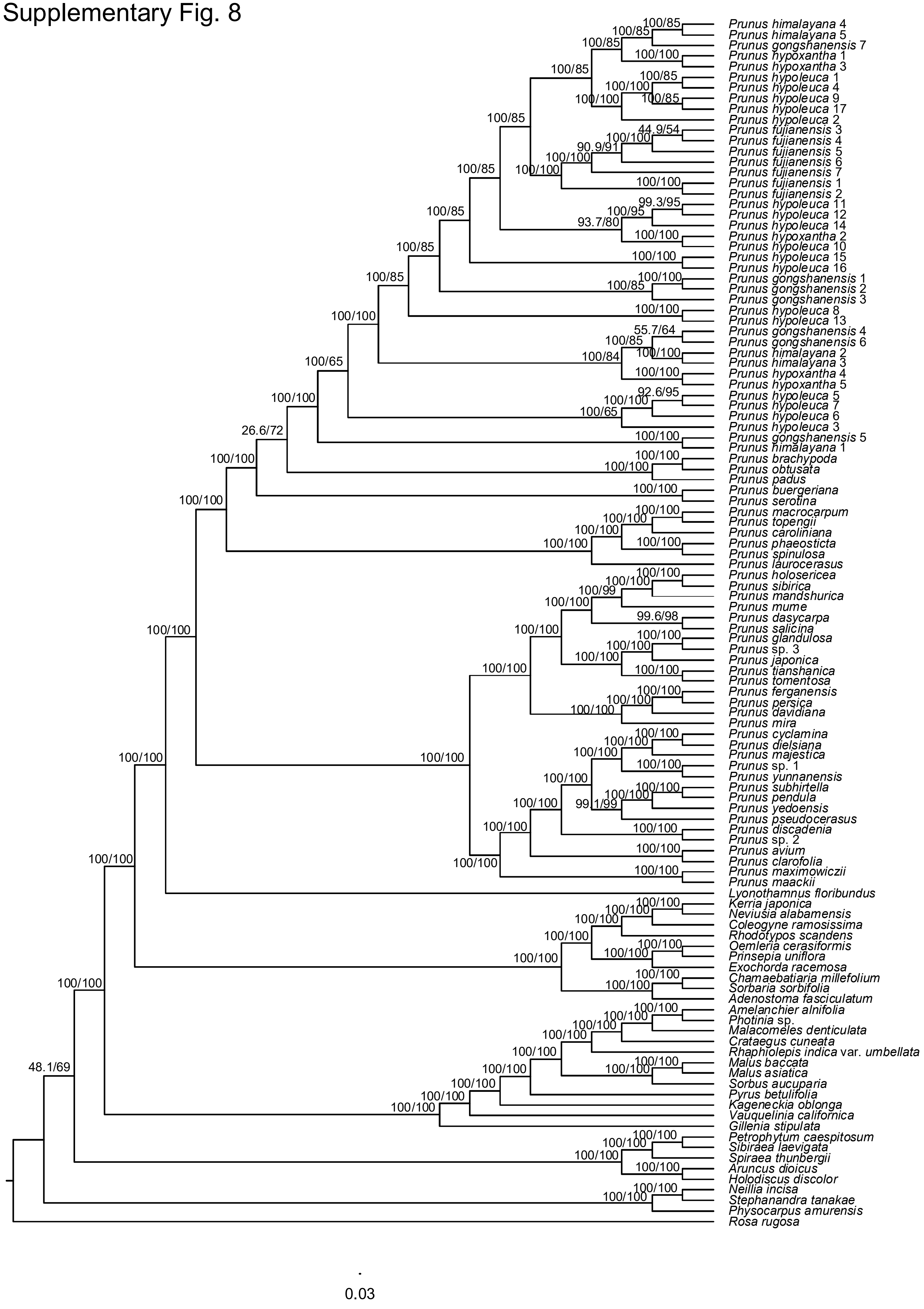

Supplementary Fig. 9

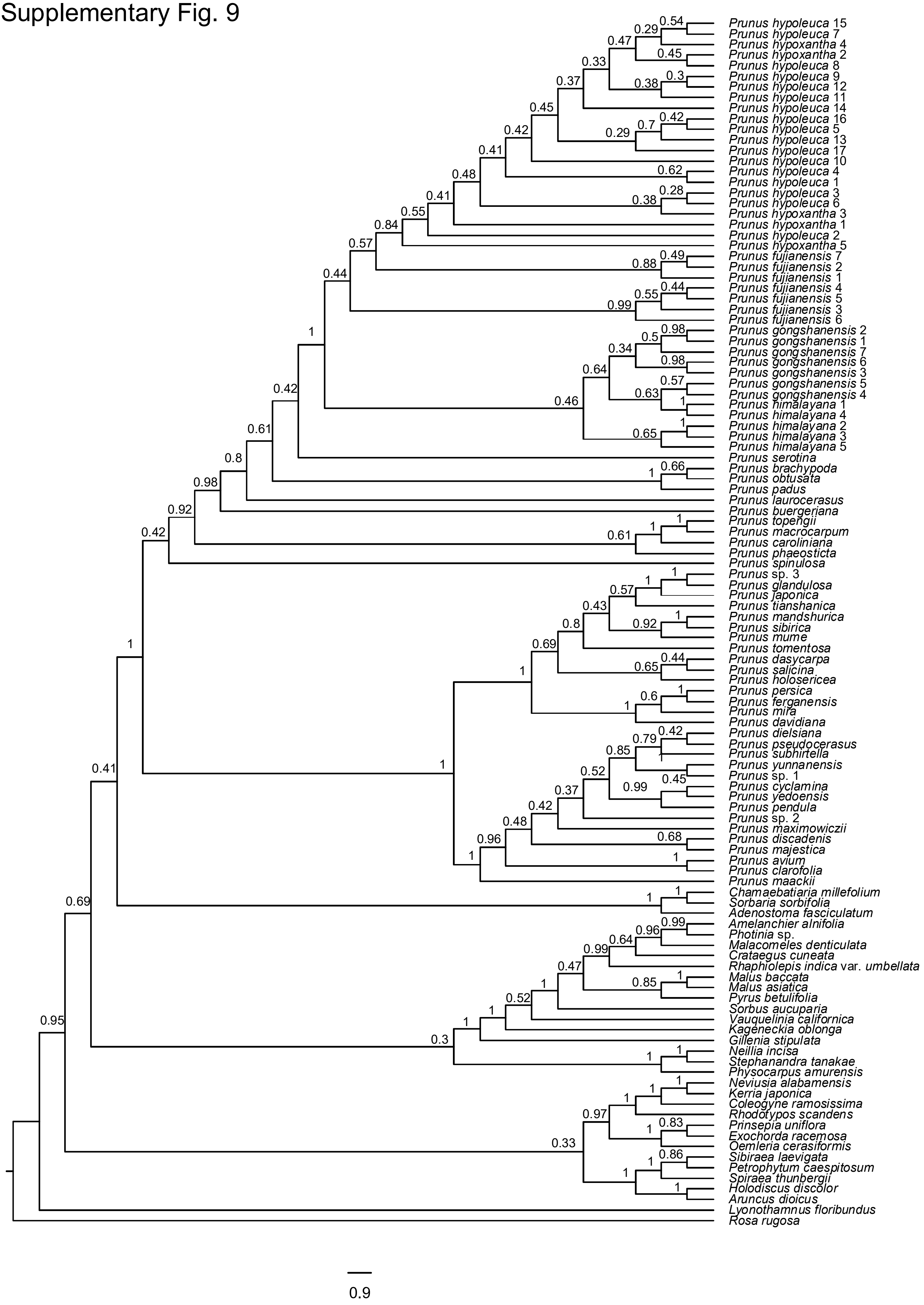

### Supplementary Fig. 10

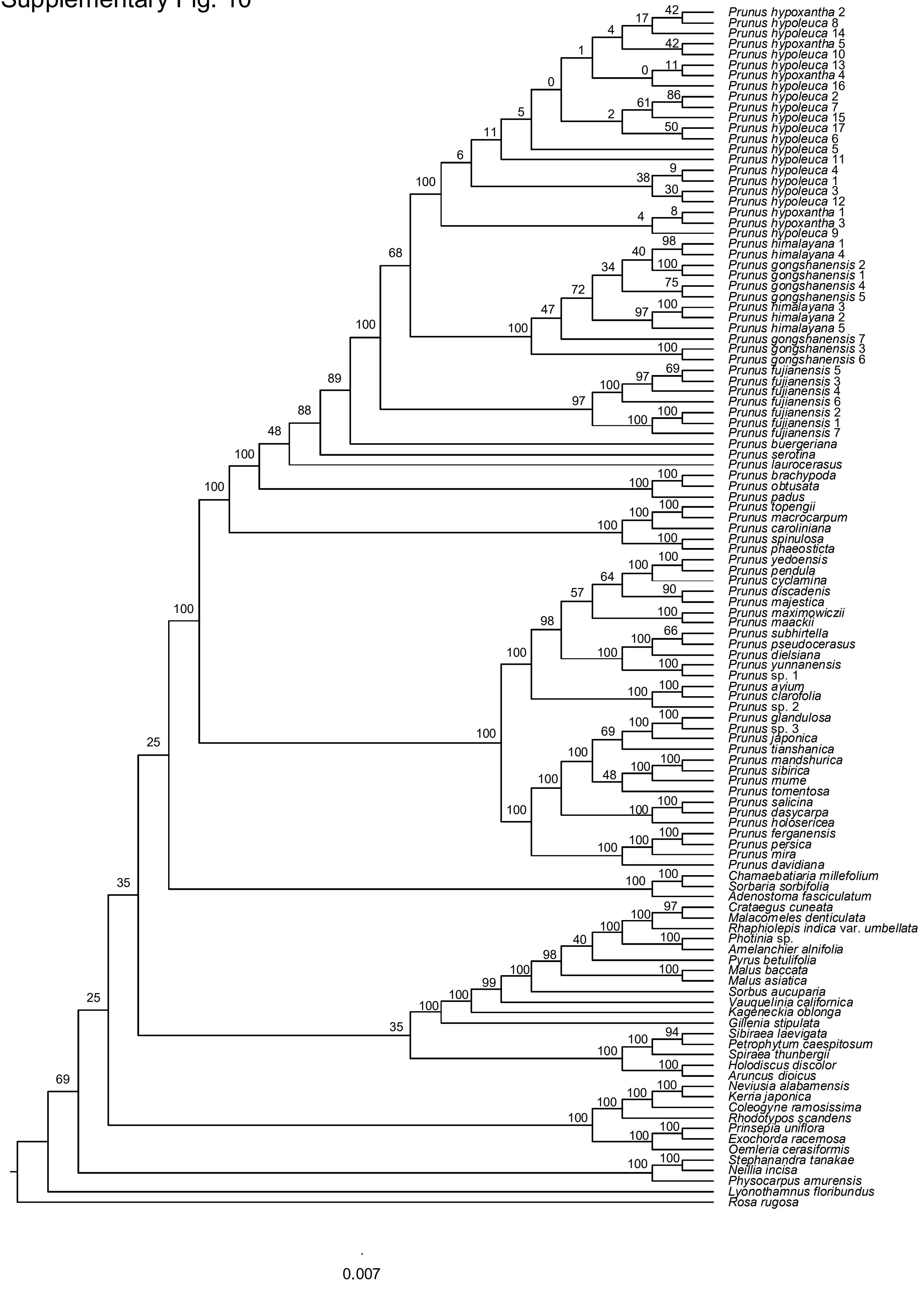

### Supplementary Fig. 11

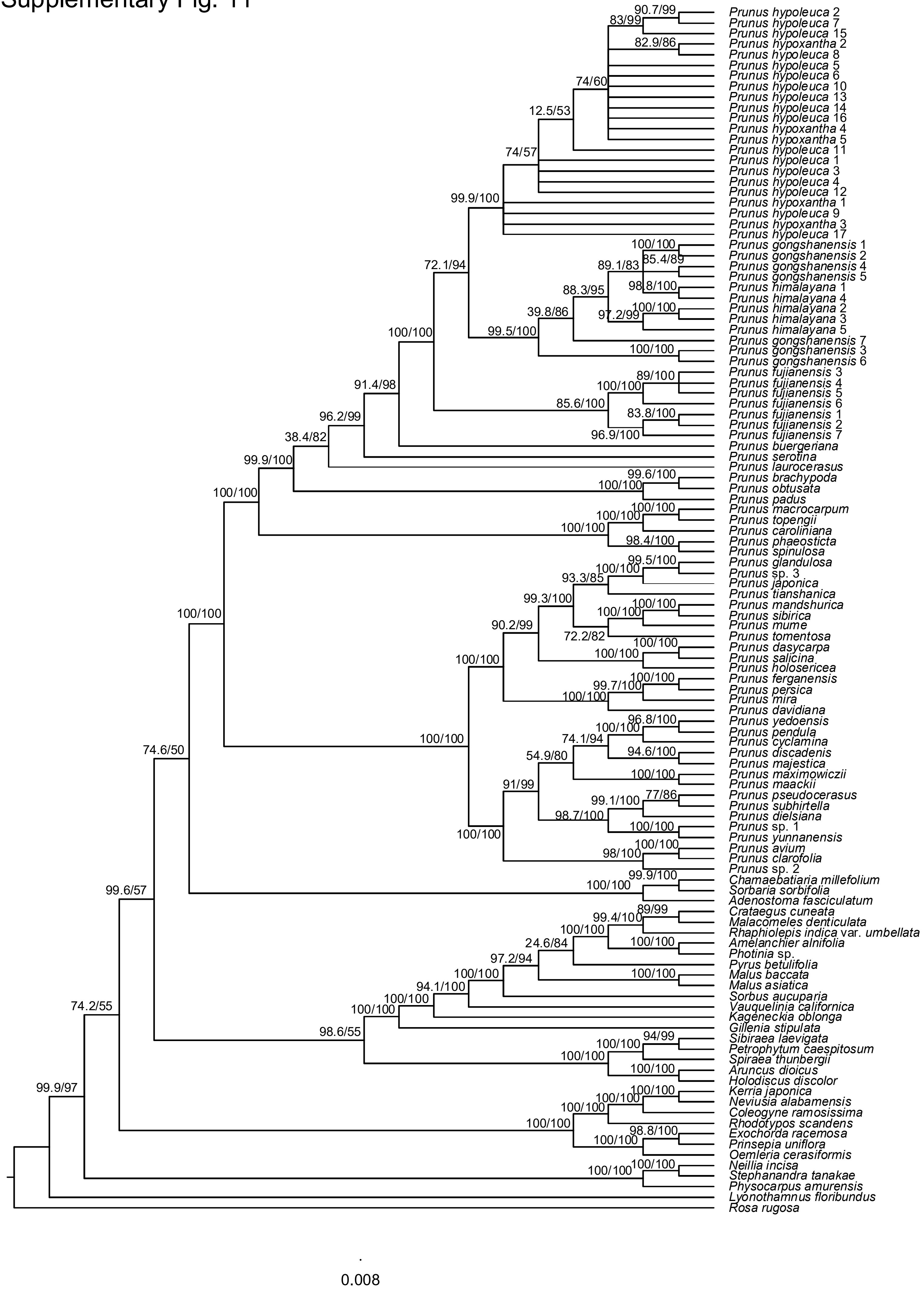

Supplementary Fig. 12

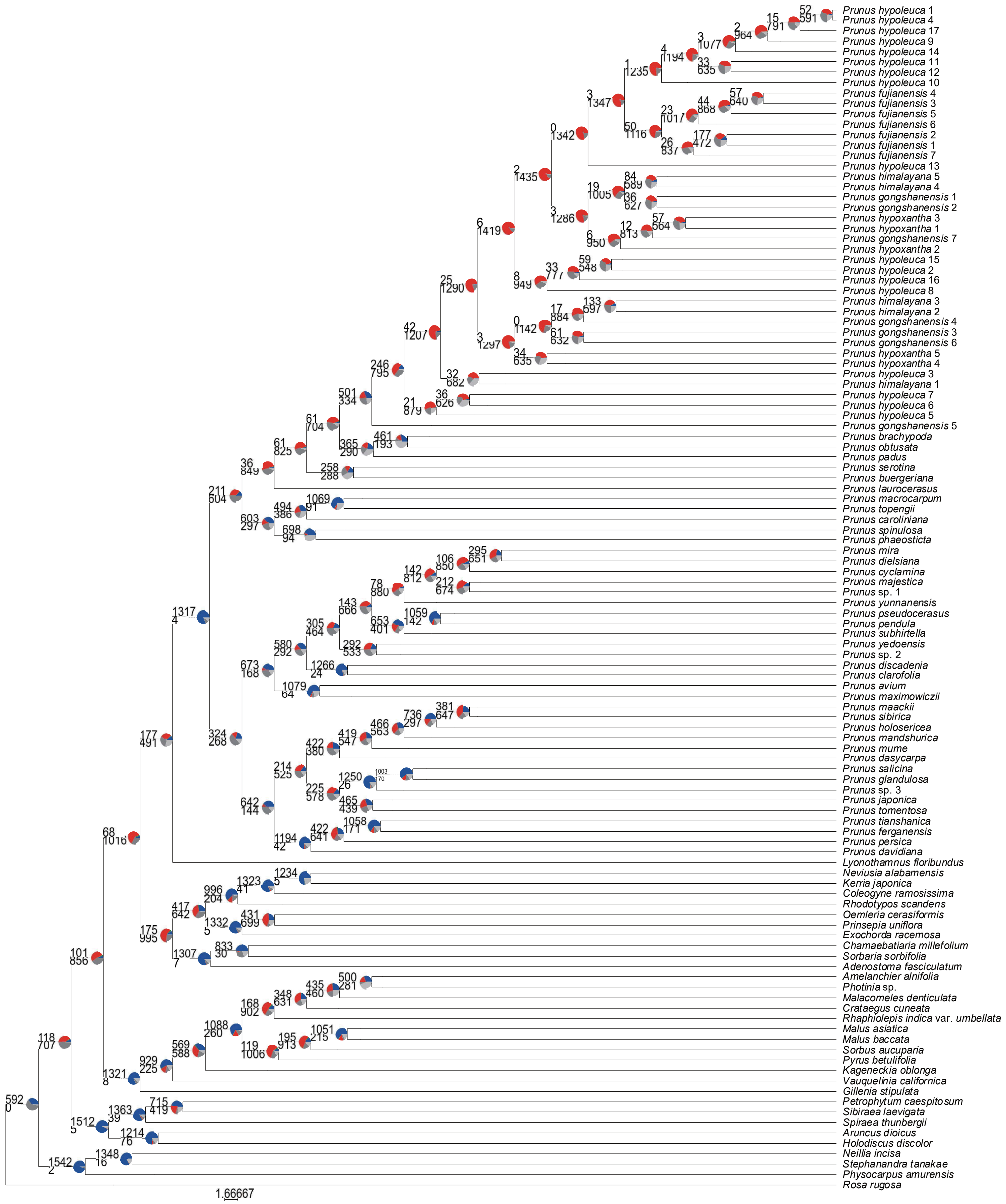

Supplementary Fig. 13

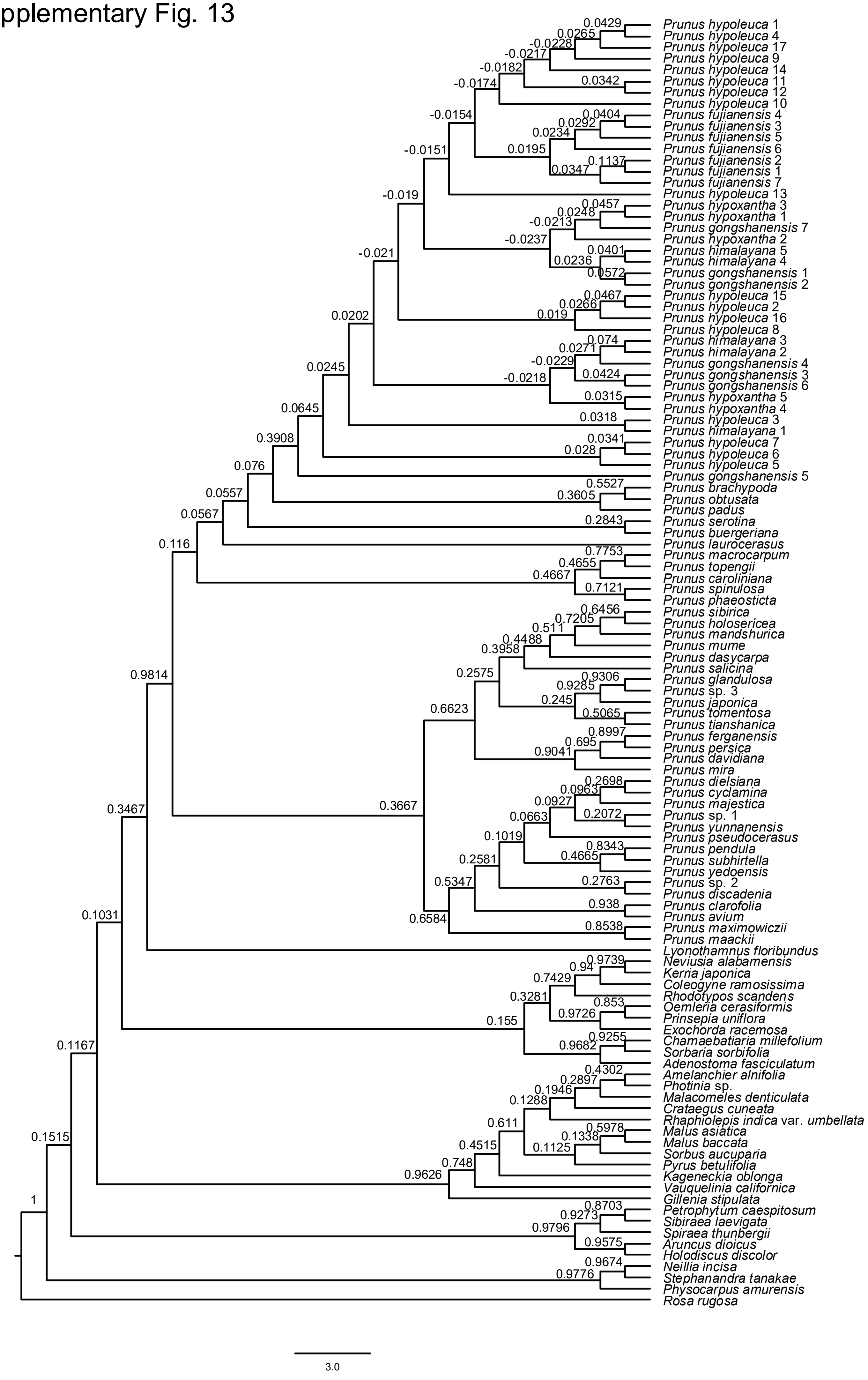

### Supplementary Fig. 14

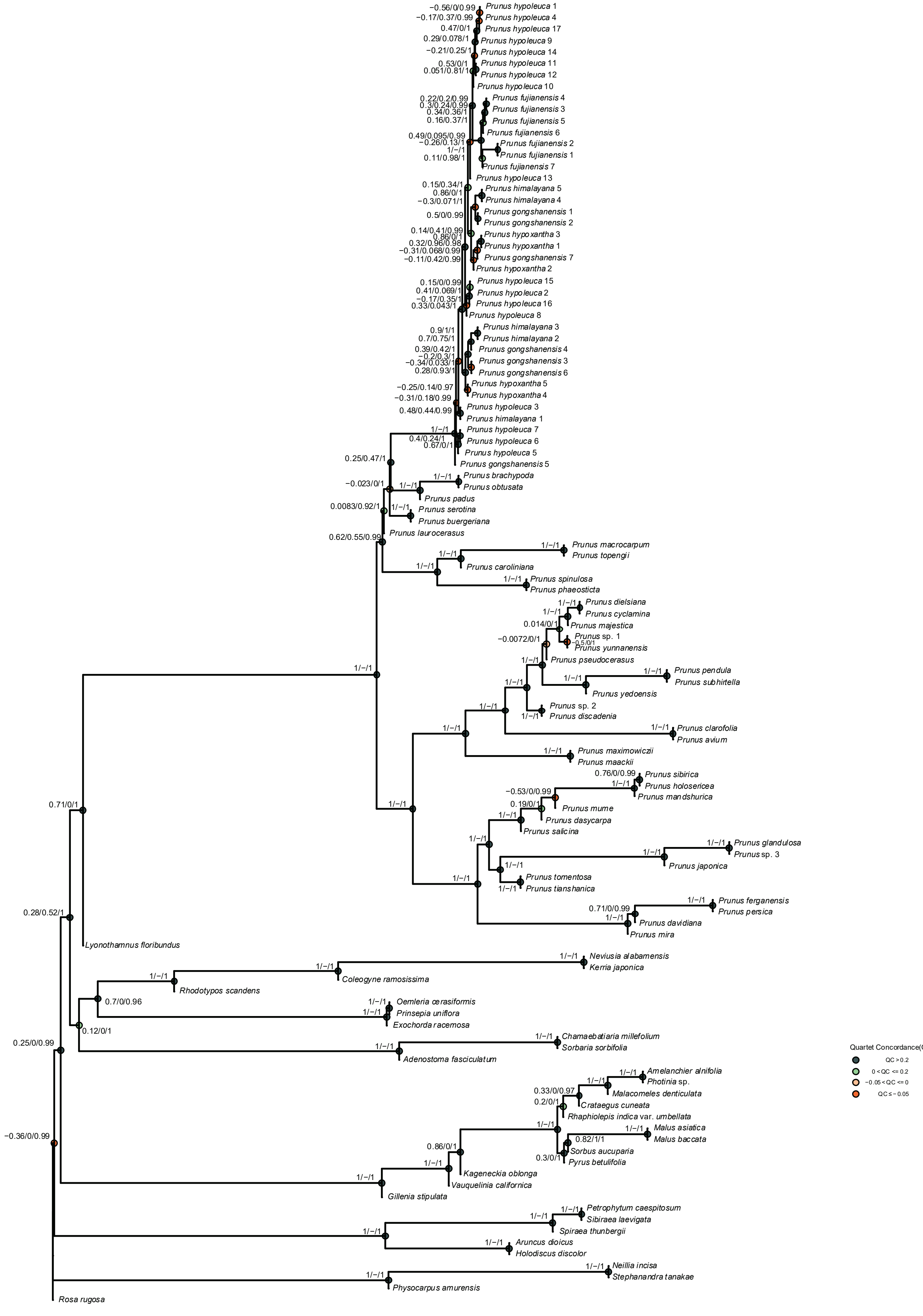

### Supplementary Fig. 15

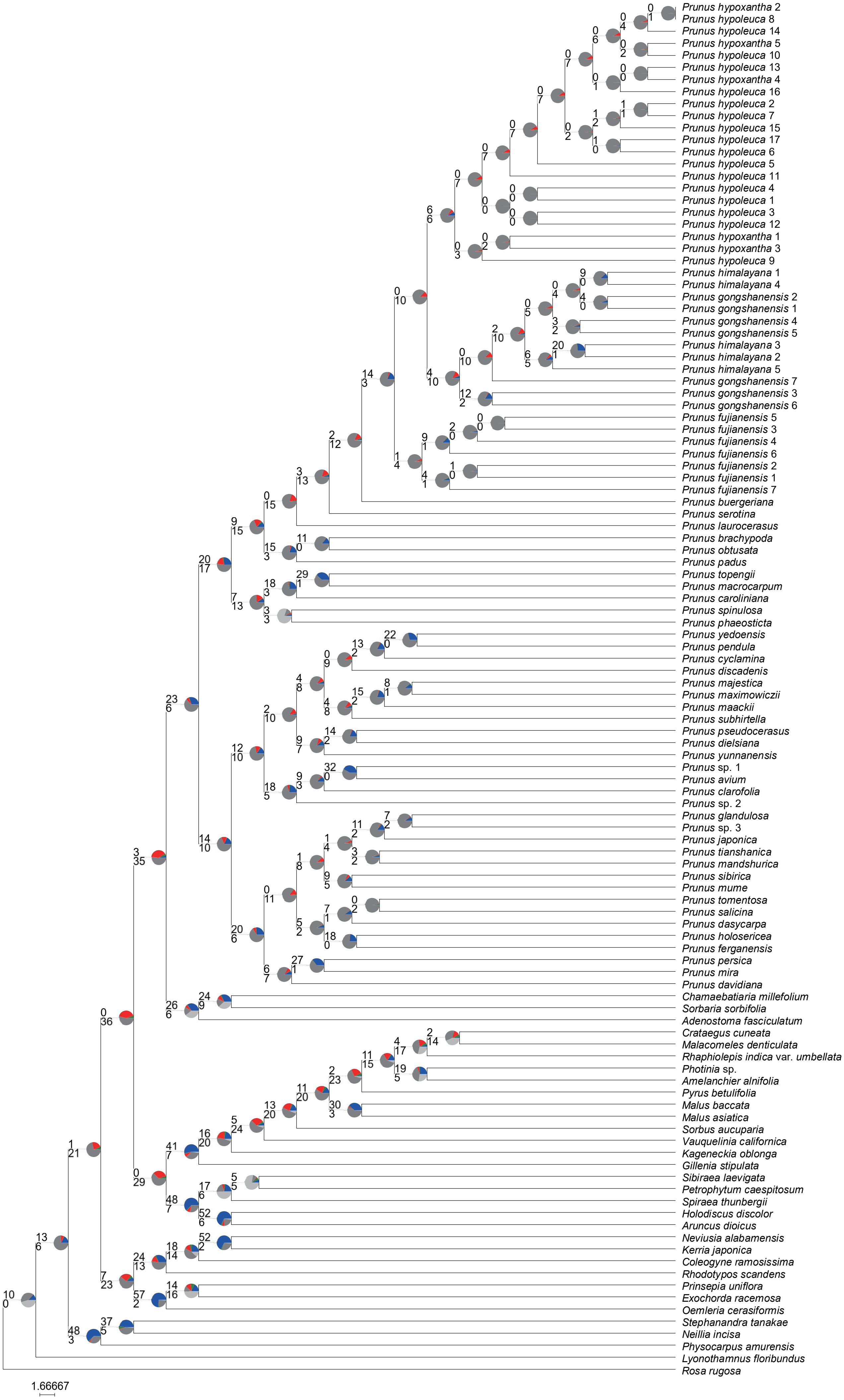

Supplementary Fig. 16

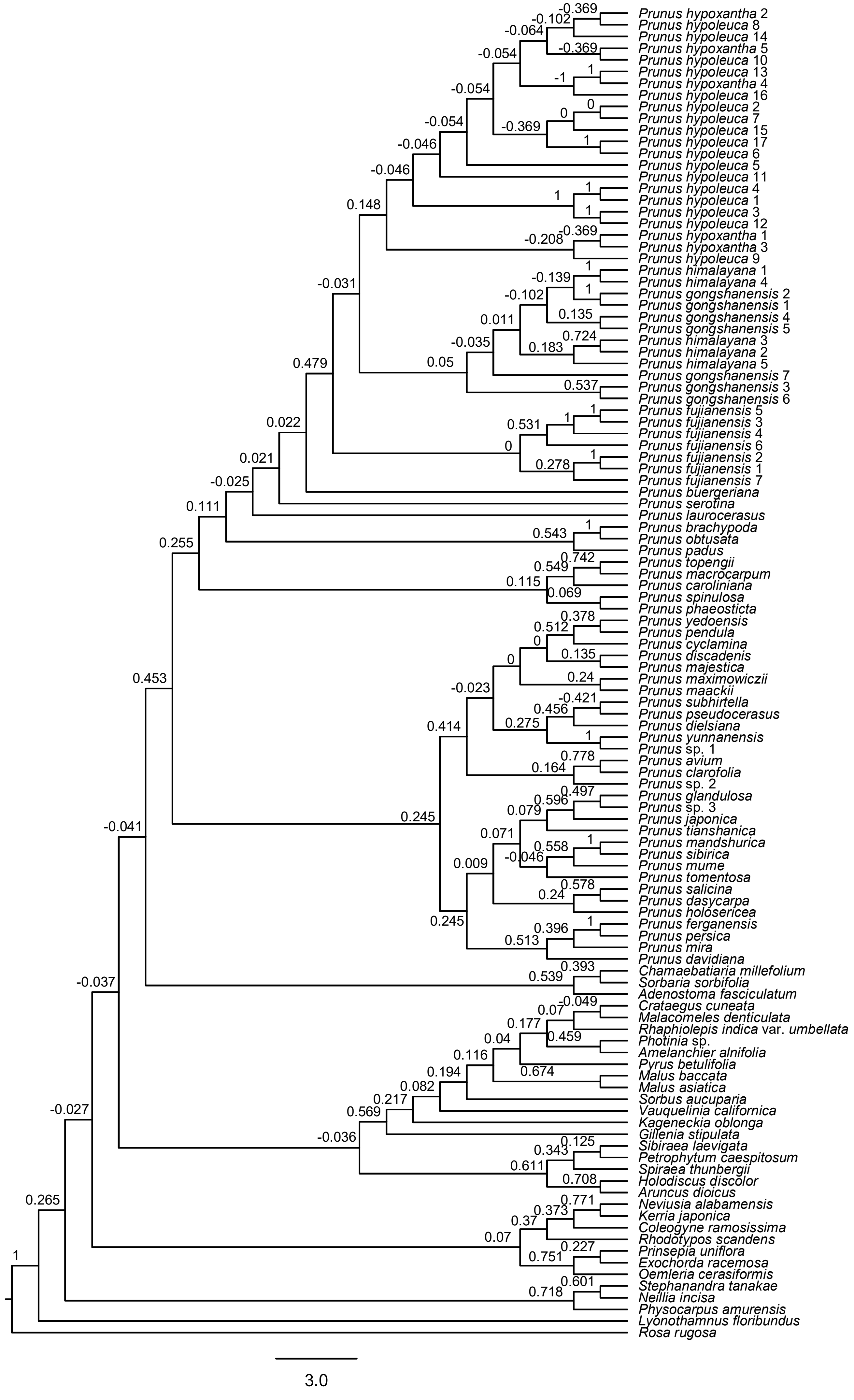

### Supplementary Fig. 17

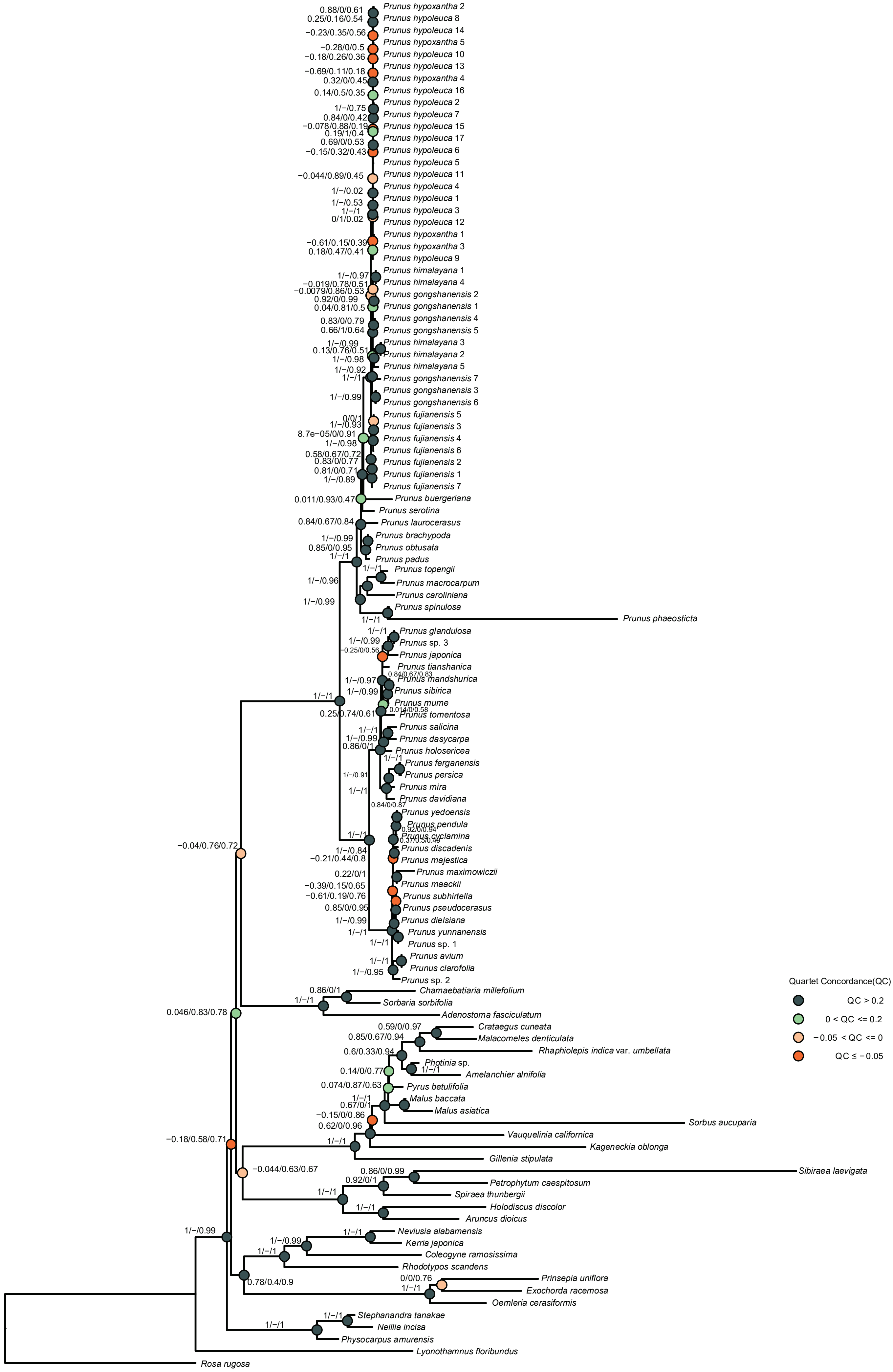

### Supplementary Fig. 18

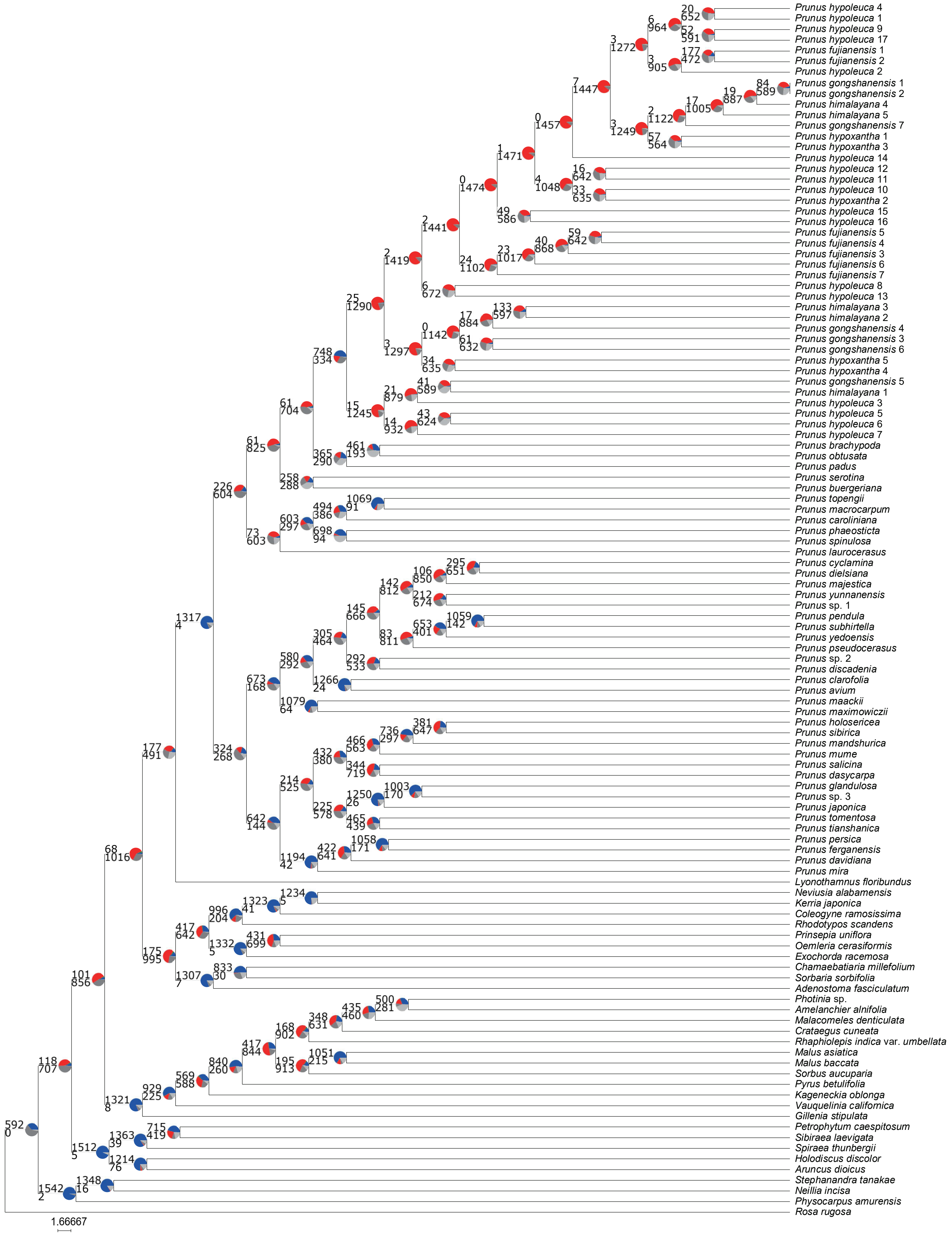

Supplementary Fig. 19

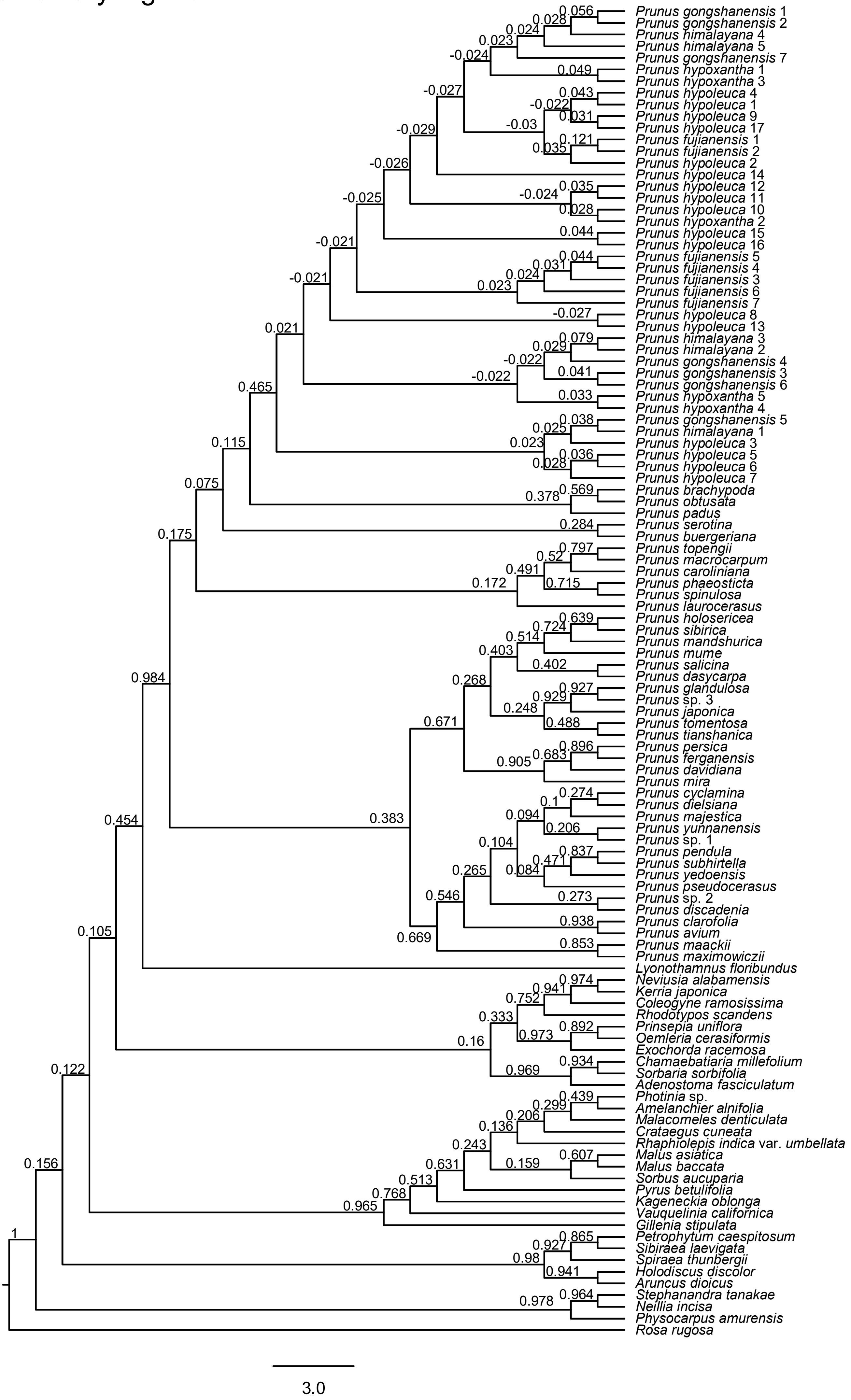

### Supplementary Fig. 20

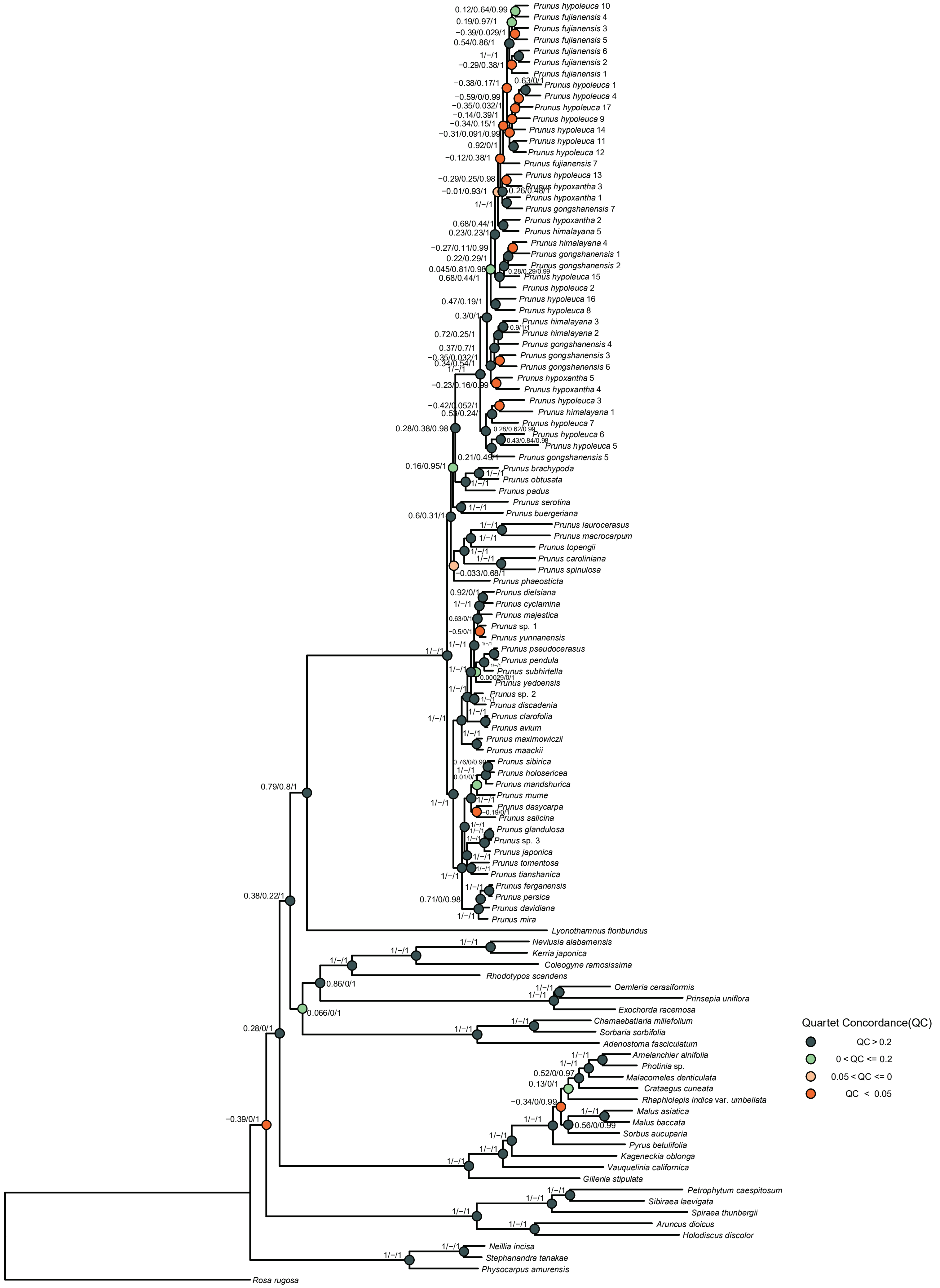

### Supplementary Fig. 21

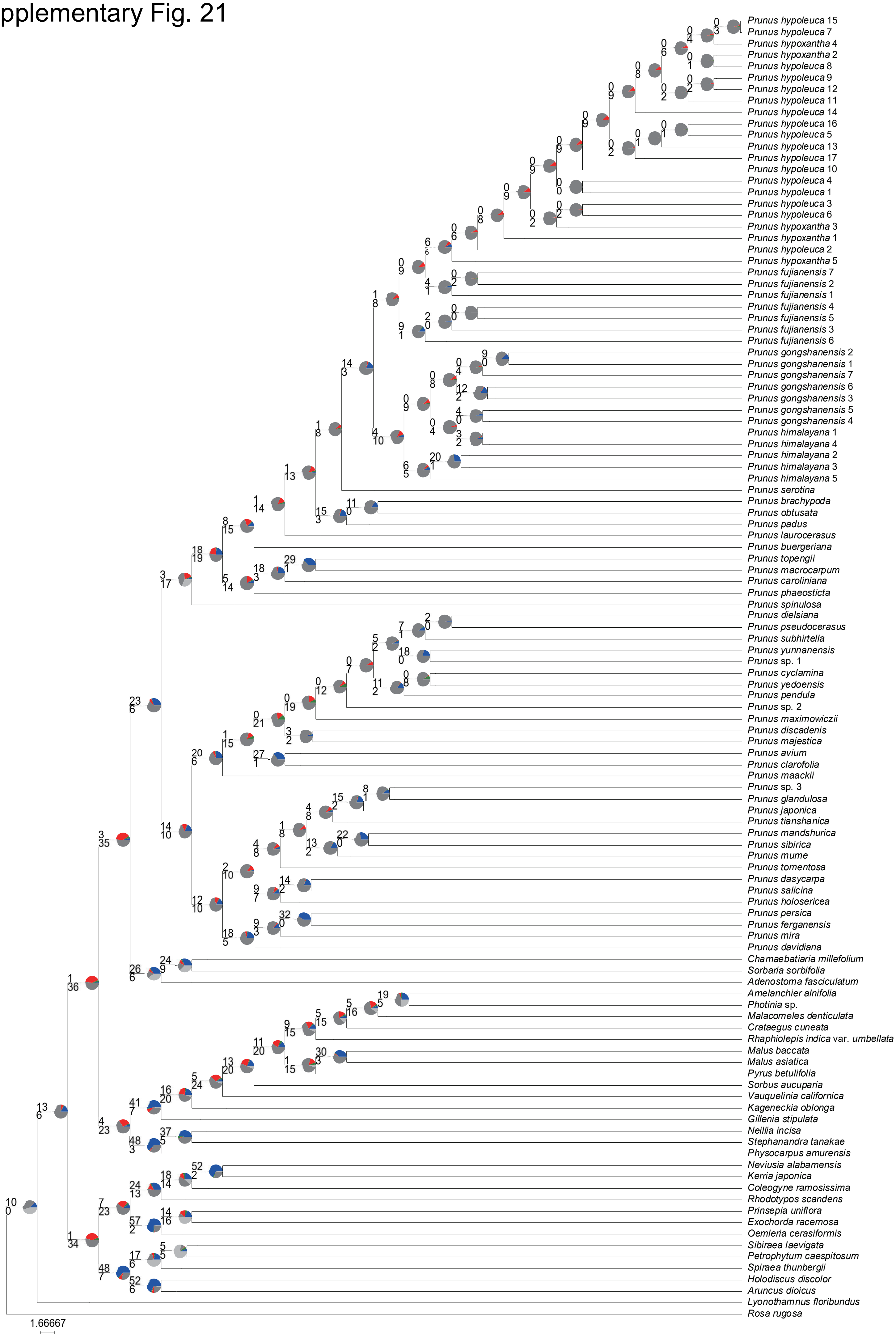

### Supplementary Fig. 22

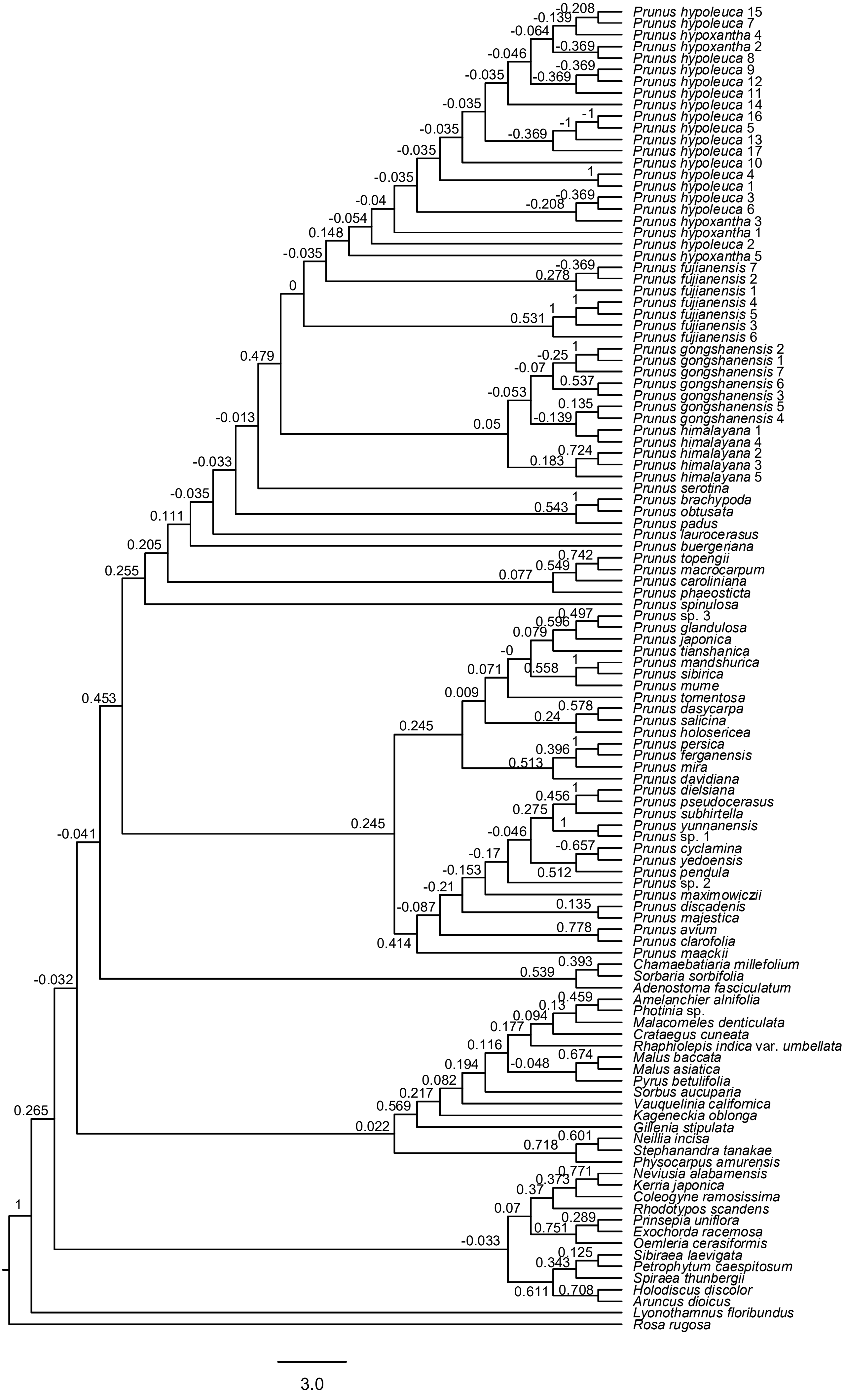

### Supplementary Fig. 23

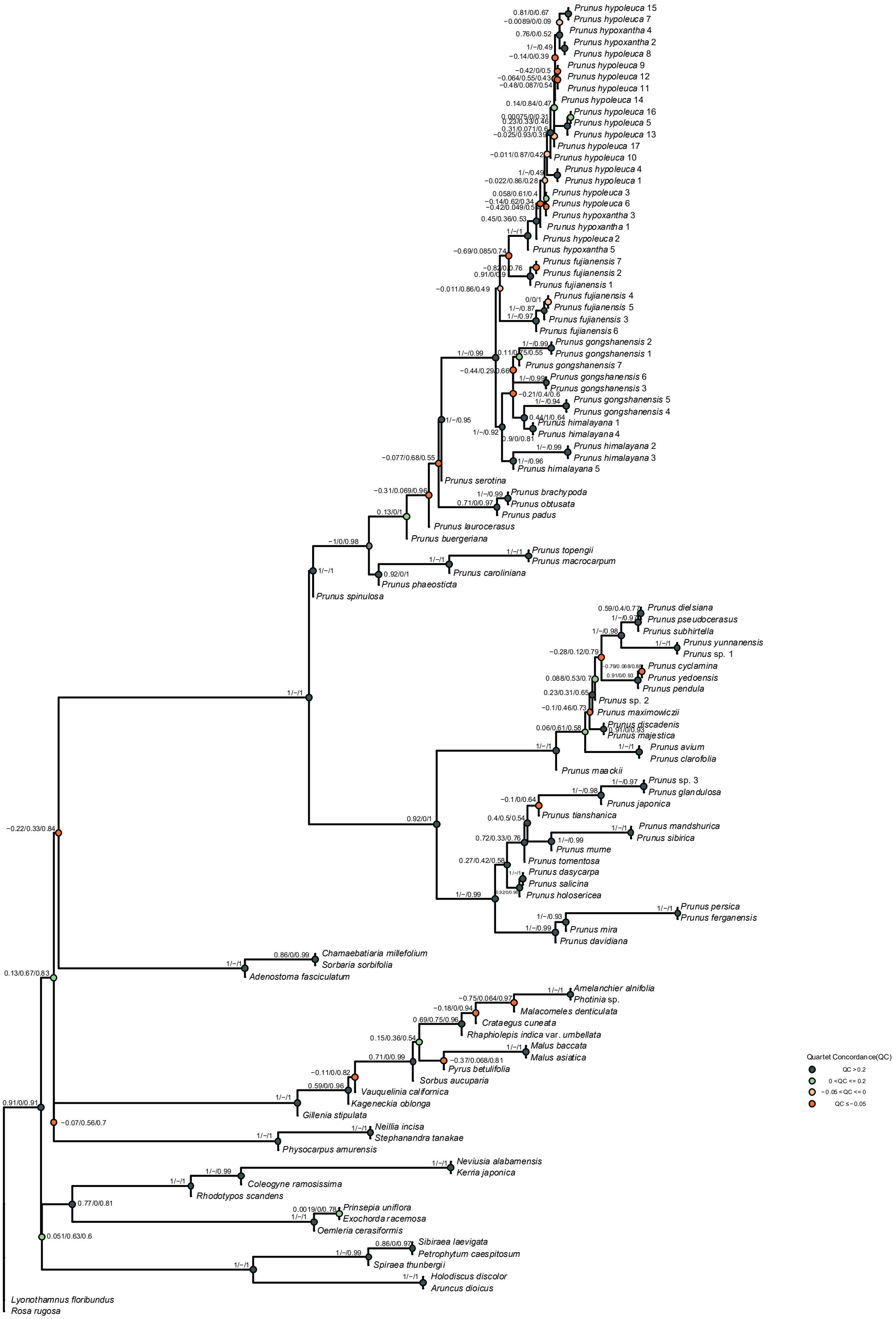

Supplementary Fig. 24

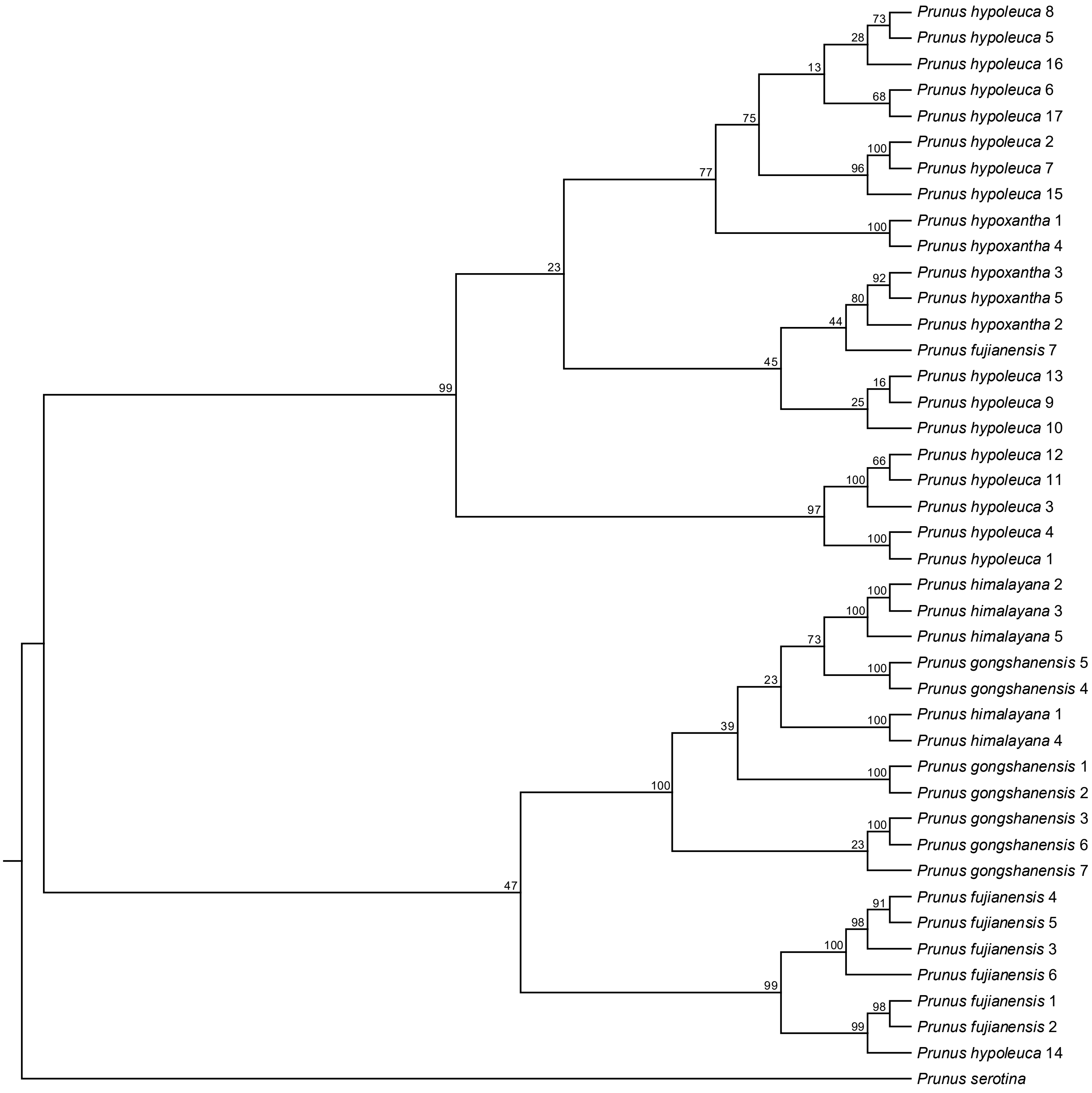

### Supplementary Fig. 25

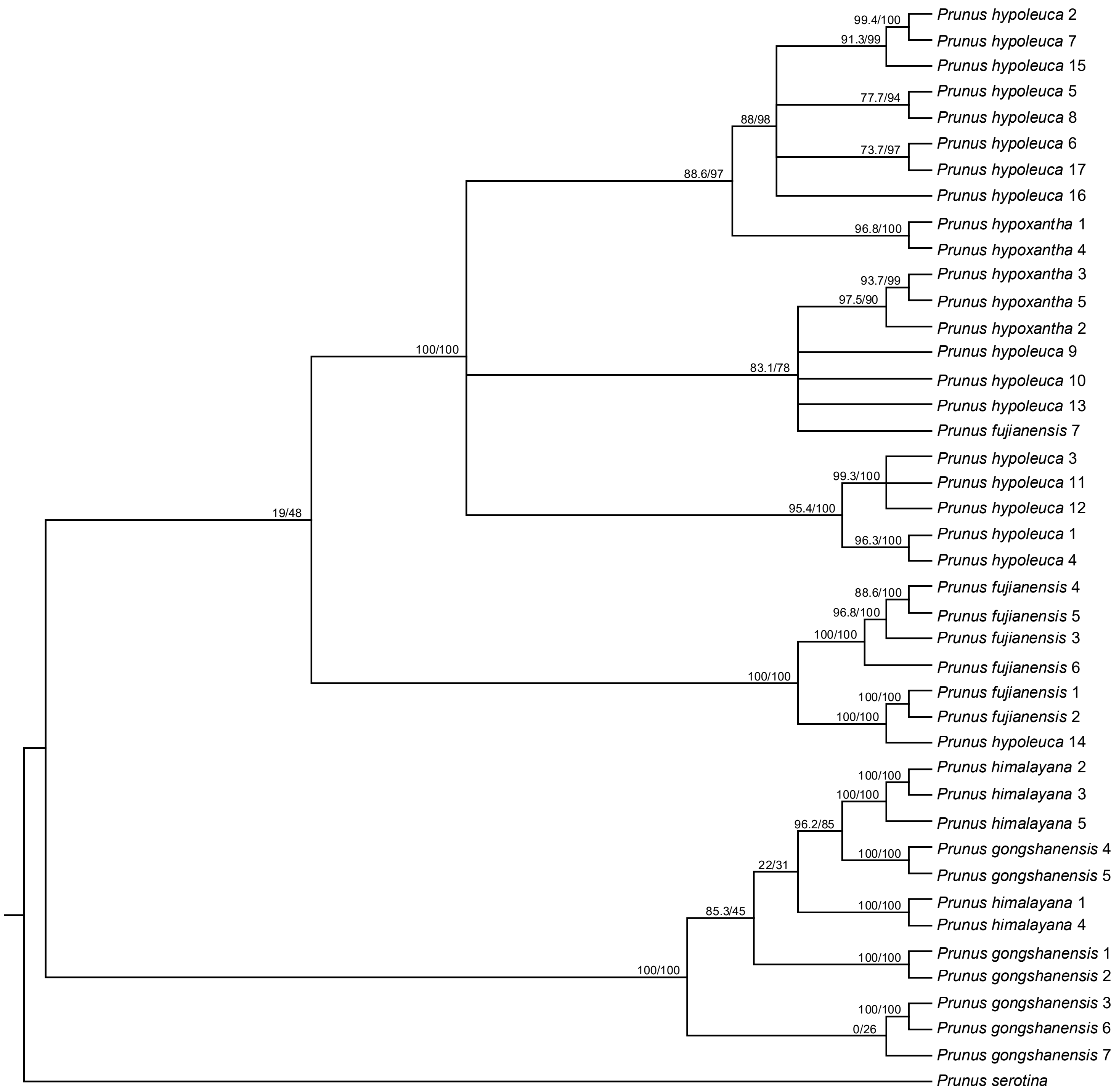

Supplementary Fig. 26·

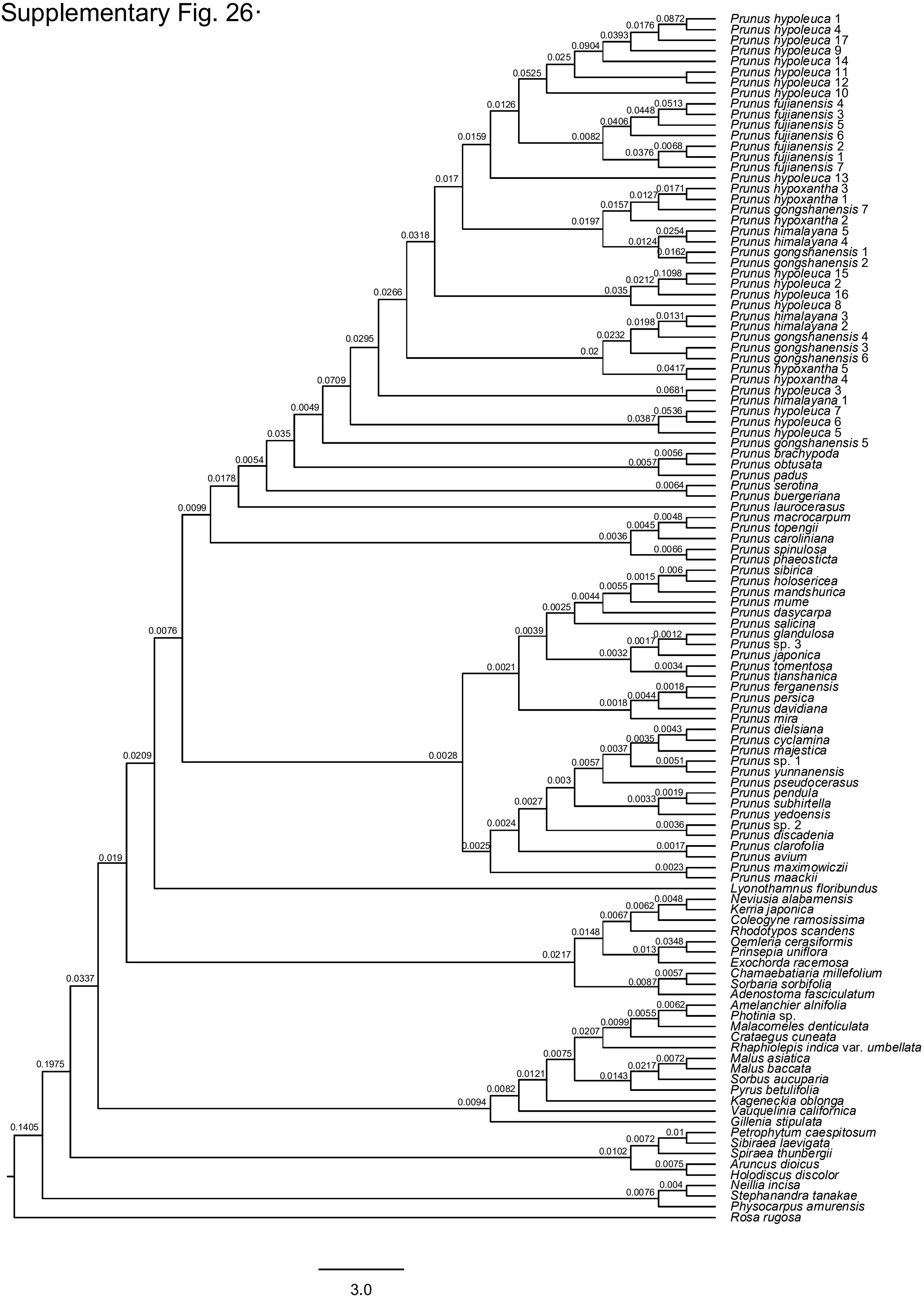

Supplementary Fig. 27

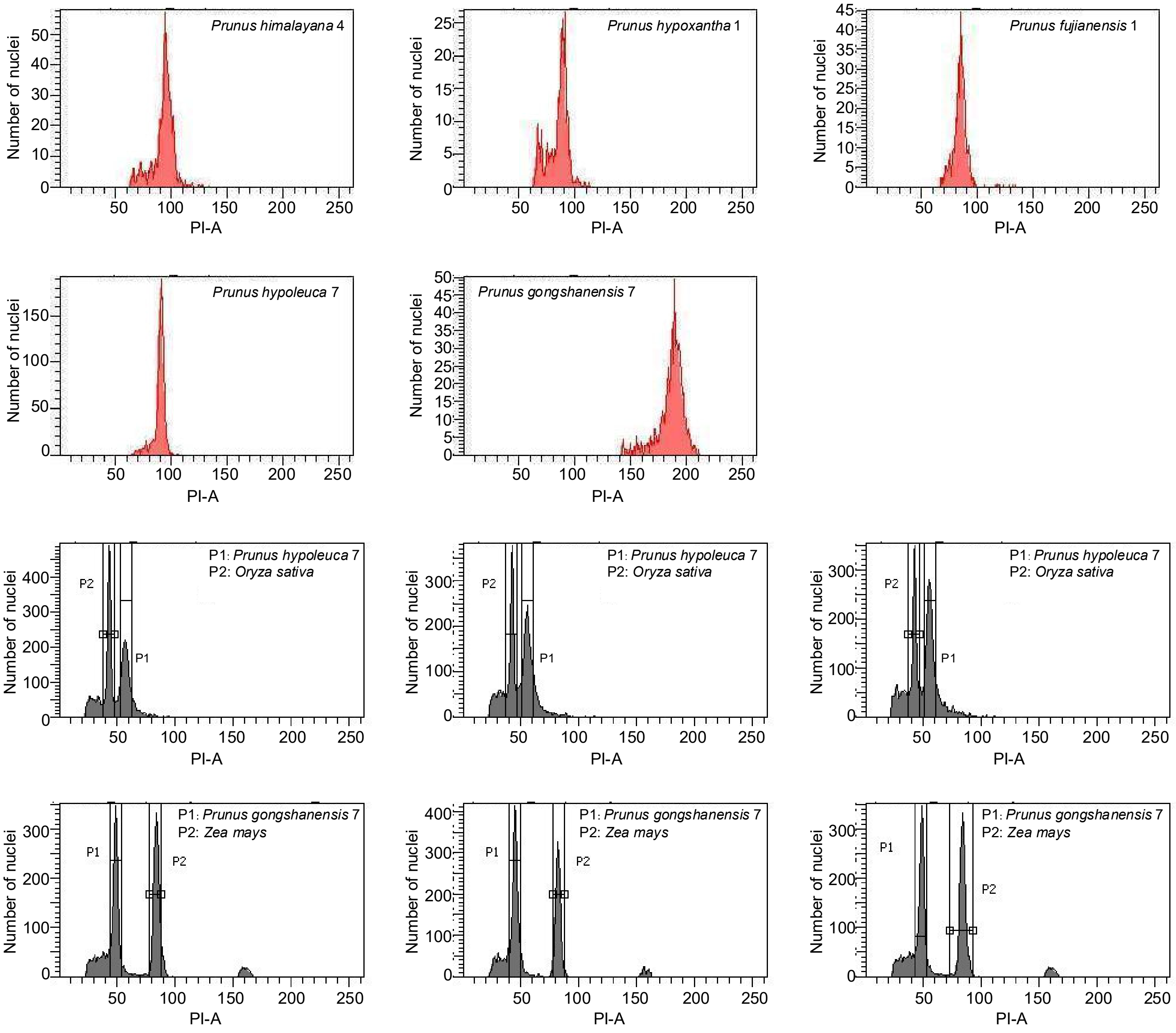

### Supplementary Fig. 28

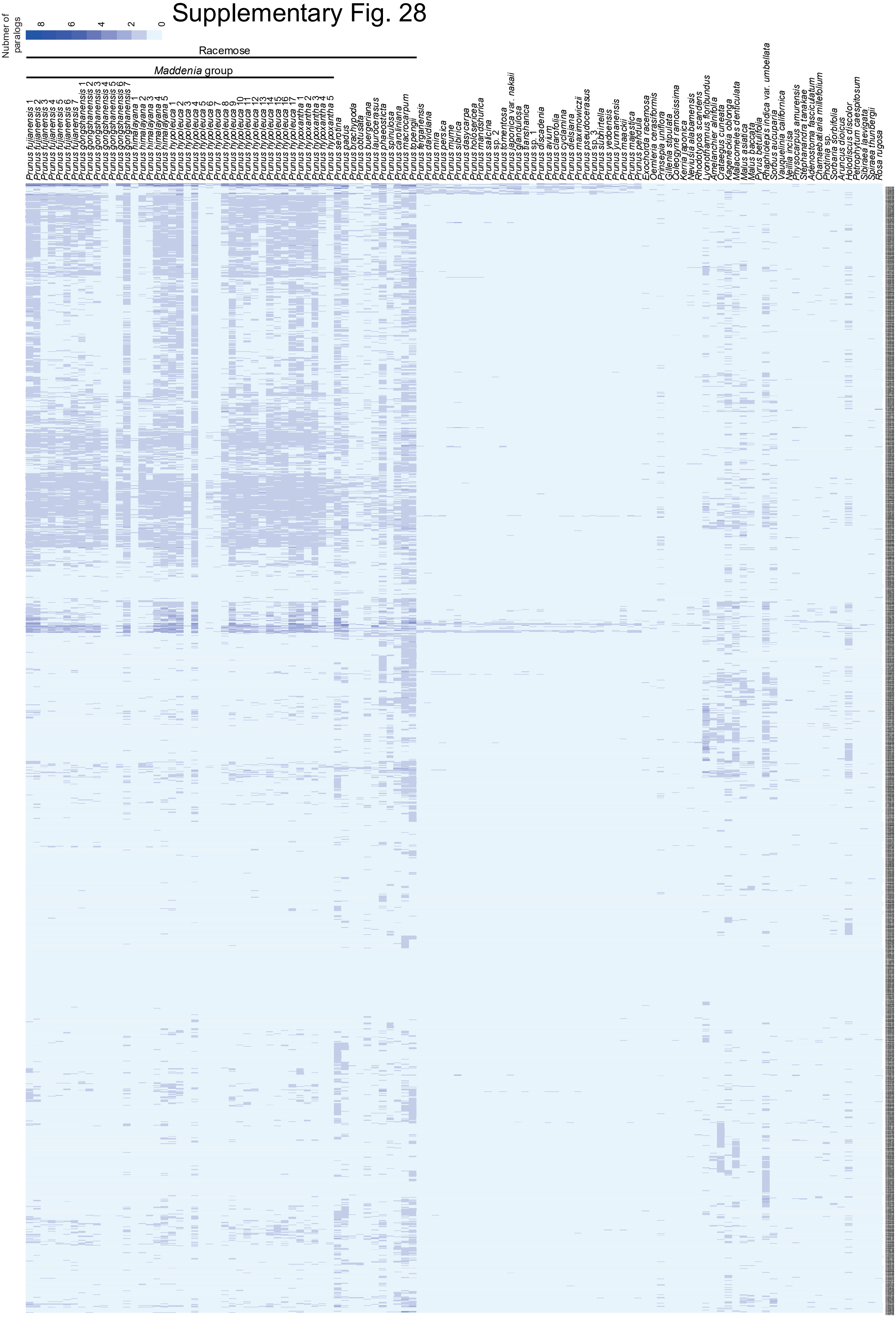

### Supplementary Fig. 29

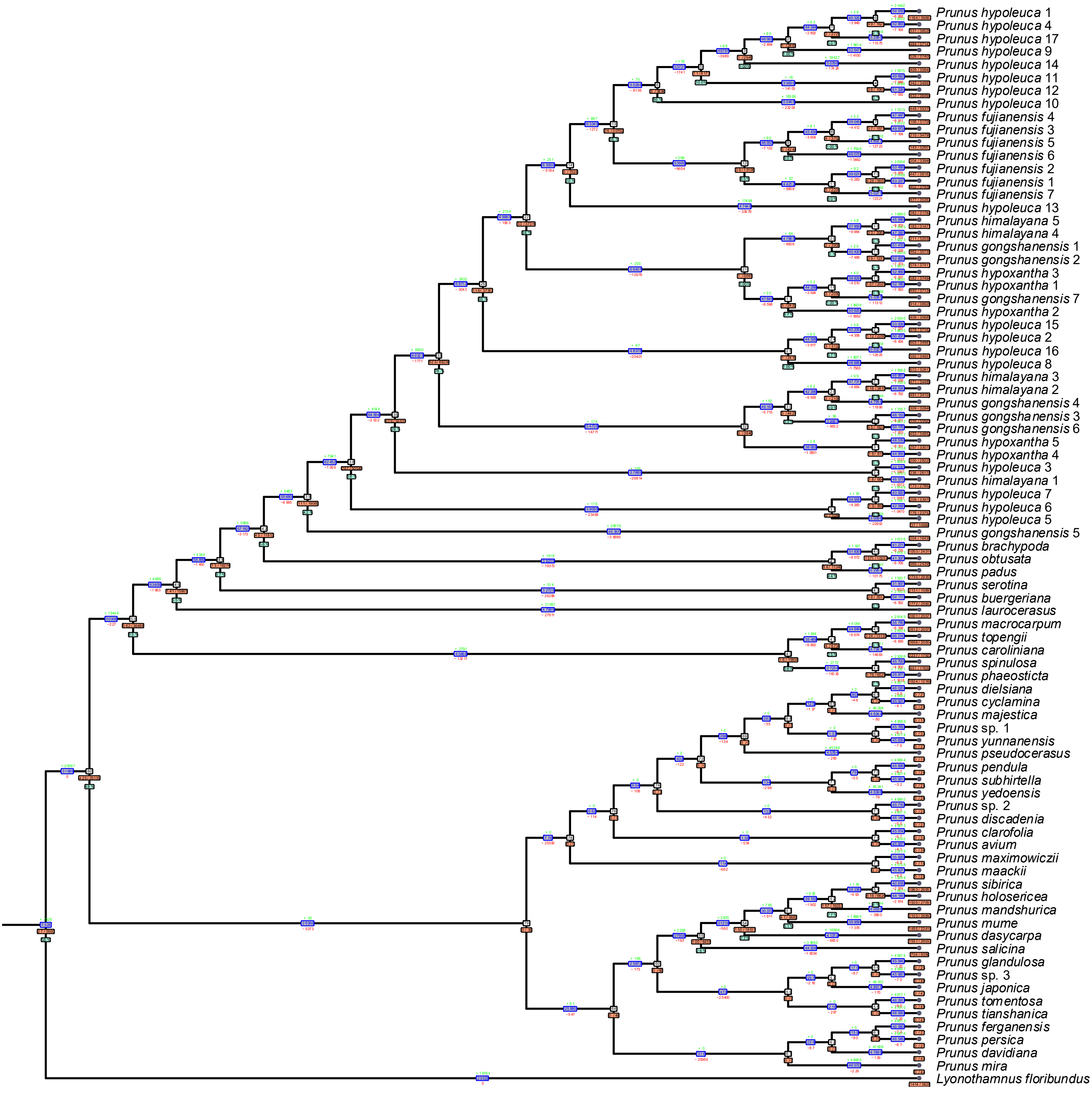

### Supplementary Fig. 30

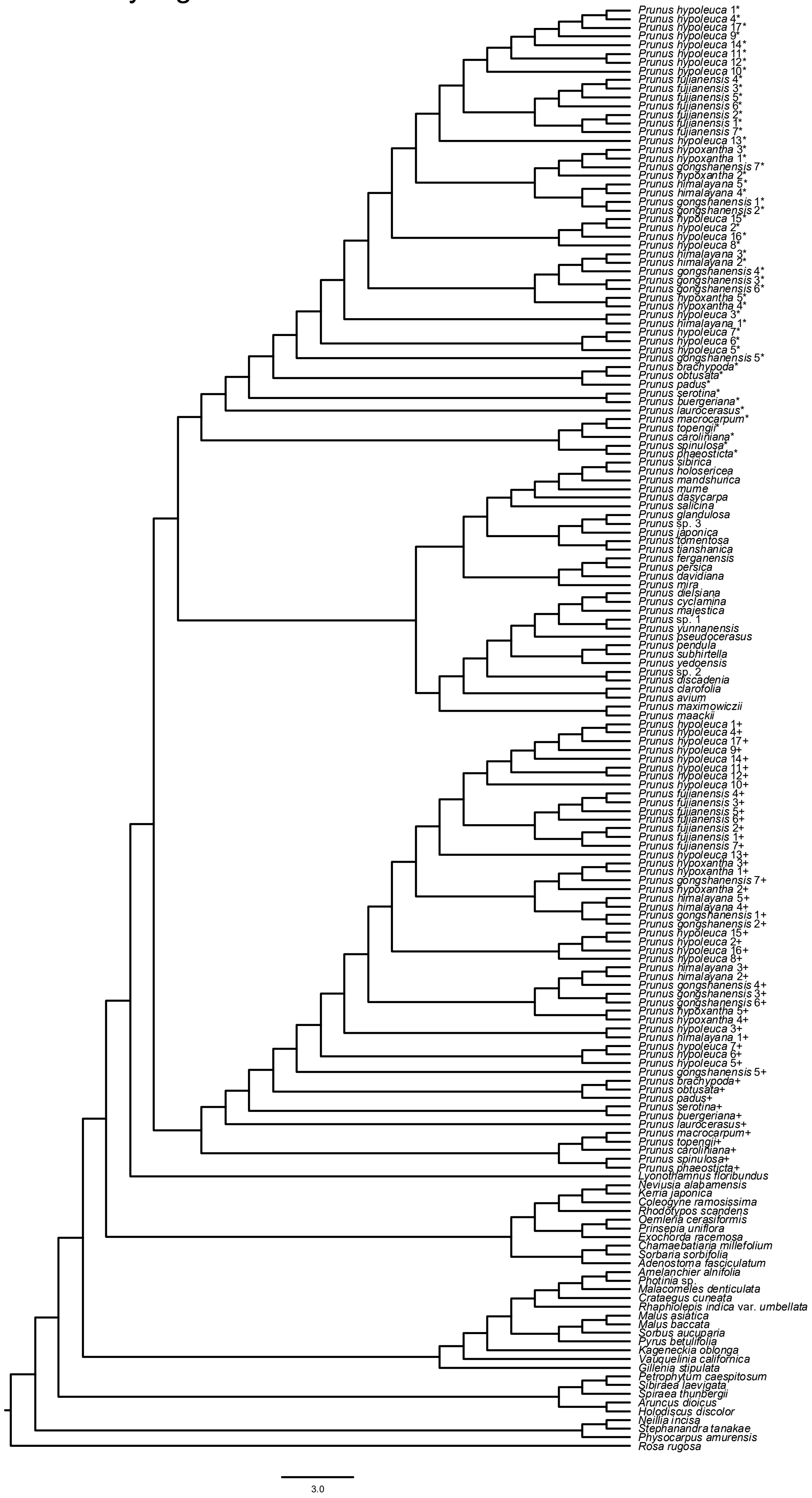

Supplementary Fig. 31

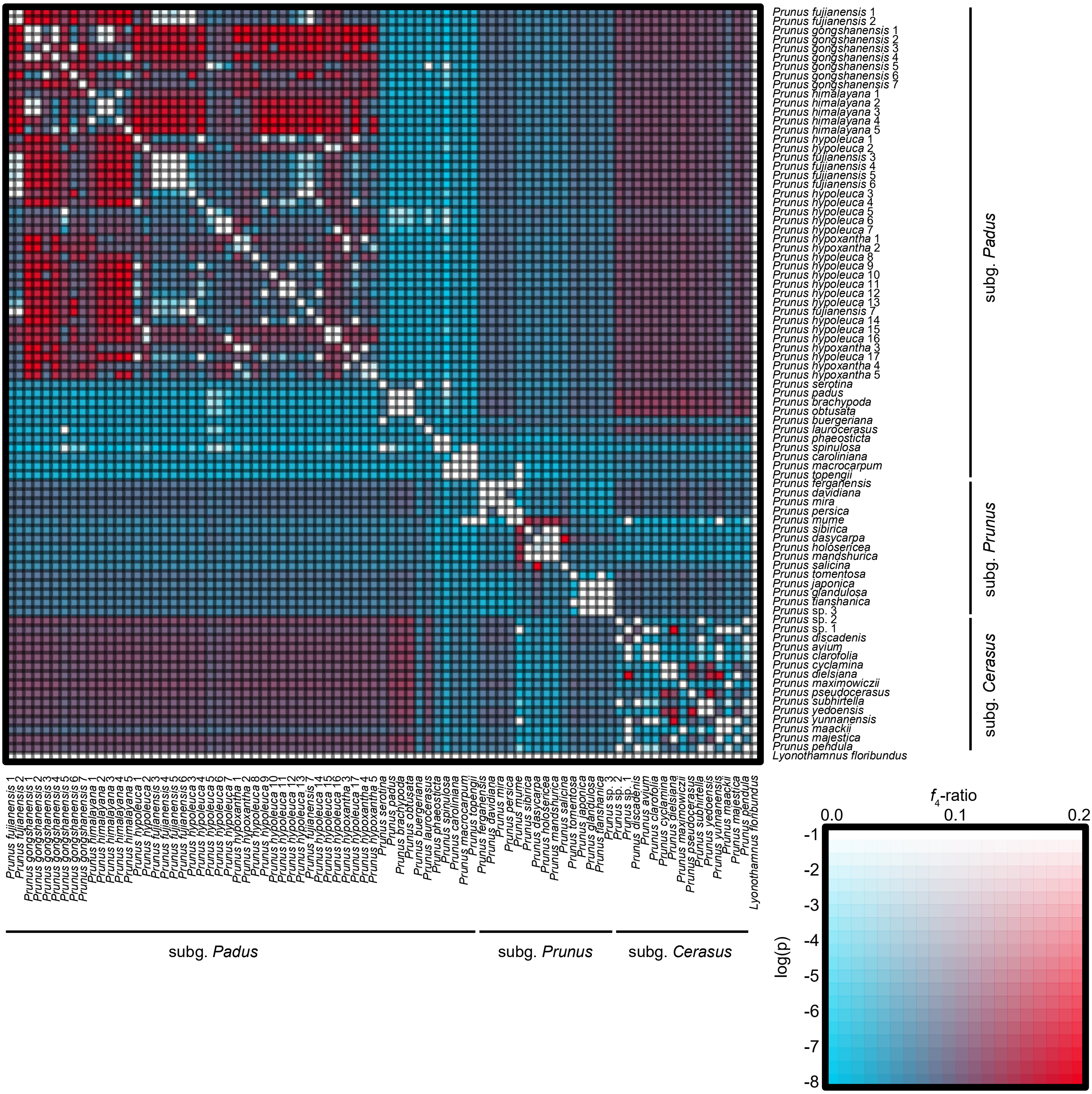

Supplementary Fig. 32

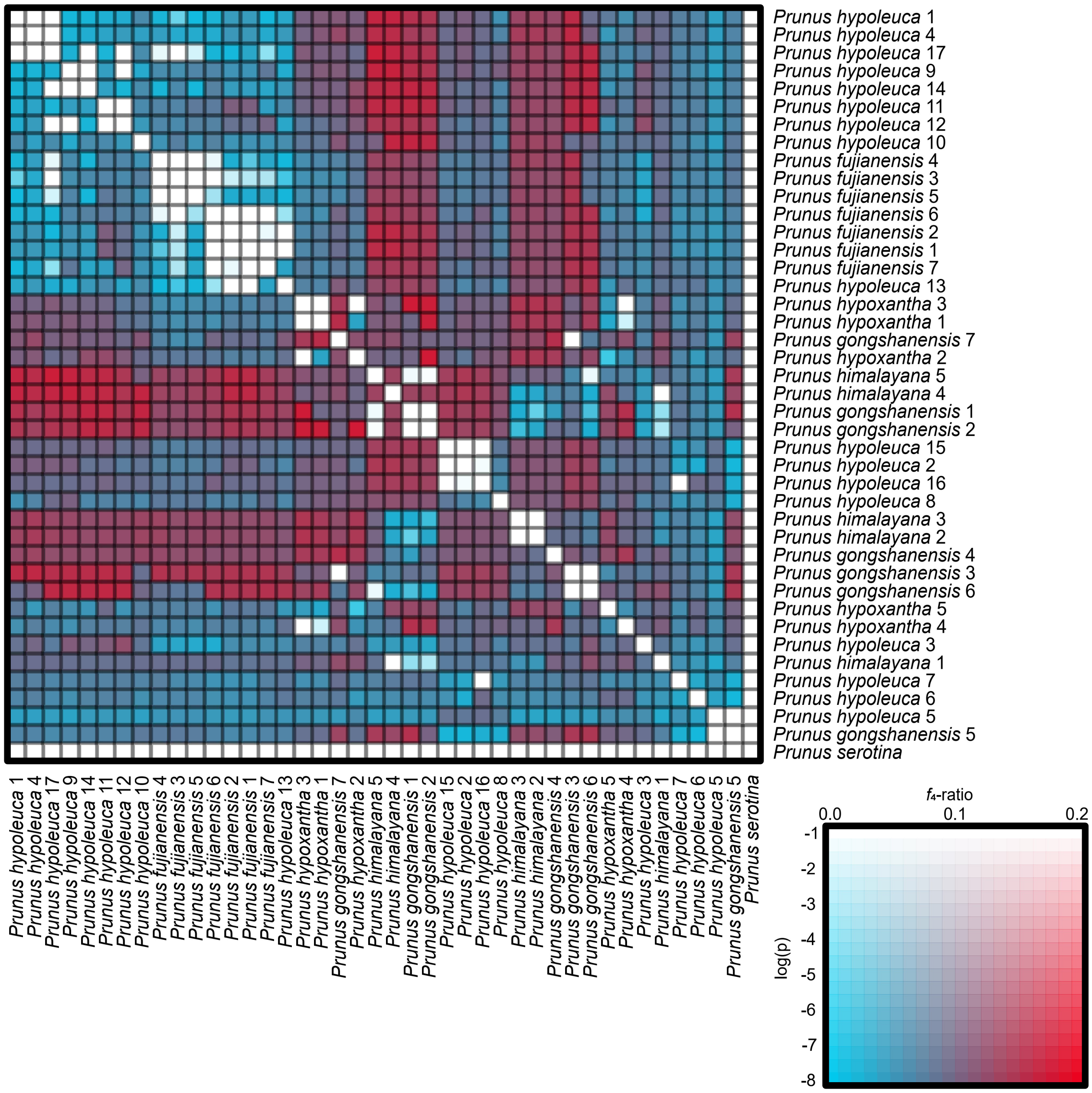

Supplementary Fig. 33

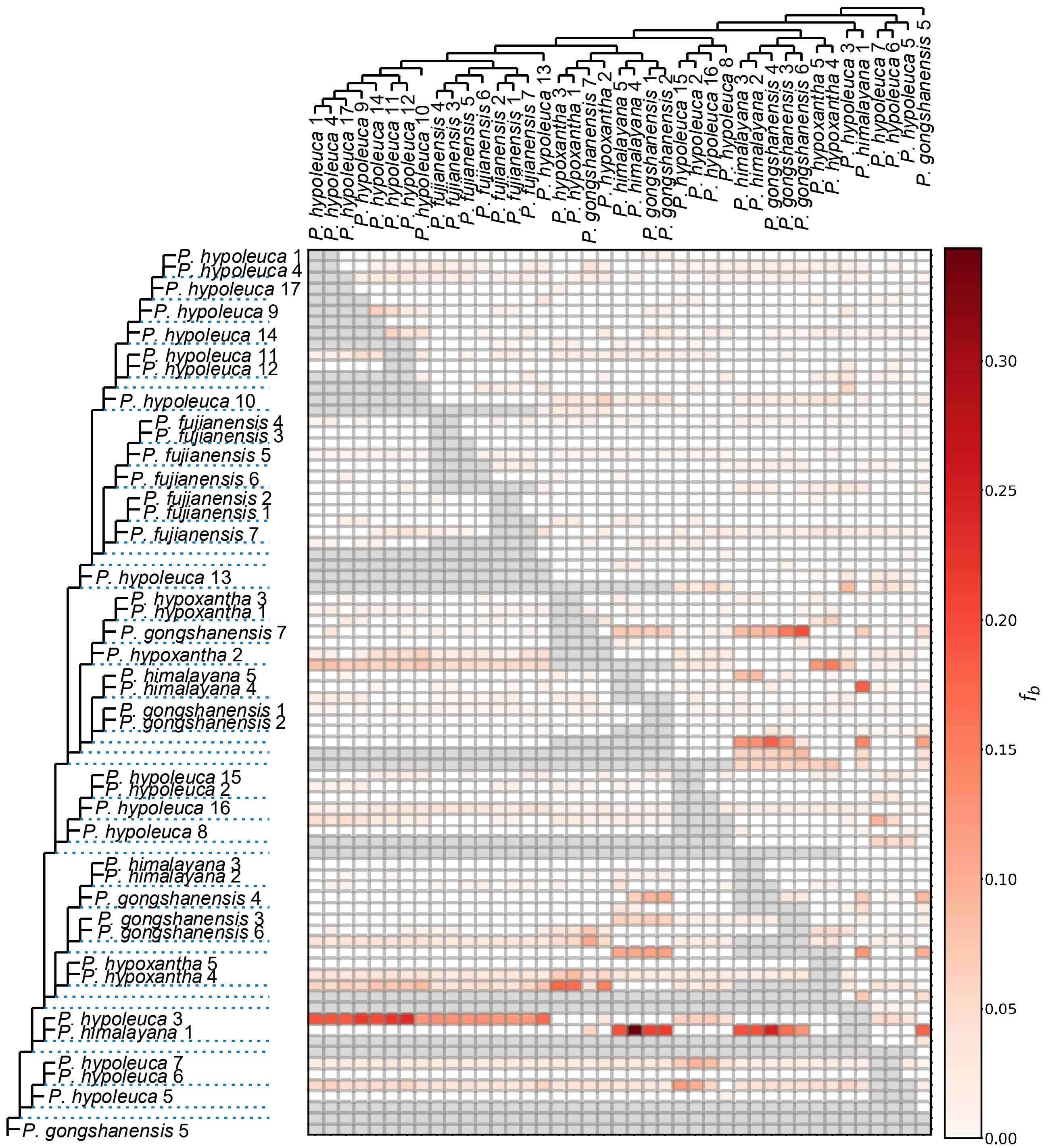

### Supplementary Fig. 34

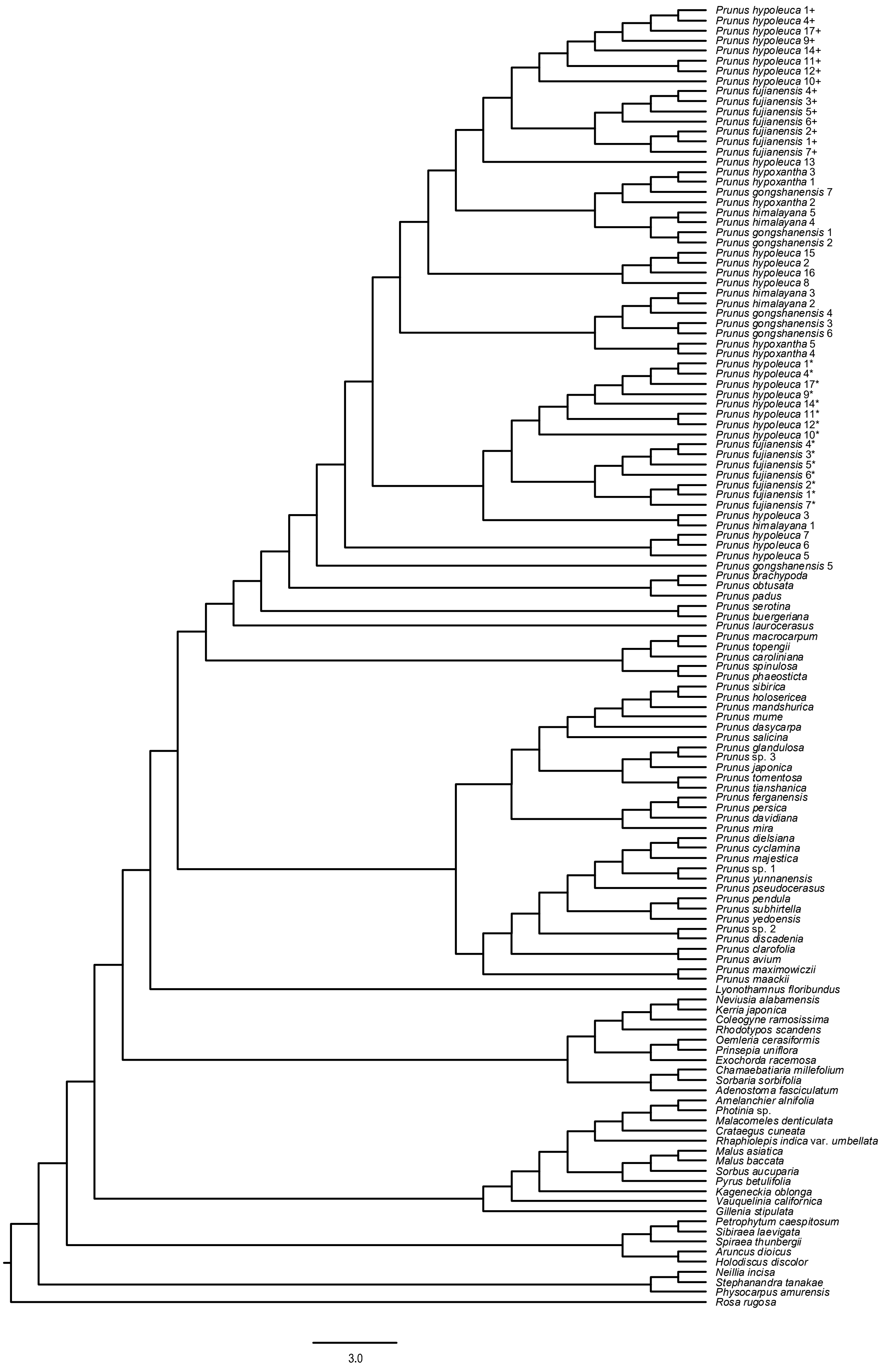

### Supplementary Fig. 35

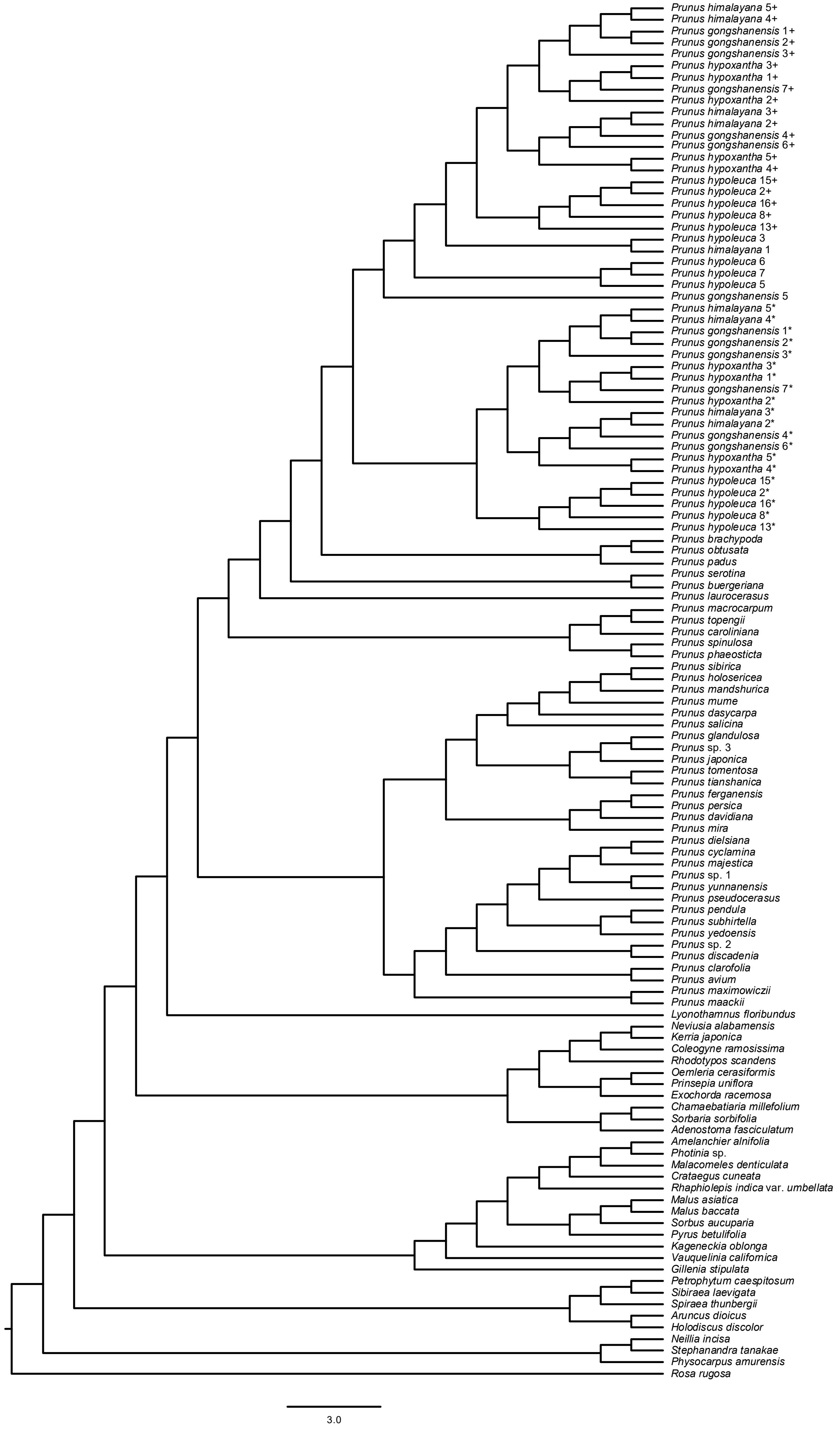

### Supplementary Fig. 36

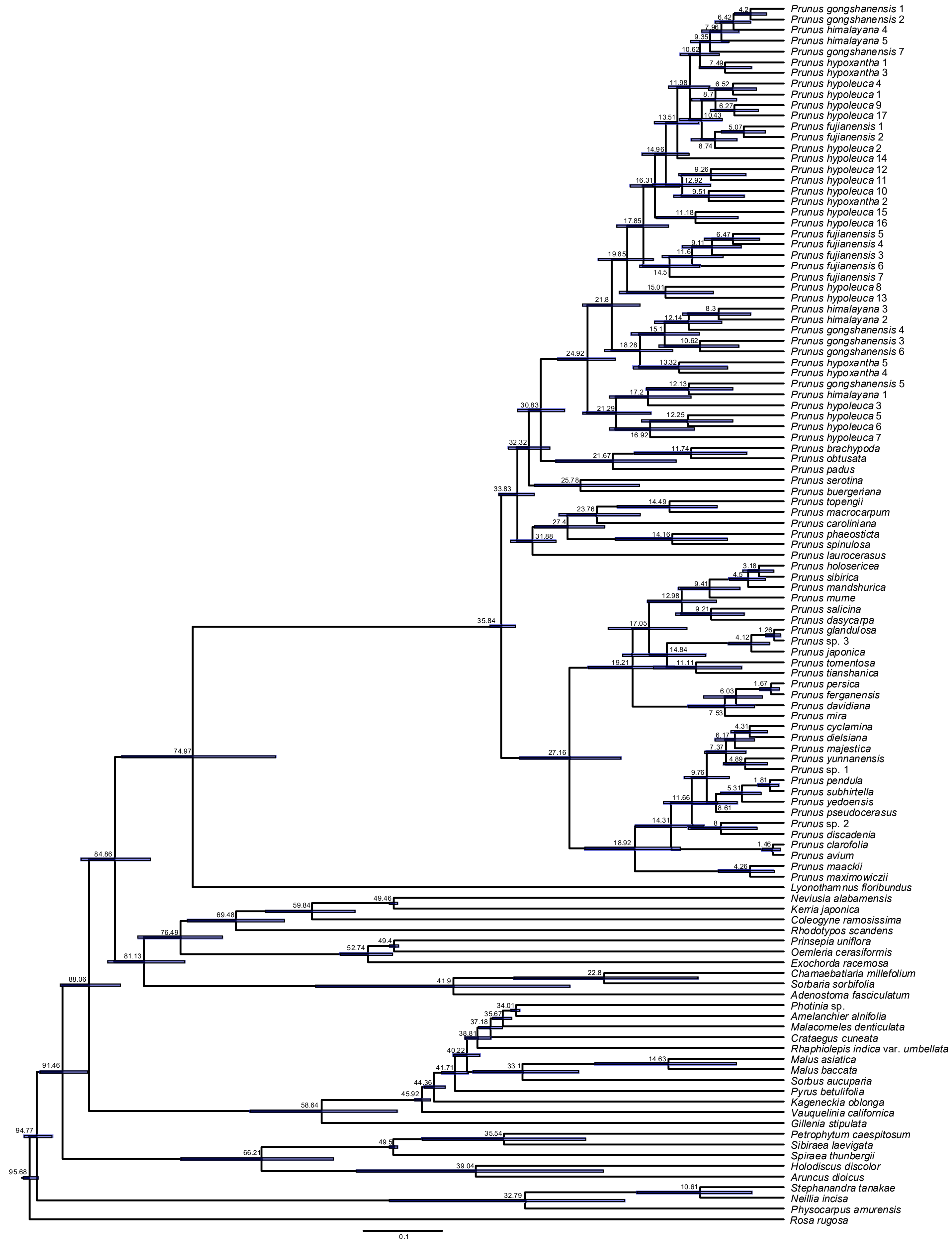

### Supplementary Fig. 37

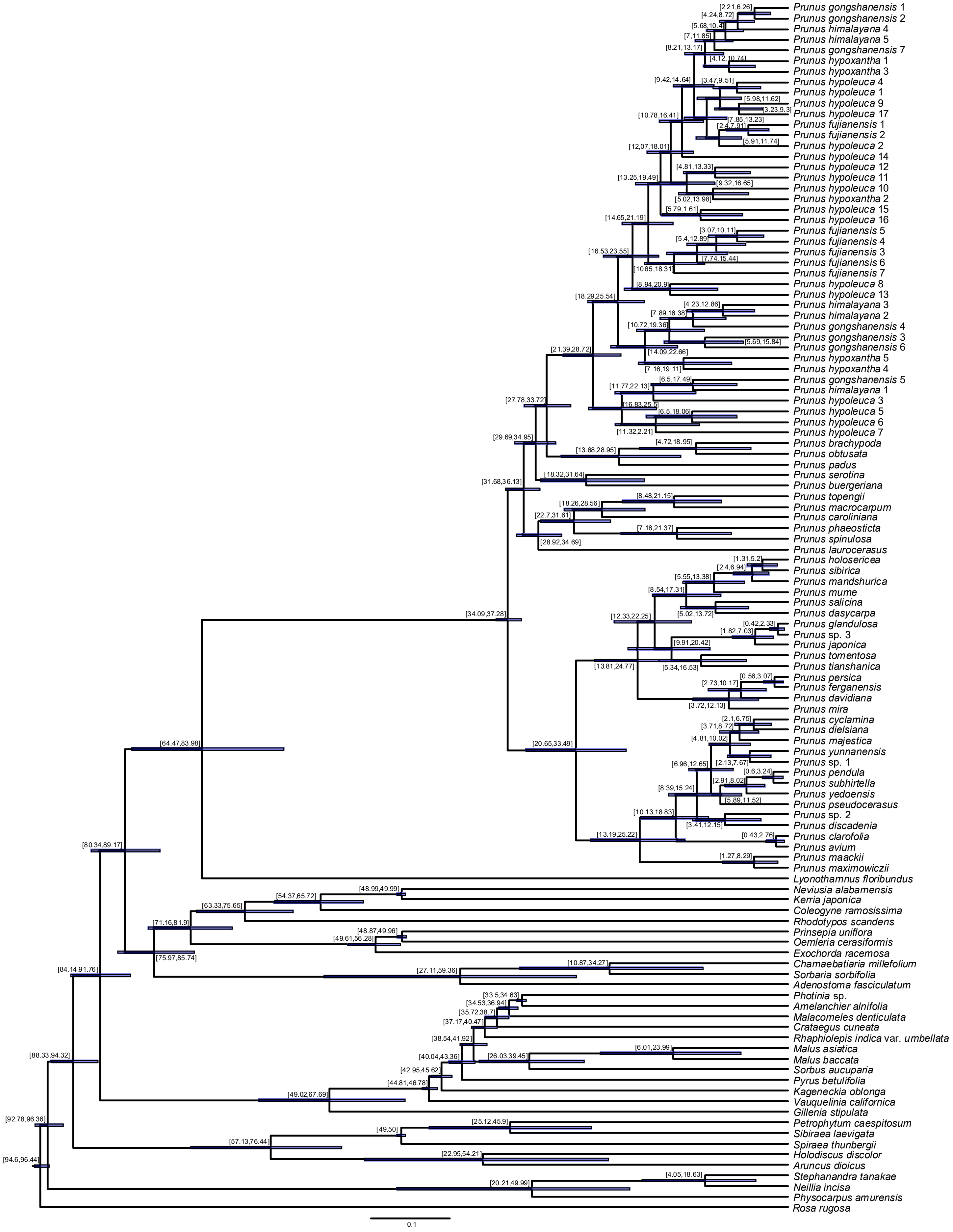

### Supplementary Fig. 38

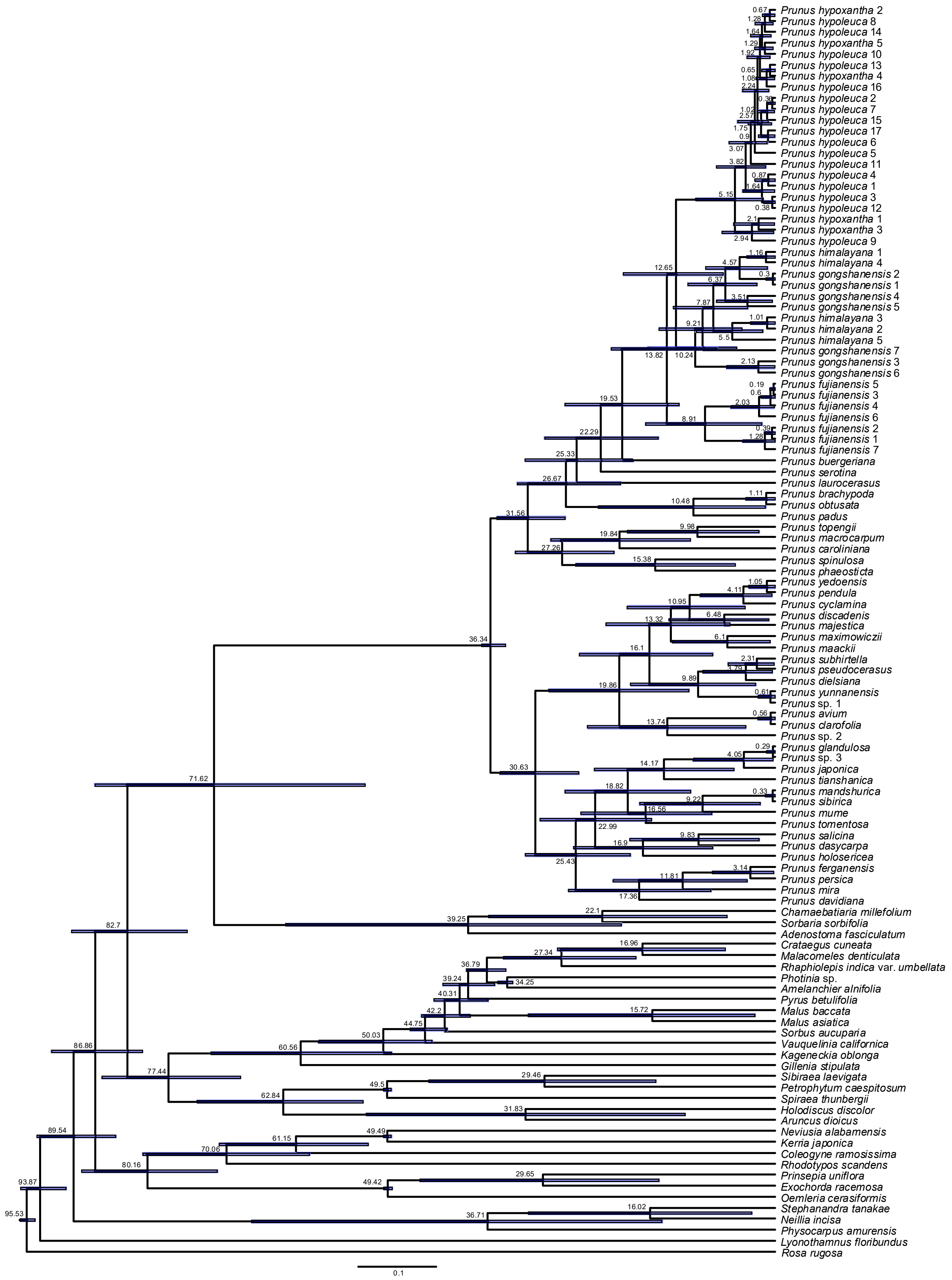

Supplementary Fig. 39

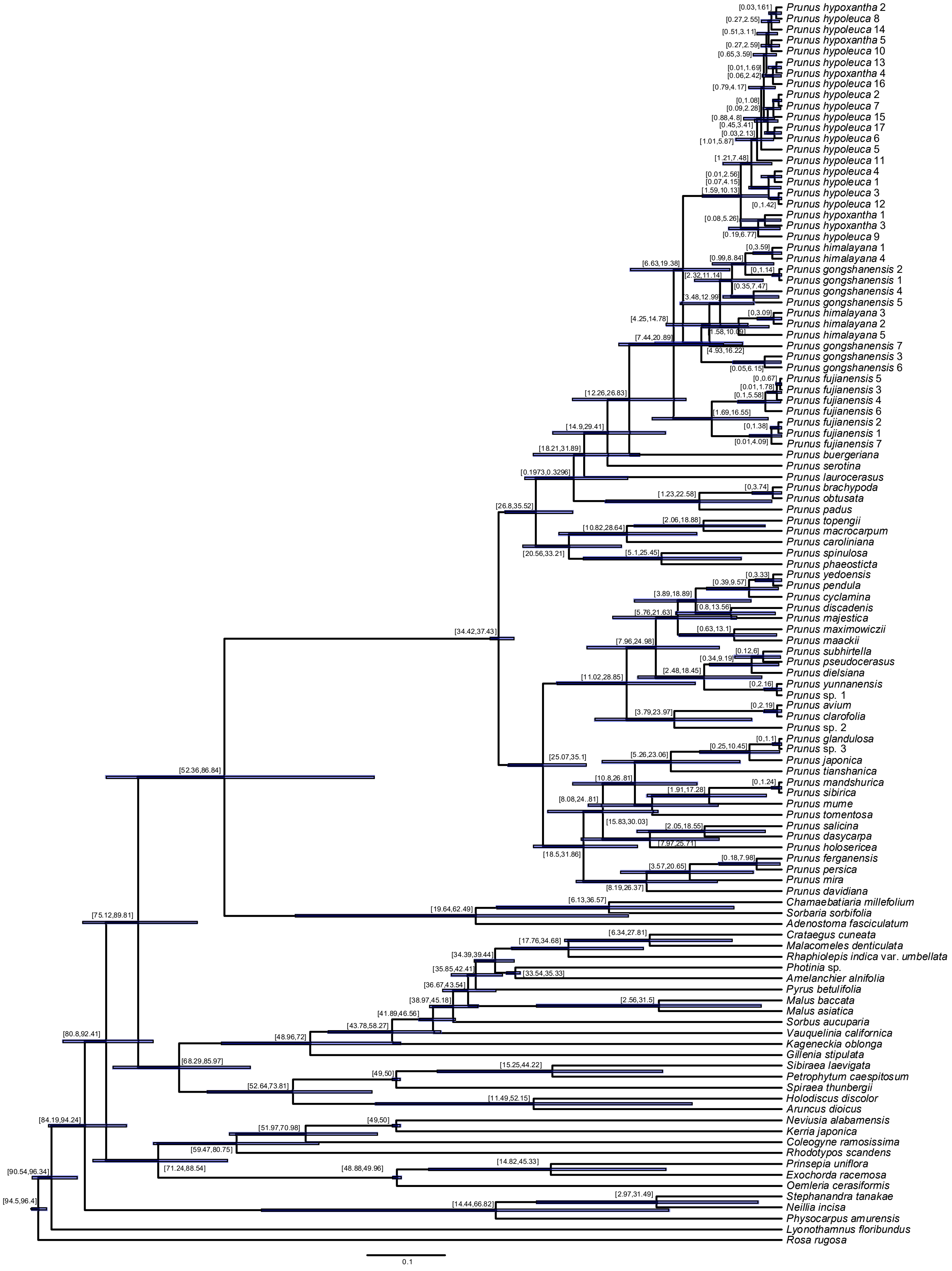

### Supplementary Fig. 40

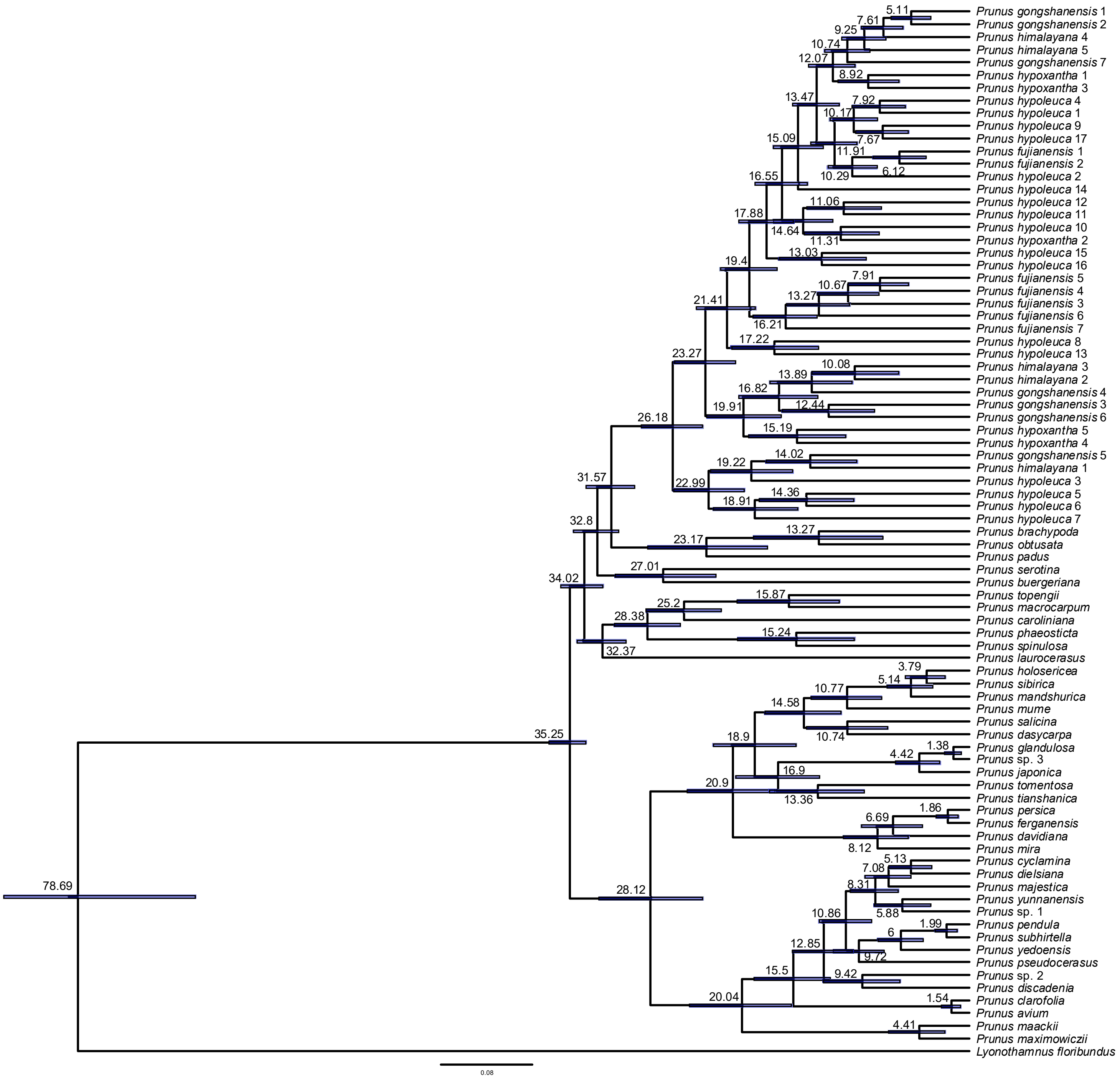

Supplementary Fig. 41

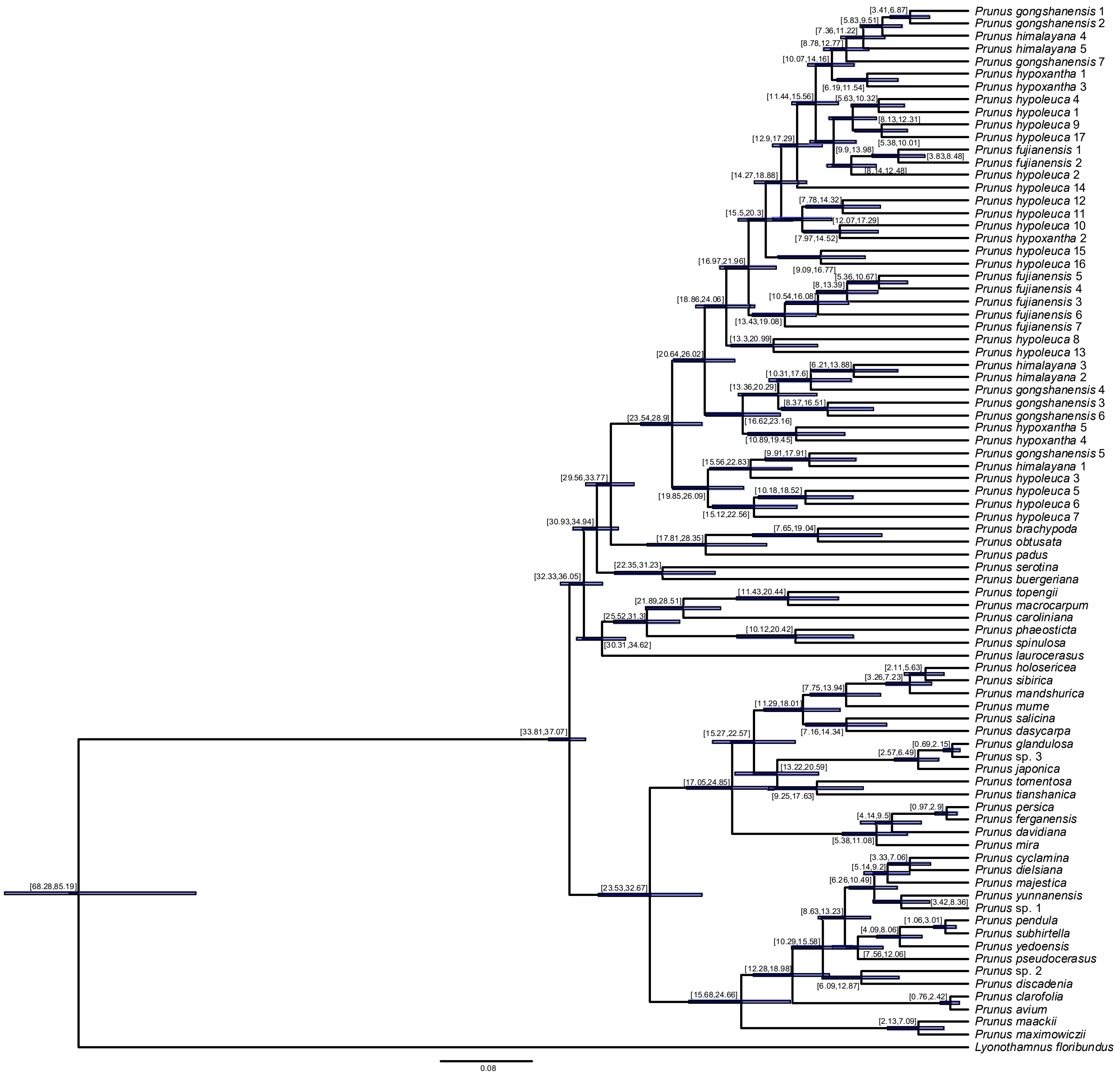

### Supplementary Fig. 42

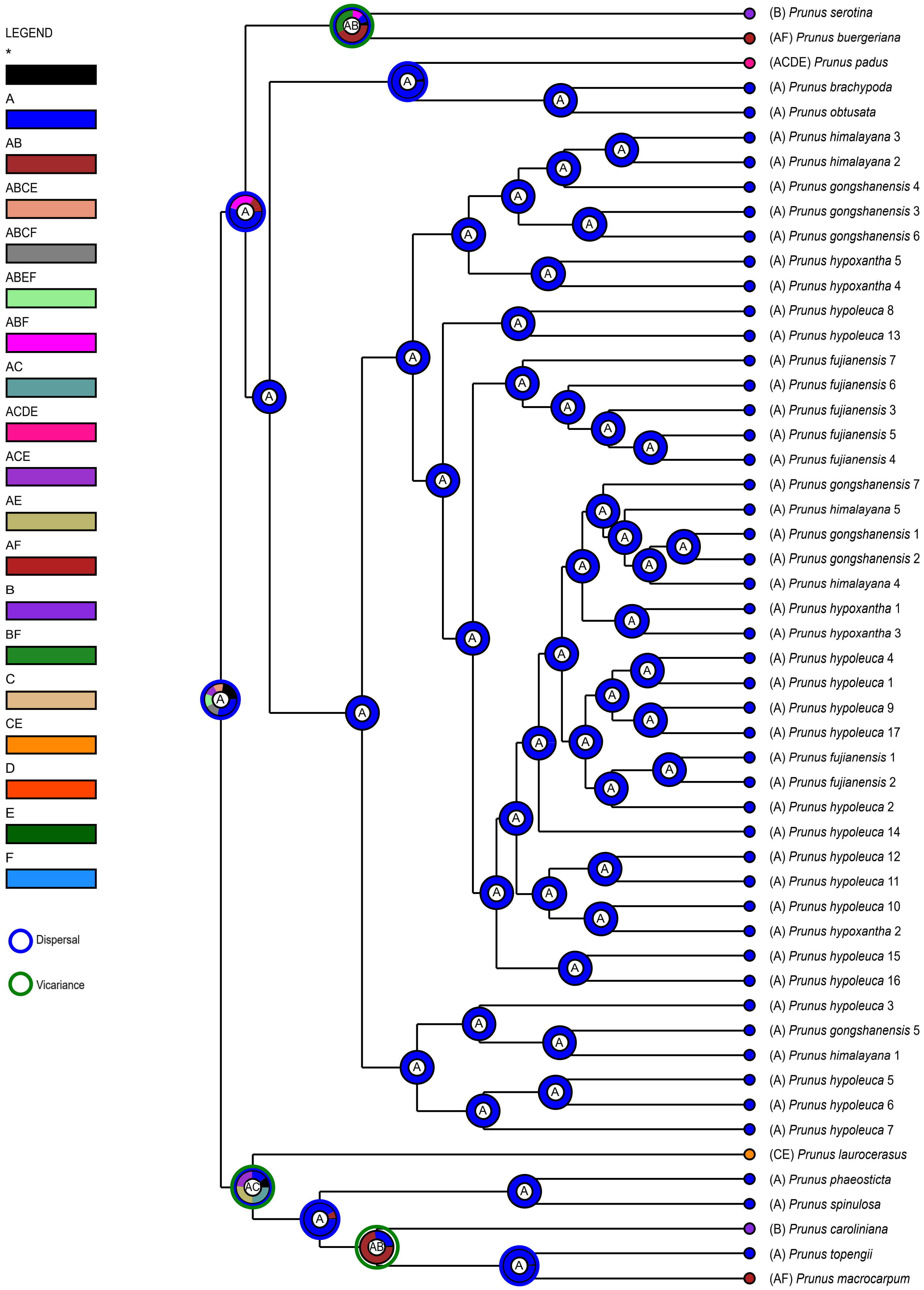

### Supplementary Fig. 43

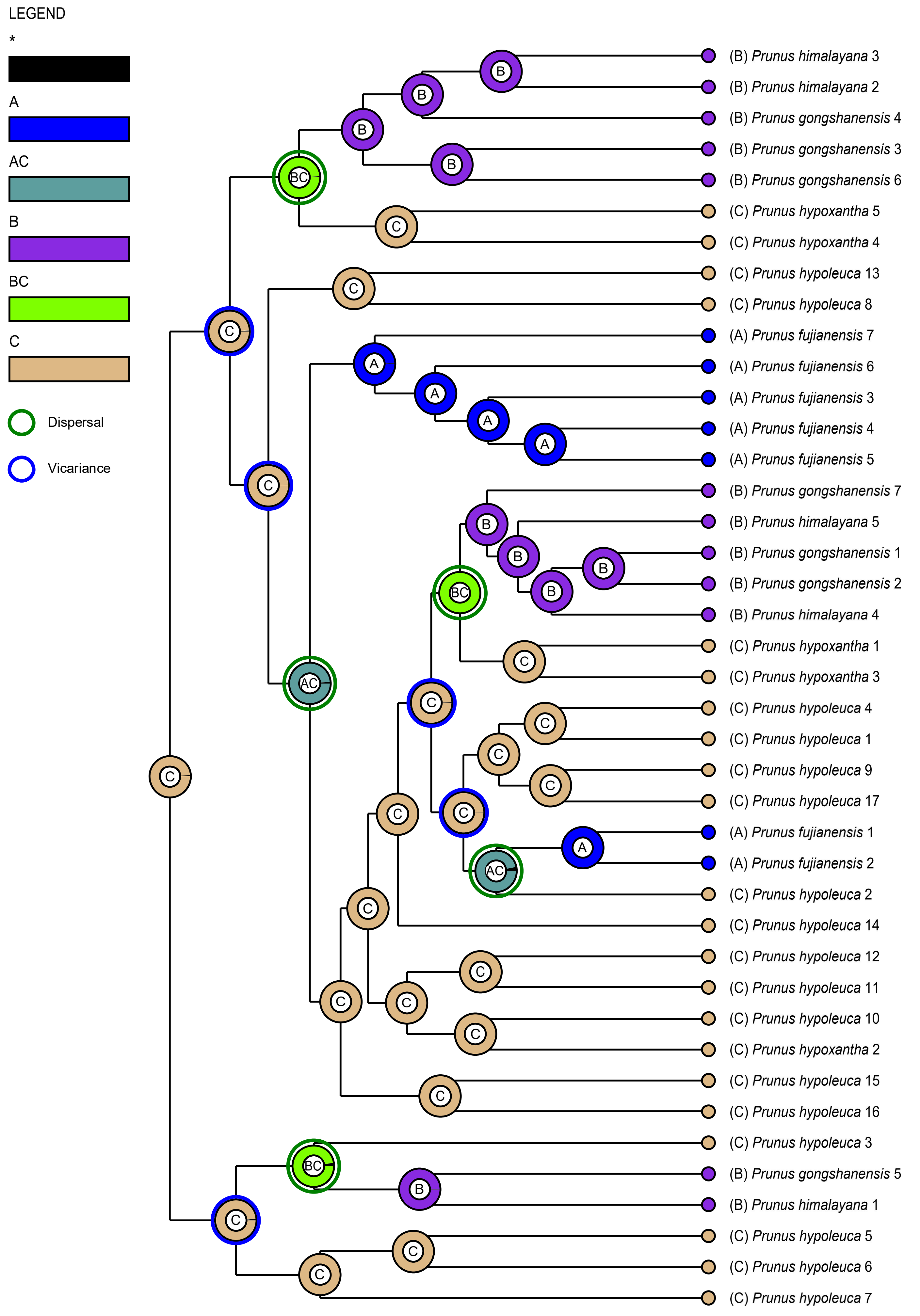

### Supplementary Fig. 44

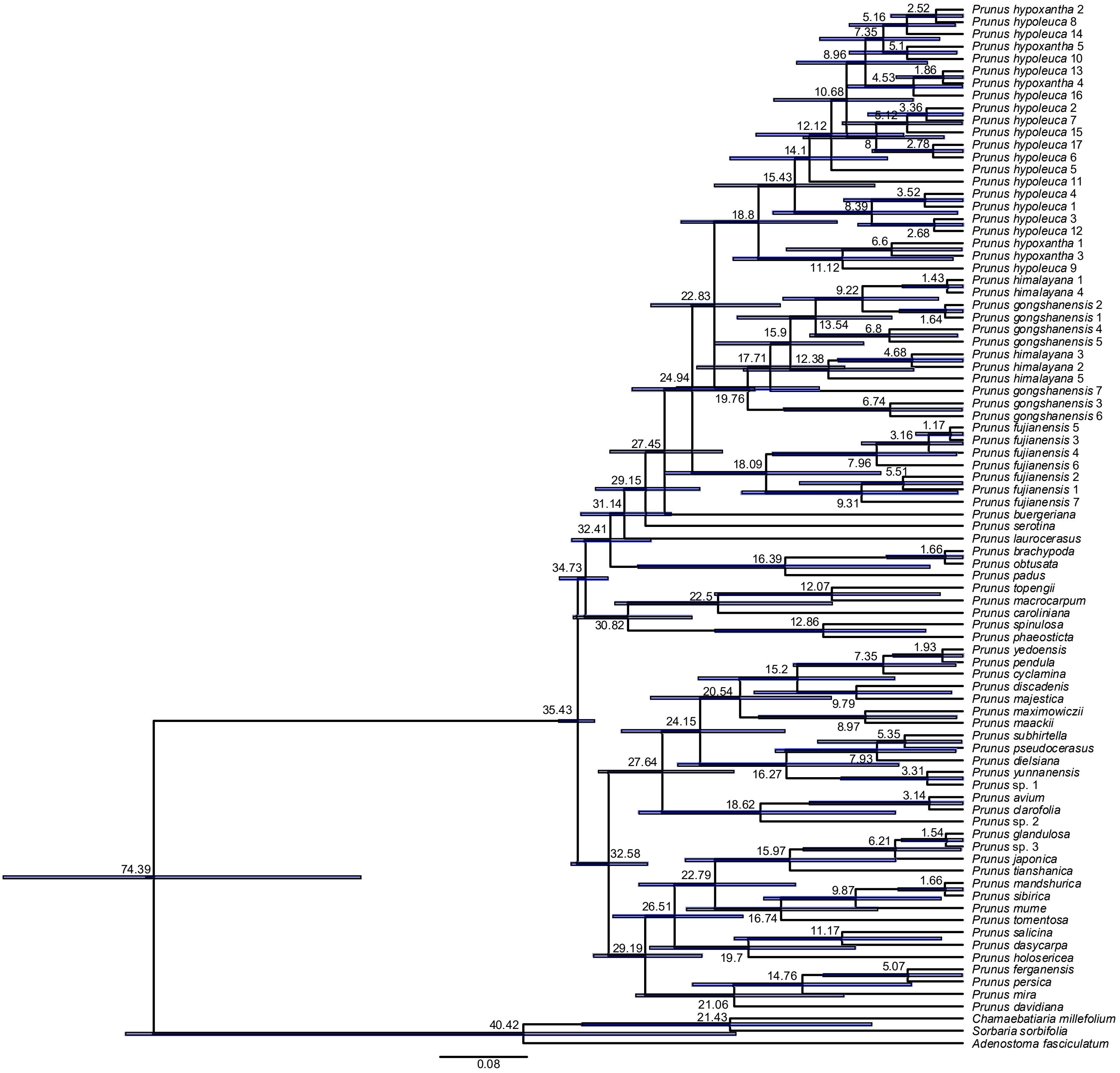

### Supplementary Fig. 45

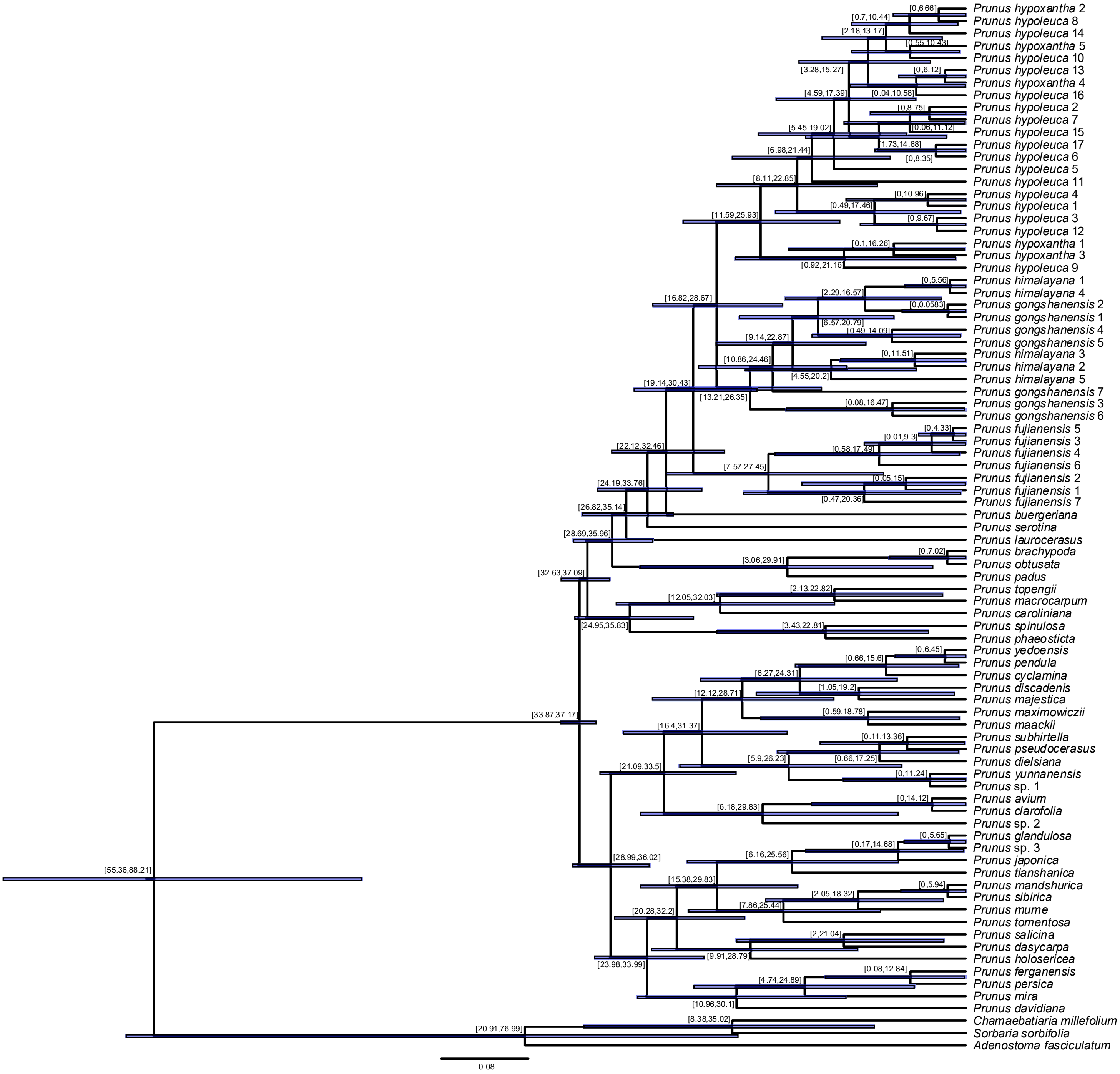

### Supplementary Fig. 46

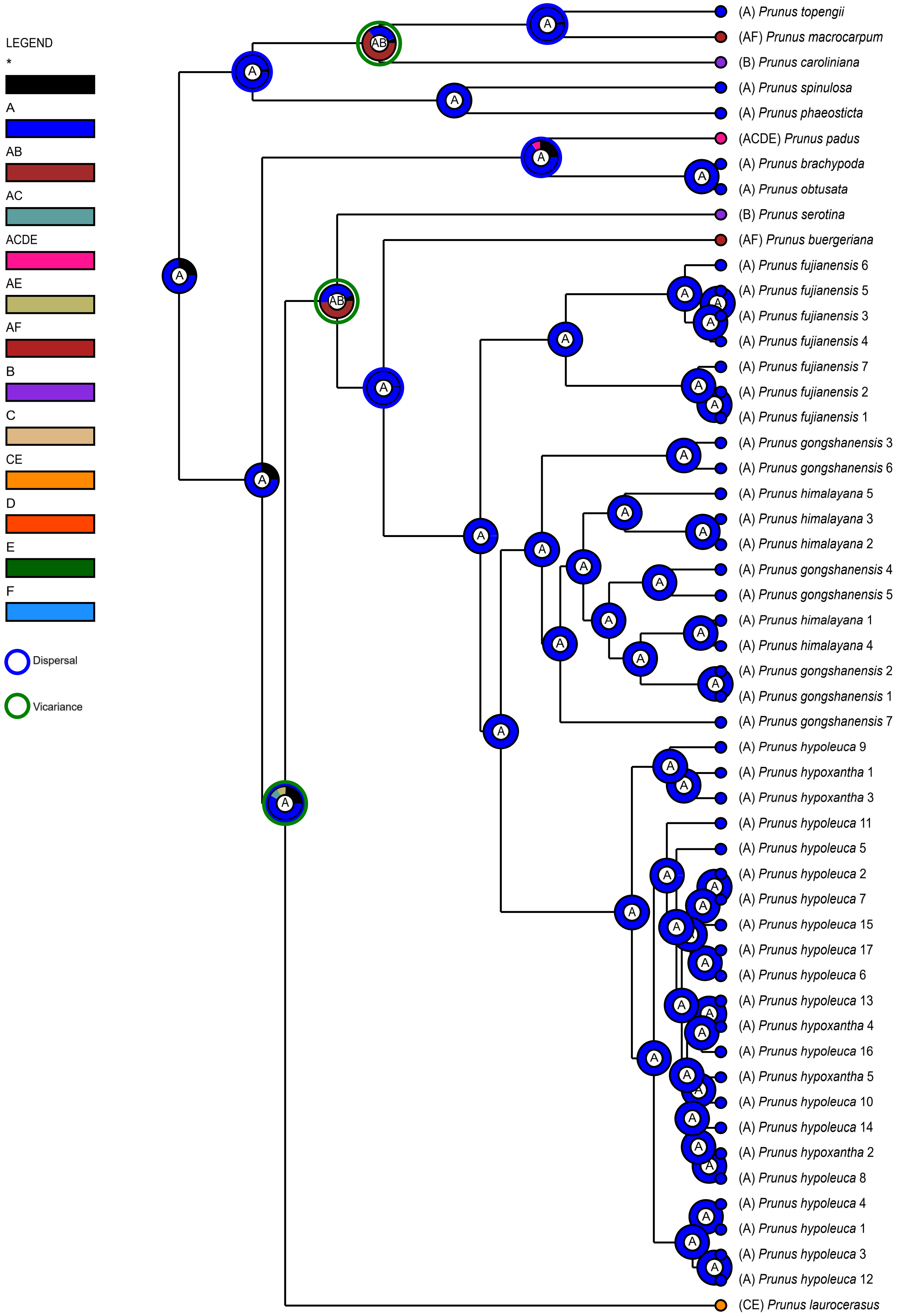

### Supplementary Fig. 47

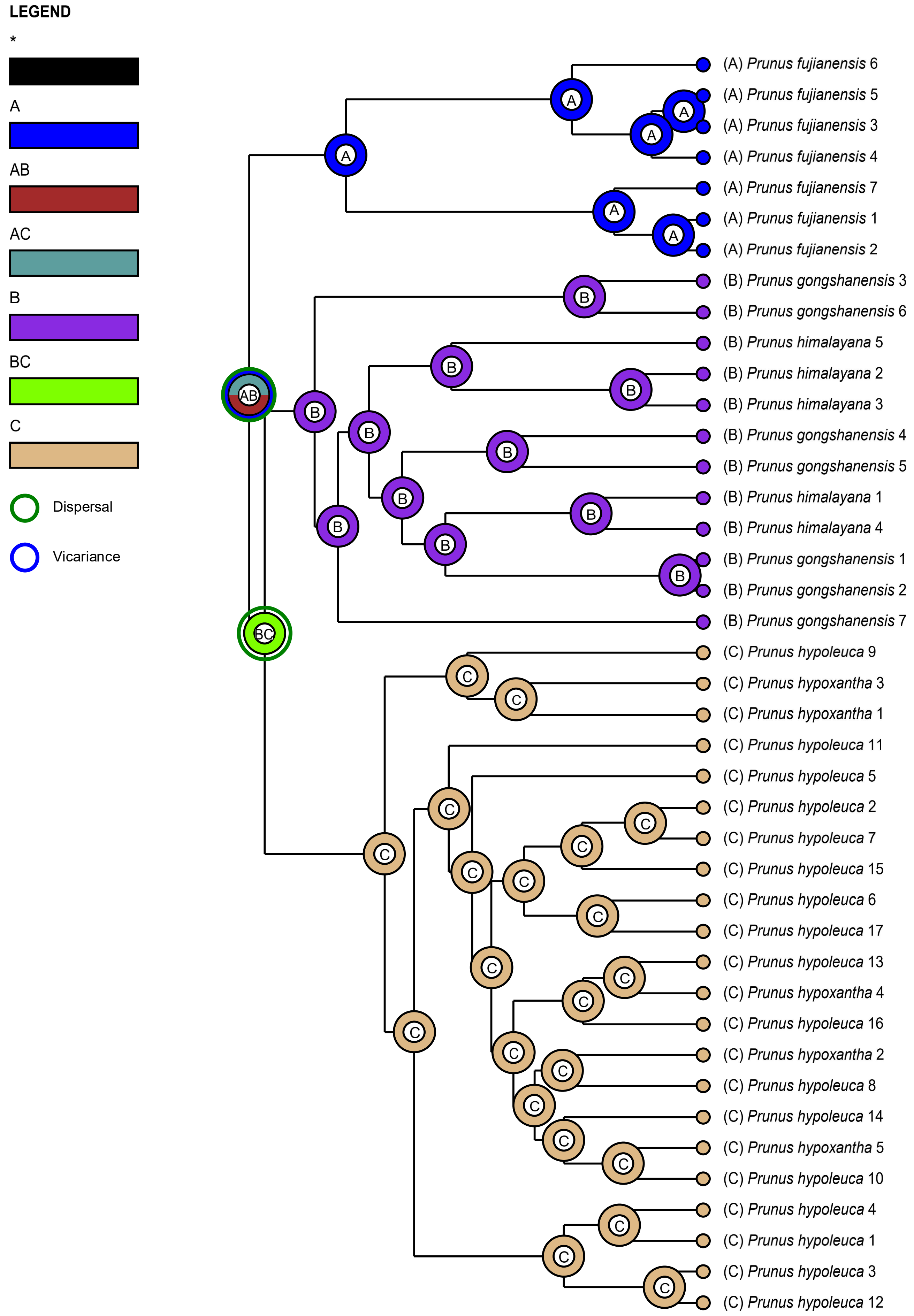

### Supporting Information

**Supplementary Fig. 1** Heat map showing the percentage of sequence length recovery for single-copy nuclear genes (SCN genes) assembled by HybPiper. Each row represents an individual sample, while each column corresponds to a specific gene. The shading intensity within each cell indicates the proportion of the length of each gene recovered for that sample, relative to the average reference length (maximum of 1.0).

**Supplementary Fig. 2** Heat map showing the percentage of sequence length recovery for plastid coding sequences (plastid CDS) assembled by HybPiper. Each row represents an individual sample, while each column corresponds to a specific gene. The shading intensity within each cell indicates the proportion of the length of each gene recovered for that sample, relative to the average reference length (maximum of 1.0).

**Supplementary Fig. 3** Species tree of polyploid *Prunus* in the framework of subfamily Amygdaloideae, inferred by ASTRAL-III based on 1,622 Monophyletic Outgroup (MO) orthologs. Local posterior probabilities (LPP) are shown above the branches.

**Supplementary Fig. 4** Maximum likelihood phylogeny of polyploid *Prunus* in the framework of Amygdaloideae, inferred by RAxML based on 1,622 Monophyletic Outgroup (MO) orthologs. Bootstrap support (BS) is shown above the branches.

**Supplementary Fig. 5** Maximum likelihood phylogeny of polyploid *Prunus* in the framework of Amygdaloideae, inferred by IQ-TREE2 based on 1,622 Monophyletic Outgroup (MO) orthologs. The SH-aLRT support and Ultrafast Bootstrap support (UFBoot) are shown above the branches.

**Supplementary Fig. 6** Species tree of polyploid *Prunus* in the framework of subfamily Amygdaloideae, inferred by ASTRAL-III based on 2,268 RooTed ingroup (RT) orthologs. Local posterior probabilities (LPP) are shown above the branches.

**Supplementary Fig. 7** Maximum likelihood phylogeny of polyploid *Prunus* in the framework of Amygdaloideae, inferred by RAxML based on 2,268 RooTed ingroup (RT) orthologs. Bootstrap support (BS) is shown above the branches.

**Supplementary Fig. 8** Maximum likelihood phylogeny of polyploid *Prunus* in the framework of Amygdaloideae, inferred by IQ-TREE2 based on 2,268 RooTed ingroup (RT) orthologs. The SH-aLRT support and Ultrafast Bootstrap support (UFBoot) are shown above the branches.

**Supplementary Fig. 9** Species tree of polyploid *Prunus* in the framework of subfamily Amygdaloideae, inferred by ASTRAL-III based on plastid coding sequences (plastid CDS) dataset. Local posterior probabilities (LPP) are shown above the branches.

**Supplementary Fig. 10** Maximum likelihood phylogeny of polyploid *Prunus* in the framework of Amygdaloideae, inferred by RAxML based on plastid coding sequences (plastid CDS) dataset. Bootstrap support (BS) is shown above the branches.

**Supplementary Fig. 11** Maximum likelihood phylogeny of polyploid *Prunus* in the framework of Amygdaloideae, inferred by IQ-TREE2 based on plastid coding sequences (plastid CDS) dataset. The SH-aLRT support and Ultrafast Bootstrap support (UFBoot) are shown above the branches.

**Supplementary Fig. 12** Species tree of polyploid *Prunus* in the framework of Amygdaloideae inferred by ASTRAL-III based on 1,622 Monophyletic Outgroup (MO) orthologs. Pie charts on nodes denote the proportion of gene trees that support that clade (blue), the proportion that support the main alternative bifurcation (green), the proportion that support the remaining alternatives (red), the proportion (conflict or support) that have $< 50\%$ bootstrap support (dark grey), and the proportion that have missing taxa (light grey). The number of gene trees concordant with that node in the nuclear phylogeny is shown above branches. The number of gene trees conflicting with that node in the nuclear phylogeny is shown below branches.

**Supplementary Fig. 13** Species tree of polyploid *Prunus* in the framework of Amygdaloideae inferred by ASTRAL-III based on 1,622 Monophyletic Outgroup (MO) orthologs. The Internode Certainty All (ICA) score are shown above branches.

**Supplementary Fig. 14** ASTRAL-III Species tree of polyploid *Prunus* in the framework of Amygdaloideae inferred from 1,622 Monophyletic Outgroup (MO) orthologs. Quartet Sampling (QS) scores for each node are shown next to branches indicating Quartet Concordance (QC) / Quartet Differential (QD) / Quartet Informativeness (QI). Quartet Concordance is also shown at the tip of each sample and is color-coded according to the legend.

**Supplementary Fig. 15** Maximum likelihood phylogeny of polyploid *Prunus* in the framework of Amygdaloideae, inferred by RAxML based on plastid coding sequences (plastid CDS) dataset. Pie charts on nodes denote the proportion of gene trees that support that clade (blue), the proportion that support the main alternative bifurcation (green), the proportion that support the remaining alternatives (red), the proportion (conflict or support) that have < 50% bootstrap support (dark grey), and the proportion that have missing taxa (light grey). The number of gene trees concordant with that node in the nuclear phylogeny are shown above branches. The number of gene trees conflicting with that node in the nuclear phylogeny are shown below branches.

**Supplementary Fig. 16** Maximum likelihood phylogeny of polyploid *Prunus* in the framework of Amygdaloideae inferred from RAxML analysis of plastid coding sequences (plastid CDS). The Internode Certainty All (ICA) score are shown above branches.

**Supplementary Fig. 17** Maximum likelihood phylogeny of polyploid *Prunus* in the framework of

Amygdaloideae inferred from RAxML analysis of plastid coding sequences (plastid CDS). Quartet Sampling (QS) scores for each node are shown next to branches indicating Quartet Concordance (QC) / Quartet Differential (QD) / Quartet Informativeness (QI). Quartet Concordance is also shown at the tip of each sample and is color-coded according to the legend.

**Supplementary Fig. 18** Maximum likelihood phylogeny of polyploid *Prunus* in the framework of Amygdaloideae, inferred by RAxML based on 1,622 Monophyletic Outgroup (MO) orthologs. Pie charts on nodes denote the proportion of gene trees that support that clade (blue), the proportion that support the main alternative bifurcation (green), the proportion that support the remaining alternatives (red), the proportion (conflict or support) that have < 50% bootstrap support (dark grey), and the proportion that have missing taxa (light grey). The number of gene trees concordant with that node in the nuclear phylogeny are shown above branches. The number of gene trees conflicting with that node in the nuclear phylogeny are shown below branches.

**Supplementary Fig. 19** Maximum likelihood phylogeny of polyploid *Prunus* in the framework of Amygdaloideae inferred from RAxML analysis of 1,622 Monophyletic Outgroup (MO) orthologs. The Internode Certainty All (ICA) score are shown above branches.

**Supplementary Fig. 20** Maximum likelihood phylogeny of polyploid *Prunus* in the framework of Amygdaloideae inferred from RAxML analysis of 1,622 Monophyletic Outgroup (MO) orthologs. Quartet Sampling (QS) scores for each node are shown next to branches indicating Quartet Concordance (QC) / Quartet Differential (QD) / Quartet Informativeness (QI). Quartet Concordance is also shown at the tip of each sample and is color-coded according to the legend.

**Supplementary Fig. 21** ASTRAL-III species tree of polyploid *Prunus* in the framework of Amygdaloideae inferred from plastid coding sequences (plastid CDS). Pie charts on nodes denote the proportion of gene trees that support that clade (blue), the proportion that support the main alternative bifurcation (green), the proportion that support the remaining alternatives (red), the proportion (conflict or support) that have < 50% bootstrap support (dark grey), and the proportion that have missing taxa (light grey). The number of gene trees concordant with that node in the nuclear phylogeny are shown above branches. The number of gene trees conflicting with that node in the nuclear phylogeny are shown below branches.

**Supplementary Fig. 22** ASTRAL-III Species tree of polyploid *Prunus* in the framework of Amygdaloideae inferred from plastid coding sequences (plastid CDS). The Internode Certainty All (ICA) scores are shown above branches.

**Supplementary Fig. 23** ASTRAL-III Species tree of polyploid *Prunus* in the framework of Amygdaloideae inferred from plastid coding sequences (plastid CDS). Quartet Sampling (QS) scores for each node are shown next to branches indicating Quartet Concordance (QC) / Quartet Differential (QD) / Quartet Informativeness (QI). Quartet Concordance is also showed in each node'pie chart and color-coded according to the legend.

**Supplementary Fig. 24** Maximum likelihood phylogeny of the *Maddenia* group, with *Prunus serotina* as the outgroup, inferred by RAxML utilizing whole plastome data. Bootstrap support values (BS) are displayed above the branches.

**Supplementary Fig. 25** Maximum likelihood phylogeny of the *Maddenia* group, with *Prunus serotina* as the outgroup, inferred by IQ-TREE2 utilizing whole plastome data. The SH-aLRT support and Ultrafast Bootstrap support (UFBoot) are shown above branches.

**Supplementary Fig. 26** ASTRAL-III Species tree of polyploid *Prunus* in the framework of Amygdaloideae inferred from 1,622 Monophyletic Outgroup (MO) orthologs. Population mutation parameter theta values are shown above branches.

**Supplementary Fig. 27** The determination of DNA ploidy in certain samples of the *Maddenia* group using flow cytometry. PI-A: PI-area.

**Supplementary Fig. 28** Heat map showing number of paralog sequences for each gene and each sample recovered by HybPiper. Each row shows a sample, and each column is a gene. The amount of shading in each box corresponds to the number of the gene recovered for that sample by the pipeline.

**Supplementary Fig. 29** WGDs identified in polyploid *Prunus* by tree reconciliation method with Tree2GD. Values of each node shows the total number of gene families (blue-purple box), the number of newly gained gene families (green), the number of lost gene families (red), the number of gene duplication (GD) / the total number of gene family trees of the node (orange box) and the promotion of gene duplication (green box).

**Supplementary Fig. 30** Optimal multi-labeled trees (MUL-trees) inferred from GRAMPA analyses by mapping homolog trees onto a species tree derived using Monophyletic Outgroup (MO) ortholog trees, with polyploid *Prunus* identified as having an allopolyploid origin. In this clade, multiple labels indicate the origins of polyploidy; a plus sign marks the first tip, while an asterisk denotes the second tip.

**Supplementary Fig. 31** Heat map showing statical support for gene flow among species pairs in *Prunus* s.l. inferred from Dsuite package. The shaded scale in boxes represents the estimated $f_4$-ratio branch value.

**Supplementary Fig. 32** Heat map showing statical support for gene flow among species pairs of the *Maddenia* group inferred from Dsuite package. The shaded scale in boxes represents the estimated $f_4$-ratio branch value.

**Supplementary Fig. 33** The matrix shows inferred introgression proportions as estimated from ASTRAL-III species trees in the introgressed species pairs, and then mapped to internal branches using the *f*-branch method. The *f*-branch statistic identifies possible gene flow from the branch of the tree on the *y* axis to the species or population on the *x* axis.

**Supplementary Fig. 34** The most parsimonious multi-labeled trees (MUL-trees) inferred from GRAMPA analyses on the trees inferred from nuclear phylogeny within the *Maddenia* group. The clade with multiple labels denotes the polyploidy origin, a plus sign indicates the first tip, and the second tip is shown with an asterisk.

**Supplementary Fig. 35** The most parsimonious multi-labeled trees (MUL-trees) inferred from GRAMPA analyses on the trees inferred from nuclear phylogeny within the *Maddenia* clade after removing the clade identified as allopolyploidy (Supplementary Fig. 34). The clade with multiple labels denotes the polyploidy origin, a plus sign indicates the first tip, and the second tip is shown with an asterisk.

**Supplementary Fig. 36** Dated chronogram of Amygdaloideae inferred from PAML based on the RAxML concatenated tree inferred from Monophyletic Outgroup (MO) orthologs. Maximum clade credibility (MCC) tree showing mean ages above branches. Light blue bars on nodes represent 95% confidence intervals of divergence time estimates.

**Supplementary Fig. 37** Dated chronogram of Amygdaloideae inferred from PAML based on the RAxML concatenated tree inferred from Monophyletic Outgroup (MO) orthologs. Node numbers and blue bars indicate 95% confidence intervals of divergence time estimates.

**Supplementary Fig. 38** Dated chronogram of Amygdaloideae inferred from PAML based on the RAxML concatenated tree inferred from plastid coding sequences (plastid CDS) dataset. Maximum clade credibility (MCC) tree showing mean ages above branches. Light blue bars on nodes represent 95% confidence intervals of divergence time estimates.

**Supplementary Fig. 39** Dated chronogram of Amygdaloideae inferred from PAML based on the RAxML concatenated tree inferred from plastid coding sequences (plastid CDS) dataset. Node numbers and blue bars indicate 95% confidence intervals of divergence time estimates.

**Supplementary Fig. 40** Dated chronogram of *Prunus* s.l. inferred from PAML based on the RAxML concatenated tree inferred from Monophyletic Outgroup (MO) orthologs. Maximum clade credibility (MCC) tree showing mean ages above branches. Light blue bars on nodes represent 95% confidence intervals of divergence time estimates.

**Supplementary Fig. 41** Dated chronogram of *Prunus* s.l. inferred from PAML based on the RAxML concatenated tree inferred from Monophyletic Outgroup (MO) orthologs. Node numbers and blue bars indicate 95% confidence intervals of divergence time estimates.

**Supplementary Fig. 42** The ancestral area reconstruction using BioGeoBEARS implemented in RASP using the dated chronogram of polyploid *Prunus* inferred from PAML based on the RAxML concatenated tree inferred from Monophyletic Outgroup (MO) orthologs, with the colored key identifying extant and possible ancestral ranges. (A) East Asia, (B) Americas, (C) West Asia, (D) Europe, (E) Africa, and (F) Australasia.

**Supplementary Fig. 43** The ancestral area reconstruction using BioGeoBEARS implemented in RASP using the dated chronogram of the *Maddenia* group inferred from PAML based on the RAxML concatenated tree inferred from Monophyletic Outgroup (MO) orthologs, with the colored key identifying extant and possible ancestral ranges. (A) Southeast China (Fujian, Zhejiang, Anhui, and Jiangxi); (B) Southwest China (Xizang and Yunnan); (C) Central China (Gansu, Qinghai,

Shaanxi, Hubei, Hunan, Henan, Chongqing, and Sichuan).

**Supplementary Fig. 44** Dated chronogram of *Prunus* s.l. inferred from PAML based on the RAxML concatenated tree inferred from plastid coding sequences (plastid CDS) dataset. Maximum clade credibility (MCC) tree showing mean ages above branches. Light blue bars on nodes represent 95% confidence intervals of divergence time estimates.

**Supplementary Fig. 45** Dated chronogram of *Prunus* s.l. inferred from PAML based on the RAxML concatenated tree inferred from plastid coding sequences (plastid CDS) dataset. Node numbers and blue bars indicate 95% confidence intervals of divergence time estimates.

**Supplementary Fig. 46** The ancestral area reconstruction using BioGeoBEARS implemented in RASP using the dated chronogram of polyploid *Prunus* inferred from PAML based on the RAxML concatenated tree inferred from plastid coding sequences (plastid CDS) dataset, with the colored key identifying extant and possible ancestral ranges. (A) East Asia, (B) Americas, (C) West Asia, (D) Europe, (E) Africa, and (F) Australasia.

**Supplementary Fig. 47** The ancestral area reconstruction using BioGeoBEARS implemented in RASP using the dated chronogram of the *Maddenia* group inferred from PAML based on the RAxML concatenated tree inferred from plastid coding sequences (plastid CDS) dataset, with the colored key identifying extant and possible ancestral ranges. (A) Southeast China (Fujian, Zhejiang, Anhui, and Jiangxi); (B) Southwest China (Xizang and Yunnan); (C) Central China (Gansu, Qinghai, Shaanxi, Hubei, Hunan, Henan, Chongqing, and Sichuan).

**Supplementary Table 1** Voucher and sequence information for taxa of Amygdaloideae used in this study. † represents the data newly sequenced for this article.

**Supplementary Table 2** HybPiper assembly and orthology inference statistics.

**Supplementary Table 3** Fossil records used for calibrition in this study.

### Supplementary Methods

#### Orthology inference and dataset generation

The identification of potential paralogs is critical for ensuring the accuracy of phylogenetic inferences. In this study, we implemented an automated orthology inference approach for nuclear genes assembled from DGS reads. This approach, initially conceptualized for RNA-Seq data analysis by Yang & Smith[1], underwent subsequent modifications for adapting the target enrichment datasets, such as Hyb-Seq[2]. Our PhyloAI team has executed further refinements to these scripts, ensuring their compatibility with DGS datasets. This enhancement has been substantiated through its successful application in recent studies, such as the taxonomic research on *Pyrus*[3].

The procedure involves utilizing the script "mafft_wrapper.py" for sequence alignment by MAFFT v. 7.520[4], followed by the trimming of columns containing more than 90% missing data using "pxclsq_wrapper.py" with *phyx*[5]. Maximum Likelihood (ML) phylogenetic trees of homologs were then inferred using "raxml_bs_wrapper.py" with RAxML v. 8.2.13[6]. Then, the script "mask_tips_by_taxonID_transcripts.py" was employed to mask both mono- and paraphyletic tips belonging to the same taxon. Additionally, TreeShrink v. 1.3.9[7] was implemented with the script "tree_shrink_wrapper.py" to detect and remove outlier branches exhibiting excessive length in the ML trees. Before inferring homolog phylogenies, two distinct orthology inference strategies were applied. In this analysis, *Rosa rugosa* was designated as the outgroup, with all other samples classified as ingroups. Initially, the "Monophyletic Outgroup" (MO) approach was employed to identify homolog trees where the outgroup taxa are monophyletic and present in a single copy. This was followed by the "RooTed ingroup" (RT) strategy, facilitating the extraction of ingroup clades by pruning paralogs from root to tip, or retaining taxa without duplication in the absence of an outgroup.

The subsequent analyses were performed on these two datasets, i.e., MO and RT. For each SCN locus, alignment was conducted utilizing MAFFT v. 7.520[4] with parameters "--maxiterate 1,000 --localpair". We performed the following strategies in the SCN loci-cleaning process to account for low-quality regions within the alignment sequences. The application of trimAl v. 1.4.1[8], with parameters "-gt 0.8 -st 0.001", further facilitated the exclusion of columns exhibiting more than 20%

missing data or similarity scores beneath 0.001. Following sequence concatenation via AMAS v. 1.0[9], spruceup v. 2022.2.4[10] was utilized to identify and excise poorly aligned sequence blocks, with window size of 50 and overlap of 25. Subsequently, we used AMAS v. 1.0[9] to separate the trimmed alignment into individual locus alignments, which were again refined using trimAl v. 1.4.1[8] with the previously specified parameters. Considering the limited informative sites in short sequences, we excluded sequences of length less than 150 bp via the "exclude_short_sequences.py" script, as delineated by Liu et al.[11]. These refined SCN loci were used to infer ML trees utilizing RAxML v. 8.2.13[6], under the GTRGAMMA model with parameters "-f a -p 12345 -x 12345" and 200 Bootstrap replicates. TreeShrink v. 1.3.9[7] was used to identify and remove anomalously long branches within the ML trees and their associate alignments, utilizing the default false positive tolerance rate and per-species mode. Following this comprehensive loci-cleaning procedure, we produced two datasets—MO and RT—comprising clean sequences and refined ML trees, serving as the foundation for subsequent phylogenetic analysis.

**Calibrations information of historical biogeographic analysis**

We integrated eight fossil calibrations and one secondary calibration node (detailed in Supplementary Table 3), and these nine calibrations were scattered across the Amygdaloideae phylogeny to refine divergence time estimations. The amber fossil of *Prunus hirsutipetala* (F1), discovered in NW Ukraine and dating back to the Priabonian of Eocene (37.2–33.9 Million years ago (Mya))[12], served to calibrate the stem clade of polyploid *Prunus*. Additionally, *P. wutuensis* (F2), discovered in Wutu Coal Mine, Shandong, China, and dating to the Early Eocene (approximately 55 Mya)[13], was assigned to the stem of *Prunus*. Fossils of *Neviusia* (F3) and *Spiraea* (F7), dated to the Early Eocene (50–49 Mya) and located in Republic, Washington, USA[14,15], were utilized to constrain the stem ages of *Neviusia* and *Spiraea*, respectively. The flower fossil of *Oemleria janhartfordae* (F4) from the Klondike Mountain Formation in Washington, USA, dating to the Early Eocene (approximately 49.96–48.88 Mya)[16], informed the calibration of stem *Oemleria*. The fossils of *Amelanchier peritula* and *A. scudderi* (F5) from the Florissant Formation in Colorado, USA, dating to the Late Eocene (37.2–33.9 Mya)[17,18], calibrated stem *Amelanchier*. The fossil of *Vauquelinia* *comptonifolia* (F6), found in Wyoming, USA, and dated to the Middle Eocene (46.2–40.4 Mya)[19], was allocated to stem *Vauquelinia*. Furthermore, the *Holodiscus lisii* fossil (F8), from Florissant, Colorado, USA, and dating to the Late Eocene (approximately 34 Mya)[20,21], calibrated stem *Holodiscus*. Lastly, the secondary calibration node (C1) for the stem Amygdaloideae was estimated at 96.36-94.46 Mya based on plastid data[22].

**Reference**


1. Yang, Y. & Smith, S. A. Orthology inference in nonmodel organisms using transcriptomes and low-coverage genomes: improving accuracy and matrix occupancy for phylogenomics. *Mol. Biol. Evol.* **31**, 3081–3092 (2014).
2. Morales-Briones, D. F. *et al.* Analysis of paralogs in target enrichment data pinpoints multiple ancient polyploidy events in *Alchemilla* s.l. (Rosaceae). *Syst. Biol.* **71**, 190–207 (2022).
3. Jin, Z. T. *et al.* Advancing *Pyrus* phylogeny: deep genome skimming-based inference coupled with paralogy analysis yields a robust phylogenetic backbone and an updated infrageneric classification of the pear genus (Maleae, Rosaceae). *TAXON* **73**, 784–799 (2024).
4. Nakamura, T., Yamada, K. D., Tomii, K. & Katoh, K. Parallelization of MAFFT for large-scale multiple sequence alignments. *Bioinformatics* **34**, 2490–2492 (2018).
5. Brown, J. W., Walker, J. F. & Smith, S. A. Phyx: phylogenetic tools for unix. *Bioinformatics* **33**, 1886–1888 (2017).
6. Stamatakis, A. RAxML version 8: a tool for phylogenetic analysis and post-analysis of large phylogenies. *Bioinformatics* **30**, 1312–1313 (2014).
7. Mai, U. & Mirarab, S. TreeShrink: fast and accurate detection of outlier long branches in collections of phylogenetic trees. *BMC Genom.* **19**, 272 (2018).
8. Capella-Gutiérrez, S., Silla-Martínez, J. M. & Gabaldón, T. trimAl: a tool for automated alignment trimming in large-scale phylogenetic analyses. *Bioinformatics* **25**, 1972–1973 (2009).
9. Borowiec, M. L. AMAS: a fast tool for alignment manipulation and computing of summary statistics. *PeerJ* **4**, e1660 (2016).
10. Borowiec, M. Spruceup: fast and flexible identification, visualization, and removal of outliers

from large multiple sequence alignments. *J. Open Source Softw.* **4**, 1635 (2019).
11. Liu, B. B. *et al.* Phylogenomic conflict analyses in the apple genus *Malus* s.l. reveal widespread hybridization and allopolyploidy driving diversification, with insights into the complex biogeographic history in the Northern Hemisphere. *J. Integr. Plant Biol.* **64**, 1020–1043 (2022).
12. Sokoloff, D. D. *et al.* Staminate flower of *Prunus* s.l. (Rosaceae) from Eocene Rovno amber (Ukraine). *J. Plant Res.* **131**, 925–943 (2018).
13. Li, Y. *et al.* Endocarps of *Prunus* (Rosaceae: Prunoideae) from the early Eocene of Wutu, Shandong Province, China. *Taxon* **60**, 555–564 (2011).
14. Mathews, W. H. Potassium-argon age determinations of Cenozoic volcanic rocks from British Columbia. *Geol. Soc. Am. Bull.* **75**, 465–468 (1964).
15. Wehr, W. C. & Hopkins, D. Q. The Eocene orchards and gardens of Republic, Washington. *Washington Geology* **22**, 27–34 (1994).
16. Benedict, J. C., DeVore, M. L. & Pigg, K. B. *Prunus* and *Oemleria* (Rosaceae) flowers from the late early Eocene republic flora of Northeastern Washington State, U.S.A. *Int. J. Plant Sci.* **172**, 948–958 (2011).
17. Cockerell, T. D. A. Fossil insects from Florissant, Colorado. *Bull. Amer. Mus. Nat. Hist.* **30**, 71–82 (1911).
18. MacGinitie, H. D. *Fossil Plants of the Florissant Beds, Colorado*. (Carnegie Institution of Washington Publication 599, Washington, D. C., 1953).
19. Macginitie, H. D. *The Eocene Green River Flora of Northwestern Colorado and Northeastern Utah*. (University of California Press, Berkeley, 1969).
20. Schorn, H. E. *Holodiscus lisii* (Rosaceae): a new species of ocean spray from the late Eocene Florissant Formation, Colorado, USA. *PaleoBios* **18**, 21–24 (1998).
21. Mclntosh, W. C. & Chapin, C. E. Geochronology of the Central Colorado volcanic field. *New Mexico Bureau of Geology and Mineral Resources Bulletin* **160**, 205–237 (2004).
22. Zhang, S. D. *et al.* Diversification of Rosaceae since the Late Cretaceous based on plastid phylogenomics. *New Phytol.* **214**, 1355–1367 (2017).
