## Supplemental Table 1 for "Unveiling allopolyploidization-driven genome duplications through progressive analysis of deep genome skimming data"

| Number | Group type | Genera | Subgenera | Species | Sample name | Sample number | NCBI BioProject No. | SRA accession | Origin | voucher | accession_cp | No. of clean reads | No. of bases (bp) | Data type | Data size (G) | chromosome ploidy |
|---|---|---|---|---|---|---|---|---|---|---|---|---|---|---|---|---|
| 1 | ingroup | *Prunus* s.l. | subg. *Padus* | *Prunus fujianensis* (Y.T.Chang) J.Wen | *Prunus fujianensis* 1 | xsy836 | PRJNA1031385† | SRR28365552 | Wuyishan, Fujian, China | L.Zhao#xsy836 (WUK) | PP546823 | 166886168 | 25032925200 | DGS | 23.31 | 2n=4x=32 |
| 2 | ingroup | *Prunus* s.l. | subg. *Padus* | *Prunus fujianensis* | *Prunus fujianensis* 2 | xsy844 | PRJNA1031385† | SRR28365551 | Wuyishan, Fujian, China | L.Zhao#xsy844 (WUK) | PP595848 | 163167226 | 24475083900 | DGS | 22.79 | |
| 3 | ingroup | *Prunus* s.l. | subg. *Padus* | *Prunus fujianensis* | *Prunus fujianensis* 3 | SN511 | PRJNA1031385† | SRR28365544 | Huangshan, Anhui, China | L.Zhao#SN511 (WUK) | PP595862 | 83808988 | 12571348200 | DGS | 11.71 | |
| 4 | ingroup | *Prunus* s.l. | subg. *Padus* | *Prunus fujianensis* | *Prunus fujianensis* 4 | SN512 | PRJNA1031385† | SRR28365543 | Huangshan, Anhui, China | L.Zhao#SN512 (WUK) | PP595863 | 121863964 | 18279594600 | DGS | 17.02 | |
| 5 | ingroup | *Prunus* s.l. | subg. *Padus* | *Prunus fujianensis* | *Prunus fujianensis* 5 | SN513 | PRJNA1031385† | SRR28365542 | Huangshan, Anhui, China | L.Zhao#SN513 (WUK) | PP595864 | 89685070 | 13452760500 | DGS | 12.53 | |
| 6 | ingroup | *Prunus* s.l. | subg. *Padus* | *Prunus fujianensis* | *Prunus fujianensis* 6 | SN514 | PRJNA1031385† | SRR28365541 | Anji, Huzhou, Zhejiang, China | L.P.Yu & M.B.Deng#97099(B) (PE 00995643) | PP595865 | 97483868 | 14622580200 | DGS | 13.62 | |
| 7 | ingroup | *Prunus* s.l. | subg. *Padus* | *Prunus fujianensis* | *Prunus fujianensis* 7 | PE909 | PRJNA1031385† | SRR28365527 | Jinzhai, Lu'an, Anhui, China | s.coll.#0181 (PE 00995640) | PP648246 | 132721316 | 19908197400 | DGS | 18.54 | |
| 8 | ingroup | *Prunus* s.l. | subg. *Padus* | *Prunus gongshanensis* J.Wen | *Prunus gongshanensis* 1 | GBJ08133 | PRJNA1031385† | SRR28456453 | Chayu, Nyingchi, Xizang, China | s.coll.#49941 (CSHM) | PP595849 | 70939130 | 10640869500 | DGS | 9.91 | |
| 9 | ingroup | *Prunus* s.l. | subg. *Padus* | *Prunus gongshanensis* | *Prunus gongshanensis* 2 | GBJ09238 | PRJNA1031385† | SRR28456452 | Chayu, Nyingchi, Xizang, China | s.coll.#50130 (CSHM) | PP595850 | 67170668 | 10075600200 | DGS | 9.38 | |
| 10 | ingroup | *Prunus* s.l. | subg. *Padus* | *Prunus gongshanensis* | *Prunus gongshanensis* 3 | JR305 | PRJNA1031385† | SRR28365540 | Tengchong, Baoshan, Yunnan, China | s.coll.#GLGS30758 (WUK) | PP595851 | 149826116 | 22473917400 | DGS | 20.93 | |
| 11 | ingroup | *Prunus* s.l. | subg. *Padus* | *Prunus gongshanensis* | *Prunus gongshanensis* 4 | JR308 | PRJNA1031385† | SRR28365529 | Mêdog, Nyingchi, Xizang, China | L.Zhao#JR308 (WUK) | PP595852 | 182209552 | 27331432800 | DGS | 25.45 | |
| 12 | ingroup | *Prunus* s.l. | subg. *Padus* | *Prunus gongshanensis* | *Prunus gongshanensis* 5 | JR379 | PRJNA1031385† | SRR28365518 | Mêdog, Nyingchi, Xizang, China | L.Zhao#JR379 (WUK) | PP595853 | 158371348 | 23755702200 | DGS | 22.12 | |
| 13 | ingroup | *Prunus* s.l. | subg. *Padus* | *Prunus gongshanensis* | *Prunus gongshanensis* 6 | JR381 | PRJNA1031385† | SRR28365507 | Fugong, Nujiang, Yunnan, China | s.coll.#GLGS20474 (WUK) | PP595854 | 174192216 | 26128832400 | DGS | 24.33 | |
| 14 | ingroup | *Prunus* s.l. | subg. *Padus* | *Prunus gongshanensis* | *Prunus gongshanensis* 7 | xsy817 | PRJNA1031385† | SRR28365496 | Gongshan, Nujiang, Yunnan, China | L.Zhao#xsy817 (WUK) | PP595855 | 481515052 | 72227257800 | DGS | 67.27 | 2n=8x=64 |
| 15 | ingroup | *Prunus* s.l. | subg. *Padus* | *Prunus himalayana* Kitam. | *Prunus himalayana* 1 | JR304 | PRJNA1031385† | SRR28365485 | Bomê, Nyingchi, Xizang, China | Qinghai-Xizang Collection Team#94 (PE 00772895) | PP595856 | 128548808 | 19282321200 | DGS | 17.96 | |
| 16 | ingroup | *Prunus* s.l. | subg. *Padus* | *Prunus himalayana* | *Prunus himalayana* 2 | JR377 | PRJNA1031385† | SRR28365479 | Yadong, Shigatse, Xizang, China | s.coll.#750177 (WUK) | PP595857 | 245486784 | 36823017600 | DGS | 34.29 | |
| 17 | ingroup | *Prunus* s.l. | subg. *Padus* | *Prunus himalayana* | *Prunus himalayana* 3 | JR378 | PRJNA1031385† | SRR28365478 | Yadong, Shigatse, Xizang, China | s.coll.#750141 (WUK) | PP595858 | 261661096 | 39249164400 | DGS | 36.55 | |
| 18 | ingroup | *Prunus* s.l. | subg. *Padus* | *Prunus himalayana* | *Prunus himalayana* 4 | xsy848 | PRJNA1031385† | SRR28365550 | Bomê, Nyingchi, Xizang, China | S.L.Zhang3424#YT20144 (CSH) | PP595859 | 127215638 | 19082345700 | DGS | 17.77 | 2n=4x=32 |
| 19 | ingroup | *Prunus* s.l. | subg. *Padus* | *Prunus himalayana* | *Prunus himalayana* 5 | xsy849 | PRJNA1031385† | SRR28365549 | Cuona, Shannan, Xizang, China | PE-Xizang Expedition#7142 (PE 02332297) | PP595860 | 178337194 | 26750579100 | DGS | 24.91 | |
| 20 | ingroup | *Prunus* s.l. | subg. *Padus* | *Prunus hypoleuca* (Koehne) J.Wen | *Prunus hypoleuca* 1 | WX219 | PRJNA1031385† | SRR28365538 | Taibaishan, Baoji, Shaanxi, China | L.Zhao#WX219 (WUK) | PP595861 | 175659460 | 26348919000 | DGS | 24.54 | |
| 21 | ingroup | *Prunus* s.l. | subg. *Padus* | *Prunus hypoleuca* | *Prunus hypoleuca* 2 | JR324 | PRJNA1031385† | SRR28365548 | Shennongjia, Hubei, China | L.Zhao#JR324 (WUK) | PP546822 | 176922850 | 26538427500 | DGS | 24.72 | 2n=4x=32 |
| 22 | ingroup | *Prunus* s.l. | subg. *Padus* | *Prunus hypoleuca* | *Prunus hypoleuca* 3 | SN515 | PRJNA1031385† | SRR28365539 | Xunhua, Haidong, Qinghai, China | s.coll.#0457608 (WUK) | PP648234 | 163680606 | 24552090900 | DGS | 22.87 | |
| 23 | ingroup | *Prunus* s.l. | subg. *Padus* | *Prunus hypoleuca* | *Prunus hypoleuca* 4 | xsy804 | PRJNA1031385† | SRR28365537 | Taibaishan, Baoji, Shaanxi, China | L.Zhao#xsy804 (WUK) | PP648235 | 438944908 | 65841736200 | DGS | 61.32 | |
| 24 | ingroup | *Prunus* s.l. | subg. *Padus* | *Prunus hypoleuca* | *Prunus hypoleuca* 5 | PE905 | PRJNA1031385† | SRR28365547 | Sangzhi, Zhangjiajie, Hunan, China | Beijing Collection Team#002632 (PE 01365138) | PP648236 | 125411946 | 18811791900 | DGS | 17.52 | |
| 25 | ingroup | *Prunus* s.l. | subg. *Padus* | *Prunus hypoleuca* | *Prunus hypoleuca* 6 | PE906 | PRJNA1031385† | SRR28365546 | Chengkou, Chongqing, China | Q.Z.Yang#61 (PE 00772924) | PP648237 | 79566196 | 11934929400 | DGS | 11.12 | |
| 26 | ingroup | *Prunus* s.l. | subg. *Padus* | *Prunus hypoleuca* | *Prunus hypoleuca* 7 | PE915 | PRJNA1031385† | SRR28365545 | Shennongjia Hubei, China | Hubei Shennongjia Collection Team#10443 (PE 00997389) | PP648238 | 79805998 | 11970899700 | DGS | 11.15 | |
| 27 | ingroup | *Prunus* s.l. | subg. *Padus* | *Prunus hypoleuca* | *Prunus hypoleuca* 8 | PE907 | PRJNA1031385† | SRR28365534 | Jiangkou, Tongren, Guizhou, China | Z.S.Zhang et al.#402013 (PE 01296248) | PP648240 | 116234522 | 17435178300 | DGS | 16.24 | |
| 28 | ingroup | *Prunus* s.l. | subg. *Padus* | *Prunus hypoleuca* | *Prunus hypoleuca* 9 | JR301 | PRJNA1031385† | SRR28365533 | Min, Dingxi, Gansu, China | L.Zhao#JR301 (WUK) | PP648241 | 161338728 | 24200809200 | DGS | 22.54 | |
| 29 | ingroup | *Prunus* s.l. | subg. *Padus* | *Prunus hypoleuca* | *Prunus hypoleuca* 10 | PE901 | PRJNA1031385† | SRR28365532 | Wen, Longnan, Gansu, China | Baishuijiang Collection Team#20070422 (PE 01651988) | PP648242 | 119578966 | 17936844900 | DGS | 16.70 | |
| 30 | ingroup | *Prunus* s.l. | subg. *Padus* | *Prunus hypoleuca* | *Prunus hypoleuca* 11 | PE902 | PRJNA1031385† | SRR28365531 | Lianhuashan, Gansu, China | X.G.Sun et al.#2757 (PE 01841627) | PP648243 | 129489286 | 19423392900 | DGS | 18.09 | |
| 31 | ingroup | *Prunus* s.l. | subg. *Padus* | *Prunus hypoleuca* | *Prunus hypoleuca* 12 | PE903 | PRJNA1031385† | SRR28365530 | Minhe, Haidong, Qinghai, China | S.W.Liu#2799 (PE 00995644) | PP648244 | 119262494 | 17889374100 | DGS | 16.66 | |
| 32 | ingroup | *Prunus* s.l. | subg. *Padus* | *Prunus hypoleuca* | *Prunus hypoleuca* 13 | PE908 | PRJNA1031385† | SRR28365528 | Lushi, Sanmenxia, Henan, China | s.coll.#0223 (PE 00772771) | PP648245 | 97336956 | 14600543400 | DGS | 13.60 | |
| 33 | ingroup | *Prunus* s.l. | subg. *Padus* | *Prunus hypoleuca* | *Prunus hypoleuca* 14 | PE910 | PRJNA1031385† | SRR28365526 | Diebu, Gannan, Gansu, China | Bailongjiang Collection Team#1062 (PE 01560561) | PP595867 | 121072690 | 18160903500 | DGS | 16.91 | |
| 34 | ingroup | *Prunus* s.l. | subg. *Padus* | *Prunus hypoleuca* | *Prunus hypoleuca* 15 | PE911 | PRJNA1031385† | SRR28365525 | Wuxi, Chongqing, China | Z.D.Chen et al.#960732 (PE 00995633) | PP648247 | 113839006 | 17075850900 | DGS | 15.90 | |
| 35 | ingroup | *Prunus* s.l. | subg. *Padus* | *Prunus hypoleuca* | *Prunus hypoleuca* 16 | PE912 | PRJNA1031385† | SRR28365524 | Yichang, Hubei, China | R.Long & B.H.Zheng#090078 (PE 01857322) | PP648248 | 120258138 | 18038720700 | DGS | 16.80 | |
| 36 | ingroup | *Prunus* s.l. | subg. *Padus* | *Prunus hypoleuca* | *Prunus hypoleuca* 17 | JR438 | PRJNA1031385† | SRR28365523 | Ningshan, Ankang, Shaanxi, China | L.Zhao#JR438 (WUK) | PP648249 | 160106950 | 24016042500 | DGS | 22.37 | |
| 37 | ingroup | *Prunus* s.l. | subg. *Padus* | *Prunus hypoxantha* (Koehne) J.Wen | *Prunus hypoxantha* 1 | JR372 | PRJNA1031385† | SRR28365536 | Kangding, Sichuan, China | L.Zhao#JR372 (WUK) | PP648239 | 159805816 | 23970872400 | DGS | 22.32 | 2n=4x=32 |
| 38 | ingroup | *Prunus* s.l. | subg. *Padus* | *Prunus hypoxantha* | *Prunus hypoxantha* 2 | PE904 | PRJNA1031385† | SRR28365535 | An, Mianyang, Sichuan, China | D.H.Zhu#3746 (PE 01841240) | PP595866 | 118128168 | 17719225200 | DGS | 16.50 | |
| 39 | ingroup | *Prunus* s.l. | subg. *Padus* | *Prunus hypoxantha* | *Prunus hypoxantha* 3 | WX202 | PRJNA1031385† | SRR28365520 | Emeishan, Leshan, Sichuan, China | L.Zhao#WX202 (WUK) | PP595884 | 143871780 | 21580767000 | DGS | 20.10 | |
| 40 | ingroup | *Prunus* s.l. | subg. *Padus* | *Prunus hypoxantha* | *Prunus hypoxantha* 4 | PE913 | PRJNA1031385† | SRR28365522 | Luding, Ganzi, Sichuan, China | s.coll.#6863 (PE 00995655) | PP648250 | 112851682 | 16927752300 | DGS | 15.77 | |
| 41 | ingroup | *Prunus* s.l. | subg. *Padus* | *Prunus hypoxantha* | *Prunus hypoxantha* 5 | PE914 | PRJNA1031385† | SRR28365521 | Jinyang, Liangshan, Sichuan, China | J.Z.Chuan#3017 (PE 00772868) | PP595885 | 87363340 | 13104501000 | DGS | 12.20 | |
| 42 | ingroup | *Prunus* s.l. | subg. *Padus* | *Prunus serotina* Ehrh. | *Prunus serotina* | WX204 | PRJNA1031385† | SRR28365519 | Washington DC, USA | L.Zhao#WX204 (WUK) | PP595879 | 182664540 | 27399681000 | DGS | 25.52 | |
| 43 | ingroup | *Prunus* s.l. | subg. *Padus* | *Prunus padus* L. | *Prunus padus* | SRR21901493 | PRJNA838254 | SRR21901493 | - | - | PP595875 | 219361394 | 32904209100 | DGS | 30.64 | |
| 44 | ingroup | *Prunus* s.l. | subg. *Padus* | *Prunus brachypoda* Batalin | *Prunus brachypoda* | BOP002669 | PRJNA1031385† | SRR28365517 | China | s.coll.#BOP002669 (PE) | PP595842 | 364695530 | 54704329500 | DGS | 50.95 | |
| 45 | ingroup | *Prunus* s.l. | subg. *Padus* | *Prunus obtusata* Koehne | *Prunus obtusata* | BOP002671 | PRJNA1031385† | SRR28365516 | China | s.coll.#BOP002671 (PE) | PP595843 | 450742030 | 67611304500 | DGS | 62.97 | |
| 46 | ingroup | *Prunus* s.l. | subg. *Padus* | *Prunus buergeriana* Miq. | *Prunus buergeriana* | XYZ026 | PRJNA1031385† | SRR28456441 | Hubei, China | H.Ma#XYZ026 (FDS) | - | 26339348 | 3950902200 | RNA-Seq | 3.68 | |
| 47 | ingroup | *Prunus* s.l. | subg. *Padus* | *Prunus laurocerasus* L. | *Prunus laurocerasus* | WX226 | PRJNA1031385† | SRR28365515 | Rockville, Maryland, USA | L.Zhao#WX227 (WUK) | PP595869 | 202726318 | 30408947700 | DGS | 28.32 | |
| 48 | ingroup | *Prunus* s.l. | subg. *Padus* | *Prunus phaeosticta* Maxim. | *Prunus phaeosticta* | SRR19348248 | PRJNA841240 | SRR19348248 | - | - | - | 58342256 | 8751338400 | DGS | 8.15 | |
| 49 | ingroup | *Prunus* s.l. | subg. *Padus* | *Prunus spinulosa* Siebold & Zucc. | *Prunus spinulosa* | XYZ027 | PRJNA1031385† | SRR28456439 | Wuhan, Hubei, China | H.Ma#XYZ027 (FDS) | - | 23611766 | 3541764900 | RNA-Seq | 3.30 | |
| 50 | ingroup | *Prunus* s.l. | subg. *Padus* | *Prunus caroliniana* (Mill.) Aiton | *Prunus caroliniana* | BOP029039 | PRJNA1031385† | SRR28365514 | USA | s.coll.#BOP029039 (PE) | PP595840 | 690495694 | 1.03574E+11 | DGS | 96.46 | |
| 51 | ingroup | *Prunus* s.l. | subg. *Padus* | *Prunus grisea* (Blume ex Müll.Berol.) Kalkman | *Prunus grisea* | SN510 | PRJNA1031385† | SRR28365513 | Yunnan, China | L.Zhao#SN510 (WUK) | PP595870 | 204088462 | 30613269300 | DGS | 28.51 | |
| 52 | ingroup | *Prunus* s.l. | subg. *Padus* | *Prunus arborea* var. *montana* (Hook.f.) Kalkman | *Prunus arborea* var. *montana* | WX229 | PRJNA1031385† | SRR28365512 | Guangdong, China | L.Zhao#WX229 (WUK) | PP595882 | 165076678 | 24761501700 | DGS | 23.06 | |
| 53 | outgroup | *Prunus* s.l. | subg. *Prunus* | *Prunus ferganensis* (Kostina & Rjabov) Y.Y.Yao ex Y.H.To | *Prunus ferganensis* | BOP002789 | PRJNA1031385† | SRR28365511 | China | s.coll.#BOP002789 (PE) | PP595847 | 383347226 | 57502083900 | DGS | 53.55 | |
| 54 | outgroup | *Prunus* s.l. | subg. *Prunus* | *Prunus davidiana* (Carrière) Franch. | *Prunus davidiana* | SRR12283376 | PRJNA645279 | SRR12283376 | - | - | PP595845 | 88527308 | 13279096200 | DGS | 12.37 | |
| 55 | outgroup | *Prunus* s.l. | subg. *Prunus* | *Prunus mira* Koehne | *Prunus mira* | BOP002793 | PRJNA1031385† | SRR28365510 | China | s.coll.#BOP002793 (PE) | PP595873 | 556074844 | 83411226600 | DGS | 77.68 | |
| 56 | outgroup | *Prunus* s.l. | subg. *Prunus* | *Prunus persica* (L.) Batsch | *Prunus persica* | BOP002795 | PRJNA1031385† | SRR28365509 | China | s.coll.#BOP002795 (PE) | PP595877 | 590282888 | 88542433200 | DGS | 82.46 | |
| 57 | outgroup | *Prunus* s.l. | subg. *Prunus* | *Prunus mume* Siebold & Zucc. | *Prunus mume* | ERR4762270 | PRJEB40984 | ERR4762270 | - | - | PP595874 | 44548438 | 6682265700 | DGS | 6.22 | |
| 58 | outgroup | *Prunus* s.l. | subg. *Prunus* | *Prunus sibirica* L. | *Prunus sibirica* | ERR4656982 | PRJEB40668 | ERR4656982 | - | - | PP595880 | 57034096 | 8555114400 | DGS | 7.97 | |
| 59 | outgroup | *Prunus* s.l. | subg. *Prunus* | *Prunus dasycarpa* Ehrh. | *Prunus dasycarpa* | BOP002749 | PRJNA1031385† | SRR28365508 | China | s.coll.#BOP002749 (PE) | PP595825 | 535138870 | 80270830500 | DGS | 74.76 | |
| 60 | outgroup | *Prunus* s.l. | subg. *Prunus* | *Prunus armeniaca* var. *holosericea* Batalin | *Prunus holosericea* | BOP002750 | PRJNA1031385† | SRR28365506 | China | s.coll.#BOP002750 (PE) | PP595826 | 434259132 | 65138869800 | DGS | 60.67 | |
| 61 | outgroup | *Prunus* s.l. | subg. *Prunus* | *Prunus mandshurica* (Maxim.) Koehne | *Prunus mandshurica* | BOP002747 | PRJNA1031385† | SRR28365505 | - | s.coll.#BOP002747 (PE) | PP595872 | 554555640 | 83183346000 | DGS | 77.47 | |
| 62 | outgroup | *Prunus* s.l. | subg. *Prunus* | *Prunus salicina* Lindl. | *Prunus salicina* | BOP002868 | PRJNA1031385† | SRR28365504 | - | s.coll.#BOP002868 (PE) | PP595878 | 649426630 | 97413994500 | DGS | 90.72 | |
| 63 | outgroup | *Prunus* s.l. | subg. *Prunus* | *Prunus tomentosa* Thunb. | *Prunus tomentosa* | WX216 | PRJNA1031385† | SRR28365502 | Shaanxi, China | L.Zhao#WX216 (WUK) | PP595881 | 207378648 | 31106797200 | DGS | 28.97 | |
| 64 | outgroup | *Prunus* s.l. | subg. *Prunus* | *Prunus japonica* var. *nakaii* (H.Lév.) Rehder | *Prunus japonica* var. *nakaii* | SN503 | PRJNA1031385† | SRR28365501 | Jilin, China | L.Zhao#SN503 (WUK) | PP595868 | 188504662 | 28275699300 | DGS | 26.33 | |
| 65 | outgroup | *Prunus* s.l. | subg. *Prunus* | *Prunus glandulosa* Thunb. | *Prunus glandulosa* | BOP002698 | PRJNA1031385† | SRR28365500 | China | s.coll.#BOP002698 (PE) | PP595831 | 455209710 | 68281456500 | DGS | 63.59 | |
| 66 | outgroup | *Prunus* s.l. | subg. *Prunus* | *Prunus tianshanica* (Pojark.) Shuo Shi | *Prunus tianshanica* | BOP016971 | PRJNA1031385† | SRR28365499 | - | s.coll.#BOP016971 (PE) | PP595836 | 594585840 | 89187876000 | DGS | 83.06 | |
| 67 | outgroup | *Prunus* s.l. | subg. *Prunus* | *Prunus* sp.3 | *Prunus* sp.3 | BOP002519 | PRJNA1031385† | SRR28365490 | - | s.coll.#BOP002519 (PE) | PP595834 | 409723226 | 61458483900 | DGS | 57.24 | |
| 68 | outgroup | *Prunus* s.l. | subg. *Cerasus* | *Prunus* sp.1 | *Prunus* sp.1 | BOP002589 | PRJNA1031385† | SRR28365498 | - | s.coll.#BOP002589 (PE) | PP595837 | 492634634 | 73895195100 | DGS | 68.82 | |
| 69 | outgroup | *Prunus* s.l. | subg. *Cerasus* | *Prunus* sp.2 | *Prunus* sp.2 | BOP002869 | PRJNA1031385† | SRR28365503 | - | s.coll.#BOP002869 (PE) | PP595883 | 576583146 | 86487471900 | DGS | 80.55 | |
| 70 | outgroup | *Prunus* s.l. | subg. *Cerasus* | *Prunus discadenia* Koehne | *Prunus discadenia* | SRR12927899 | PRJNA671803 | SRR12927899 | - | - | PP595846 | 65331840 | 9799776000 | DGS | 9.13 | |
| 71 | outgroup | *Prunus* s.l. | subg. *Cerasus* | *Prunus avium* L. | *Prunus avium* | BOP002693 | PRJNA1031385† | SRR28365497 | - | s.coll.#BOP002693 (PE) | PP595827 | 437085744 | 65562861600 | DGS | 61.06 | |
| 72 | outgroup | *Prunus* s.l. | subg. *Cerasus* | *Prunus clarofolia* Schneider | *Prunus clarofolia* | BOP002703 | PRJNA1031385† | SRR28365495 | China | s.coll.#BOP002703 (PE) | PP595828 | 433687596 | 65053139400 | DGS | 60.59 | |
| 73 | outgroup | *Prunus* s.l. | subg. *Cerasus* | *Prunus cyclamina* Koehne | *Prunus cyclamina* | BOP002628 | PRJNA1031385† | SRR28365494 | China | s.coll.#BOP002628 (PE) | PP595829 | 428033760 | 64205064000 | DGS | 59.80 | |
| 74 | outgroup | *Prunus* s.l. | subg. *Cerasus* | *Prunus dielsiana* C.K.Schneid. | *Prunus dielsiana* | BOP002629 | PRJNA1031385† | SRR28365493 | China | s.coll.#BOP002629 (PE) | PP595830 | 453256438 | 67988465700 | DGS | 63.32 | |
| 75 | outgroup | *Prunus* s.l. | subg. *Cerasus* | *Prunus maximowiczii* Rupr. | *Prunus maximowiczii* | BOP016576 | PRJNA1031385† | SRR28365492 | - | s.coll.#BOP016576 (PE) | PP595832 | 614396216 | 92159432400 | DGS | 85.83 | |
| 76 | outgroup | *Prunus* s.l. | subg. *Cerasus* | *Prunus pseudocerasus* Lindl. | *Prunus pseudocerasus* | BOP002626 | PRJNA1031385† | SRR28365491 | China | s.coll.#BOP002626 (PE) | PP595833 | 477165342 | 71574801300 | DGS | 66.66 | |
| 77 | outgroup | *Prunus* s.l. | subg. *Cerasus* | *Prunus* × *subhirtella* Miq. | *Prunus* × *subhirtella* | BOP002663 | PRJNA1031385† | SRR28365489 | USA | s.coll.#BOP002663 (PE) | PP595835 | 389710634 | 58456595100 | DGS | 54.44 | |
| 78 | outgroup | *Prunus* s.l. | subg. *Cerasus* | *Prunus* × *yedoensis* Matsum. | *Prunus* × *yedoensis* | BOP002623 | PRJNA1031385† | SRR28365488 | USA | s.coll.#BOP002623 (PE) | PP595838 | 389344286 | 58401642900 | DGS | 54.39 | |
| 79 | outgroup | *Prunus* s.l. | subg. *Cerasus* | *Prunus yunnanensis* Franch. | *Prunus yunnanensis* | BOP002590 | PRJNA1031385† | SRR28365487 | China | s.coll.#BOP002590 (PE) | PP595839 | 533773116 | 80065967400 | DGS | 74.57 | |
| 80 | outgroup | *Prunus* s.l. | subg. *Cerasus* | *Prunus maackii* Rupr. | *Prunus maackii* | SRR18715741 | PRJNA788510 | SRR18715741 | - | - | - | 63808228 | 9571234200 | RNA-Seq | 8.91 | |
| 81 | outgroup | *Prunus* s.l. | subg. *Cerasus* | *Prunus cerasoides* D.Don | *Prunus cerasoides* | BOP002855 | PRJNA1031385† | SRR28365486 | China | s.coll.#BOP002855 (PE) | PP595871 | 610420224 | 91563033600 | DGS | 85.27 | |
| 82 | outgroup | *Prunus* s.l. | subg. *Cerasus* | *Prunus itosakura* Siebold | *Prunus itosakura* | BOP204473 | PRJNA1031385† | SRR28365484 | Japan | s.coll.#BOP204473 (PE) | PP595876 | 633874790 | 95081218500 | DGS | 88.55 | |
| 83 | outgroup | *Exochorda* | | *Exochorda racemosa* (Lindl.) Rehder | *Exochorda racemosa* | XJ006 | PRJNA1031385† | SRR28456438 | USA | H.Ma#XJ006 (FDS) | - | 24947790 | 3742168500 | RNA-seq | 3.49 | |
| 84 | outgroup | *Oemleria* | | *Oemleria cerasiformis* (Torr. & A.Gray ex Hook. & Arn.) J.` | *Oemleria cerasiformis* | WYQ11 | PRJNA1031385† | SRR28456437 | USA | H.Ma#WYQ11 (FDS) | - | 49307926 | 7396188900 | RNA-Seq | 6.89 | |
| 85 | outgroup | *Prinsepia* | | *Prinsepia uniflora* Batalin | *Prinsepia uniflora* | WX231 | PRJNA1031385† | SRR28365483 | Xianyang, Shaanxi, China | L.Zhao#WX231 (WUK) | PP595844 | 165828450 | 24874267500 | DGS | 23.17 | |
| 86 | outgroup | *Gillenia* | | *Gillenia stipulata* (Muhl. ex Willd.) Nutt. | *Gillenia stipulata* | SRR3166477 | PRJNA293974 | SRR3166477 | - | - | - | 88956446 | 13343466900 | DGS | 12.43 | |
| 87 | outgroup | *Coleogyne* | | *Coleogyne ramosissima* Torr. | *Coleogyne ramosissima* | HM918 | PRJNA1031385† | SRR28456436 | USA | H.Ma#HM918 (FDS) | - | 157688268 | 23653240200 | RNA-Seq | 22.03 | |
| 88 | outgroup | *Kerria* | | *Kerria japonica* (L.) DC. | *Kerria japonica* | ERR2040423 | PRJEB3060 | ERR204023 | - | - | - | 21498478 | 3224771700 | RNA-Seq | 3.00 | |
| 89 | outgroup | *Neviusia* | | *Neviusia alabamensis* A.Gray | *Neviusia alabamensis* | HM961 | PRJNA1031385† | SRR28456435 | National Arboretum, Washington DC, USA | H.Ma#HM961 (FDS) | - | 169749288 | 25462393200 | RNA-Seq | 23.71 | |
| 90 | outgroup | *Rhodotypos* | | *Rhodotypos scandens* (Thunb.) Makino | *Rhodotypos scandens* | HM935 | PRJNA1031385† | SRR28456434 | National Arboretum, Washington DC, USA | H.Ma#HM935 (FDS) | - | 217413144 | 32611971600 | RNA-Seq | 30.37 | |
| 91 | outgroup | *Lyonothamnus* | | *Lyonothamnus floribundus* A.Gray | *Lyonothamnus floribundus* | WYQ-9A | PRJNA1031385† | SRR28456433 | USA | H.Ma#WYQ-9A (FDS) | - | 36615026 | 5492253900 | RNA-Seq | 5.12 | |
| 92 | outgroup | *Amelanchier* | | *Amelanchier alnifolia* (Nutt.) Nutt. ex M.Roem. | *Amelanchier alnifolia* | HM976 | PRJNA1031385† | SRR15691199 | Pennsylvania, USA | H.Ma#HM976 (FDS) | - | 119122776 | 17868416400 | RNA-Seq | 16.64 | |
| 93 | outgroup | *Crataegus* | | *Crataegus cuneata* Siebold & Zucc. | *Crataegus cuneata* | XJC | PRJNA1031385† | SRR28456451 | Hubei, China | H.Ma#XJC (FDS) | - | 27080646 | 4062096900 | RNA-Seq | 3.78 | |
| 94 | outgroup | *Kageneckia* | | *Kageneckia oblonga* Ruiz & Pav. | *Kageneckia oblonga* | SRR3157483 | PRJNA293974 | SRR3157483 | California, USA | H.Ma#WUQ-8 (FDS) | - | 201338796 | 30200819400 | DGS | 28.13 | |
| 95 | outgroup | *Malacomeles* | | *Malacomeles denticulata* (Kunth) G.N.Jones | *Malacomeles denticulata* | WYQ-8 | PRJNA1031385† | SRR15691197 | California, USA | H.Ma#WYQ-9 (FDS) | - | 37202332 | 5580349800 | RNA-Seq | 5.20 | |
| 96 | outgroup | *Malus* | | *Malus asiatica* Nakai | *Malus asiatica* | BOP002803 | PRJNA1031385† | SRR28365482 | China | s.coll.#BOP002804 (PE) | PP595841 | 602407378 | 90361106700 | DGS | 84.16 | |
| 97 | outgroup | *Malus* | | *Malus baccata* (L.) Borkh. | *Malus baccata* | WYQ-14A | PRJNA1031385† | SRR15691197 | Hubei, China | H.Ma#WYQ-15A (FDS) | - | 28368454 | 4255268100 | RNA-Seq | 3.96 | |
| 98 | outgroup | *Pyrus* | | *Pyrus betulifolia* Bunge | *Pyrus betulifolia* | SRR7135592 | PRJNA381668 | SRR7135592 | Hubei, China | - | - | 69785618 | 10467842700 | DGS | 9.75 | |
| 99 | outgroup | *Rhaphiolepis* | | *Rhaphiolepis umbellata* (Thunb.) Makino | *Rhaphiolepis umbellata* | HM1227 | PRJNA1031385† | SRR28456450 | Beijing, China | H.Ma#HM1227 (FDS) | - | 79238440 | 11885766000 | RNA-Seq | 11.07 | |
| 100 | outgroup | *Sorbus* | | *Sorbus aucuparia* L. | *Sorbus aucuparia* | Z1051 | PRJNA1031385† | SRR28456449 | - | H.Ma#Z1051 (FDS) | - | 182441736 | 27366260400 | RNA-Seq | 25.49 | |
| 101 | outgroup | *Vauquelinia* | | *Vauquelinia californica* (Torr.) Sarg. | *Vauquelinia californica* | SRR3130625 | PRJNA293974 | SRR3130625 | - | - | - | 141008986 | 21151347900 | DGS | 19.70 | |
| 102 | outgroup | *Photinia* | | *Photinia* sp. | *Photinia* sp. | SRR6434988 | PRJNA422719 | SRR6434988 | - | - | - | 64078640 | 9611796000 | DGS | 8.95 | |
| 103 | outgroup | *Neillia* | | *Neillia incisa* (Thunb.) S.H.Oh | *Neillia incisa* | SRR13065771 | PRJNA678476 | SRR13065771 | - | - | - | 41003194 | 6150479100 | DGS | 5.73 | |
| 104 | outgroup | *Physocarpus* | | *Physocarpus amurensis* (Maxim.) Maxim. | *Physocarpus amurensis* | WX230 | PRJNA1031385† | SRR28365481 | Xianyang, Shaanxi, China | L.Zhao#WX230 (WUK) | PP648233 | 165838140 | 24875721000 | DGS | 23.17 | |
| 105 | outgroup | *Stephanandra* | | *Neillia tanakae* Franch. & Sav. | *Neillia tanakae* | HM803 | PRJNA1031385† | SRR28456448 | Hubei, China | H.Ma#HM803 (FDS) | - | 25738654 | 3860798100 | RNA-Seq | 3.60 | |
| 106 | outgroup | *Adenostoma* | | *Adenostoma fasciculatum* Hook. & Arn. | *Adenostoma fasciculatum* | WYQ-2B | PRJNA1031385† | SRR28456447 | USA | H.Ma#WYQ-2B (FDS) | - | 37202332 | 5580349800 | RNA-Seq | 5.20 | |
| 107 | outgroup | *Chamaebatiaria* | | *Chamaebatiaria millefolium* (Torr.) Maxim. | *Chamaebatiaria millefolium* | HM1264 | PRJNA1031385† | SRR28456446 | USA | H.Ma#HM1264 (FDS) | - | 221453344 | 33218001600 | RNA-Seq | 30.94 | |
| 108 | outgroup | *Sorbaria* | | *Sorbaria sorbifolia* (L.) A.Braun | *Sorbaria sorbifolia* | HM1085 | PRJNA1031385† | SRR28456445 | Pennsylvania, USA | H.Ma#HM1085 (FDS) | - | 39522674 | 5928401100 | RNA-Seq | 5.52 | |
| 109 | outgroup | *Aruncus* | | *Aruncus dioicus* (Walter) Fernald | *Aruncus dioicus* | HM1078 | PRJNA1031385† | SRR28456444 | Hortus Botanicus Amsterdam, Netherlands | H.Ma#HM1078 (FDS) | - | 48068974 | 7210346100 | RNA-Seq | 6.72 | |
| 110 | outgroup | *Holodiscus* | | *Holodiscus discolor* (Pursh) Maxim. | *Holodiscus discolor* | HM832 | PRJNA1031385† | SRR28456443 | Hortus Botanicus Amsterdam, Netherlands | H.Ma#HM832 (FDS) | - | 26342326 | 3951348900 | RNA-Seq | 3.68 | |
| 111 | outgroup | *Petrophytum* | | *Petrophytum caespitosum* (Nutt.) Rydb. | *Petrophytum caespitosum* | HM1385 | PRJNA1031385† | SRR28456442 | Bonn, Germany | H.Ma#HM1385 (FDS) | - | 27175908 | 4076386200 | RNA-Seq | 3.80 | |
| 112 | outgroup | *Sibiraea* | | *Sibiraea laevigata* (L.) Maxim. | *Sibiraea laevigata* | HM1225 | PRJNA1031385† | SRR28456440 | Inner Mongolia, China | H.Ma#HM1225 (FDS) | - | 21529898 | 3229484700 | RNA-Seq | 3.01 | |
| 113 | outgroup | *Spiraea* | | *Spiraea thunbergii* Siebold ex Blume | *Spiraea thunbergii* | SRR19090745 | PRJNA835237 | SRR19090745 | - | - | PP595886 | 46992320 | 7048848000 | DGS | 6.56 | |
| 114 | outgroup | *Rosa* | | *Rosa rugosa* Thunb. | *Rosa rugosa* | BOP010536 | PRJNA1031385† | SRR28365480 | China | s.coll.#BOP010536 (PE) | PP648251 | 670455110 | 100568266500 | DGS | 93.66 | |
