## Supplemental Table 2 for "Unveiling allopolyploidization-driven genome duplications through progressive analysis of deep genome skimming data"

| Name | NumReads | ReadsMapped | PctOnTarget | GenesMapped | GenesWithContigs | GenesWithSeqs | GenesAt25pct | GenesAt50pct | GenesAt75pct | GenesAt150pct | ParalogWarningsLong | ParalogWarningsDepth | GenesWithoutStitchedContigs | GenesWithStitchedContigs | GenesWithStitchedContigsSkipped | GenesWithChimeraWarning |
|---|---|---|---|---|---|---|---|---|---|---|---|---|---|---|---|---|
| *Adenostoma fasciculatum* | 37202332 | 2143652 | 5.8 | 1800 | 1739 | 1637 | 1591 | 1555 | 1491 | 2 | 16 | 25 | 1418 | 219 | 0 | 0 |
| *Amelanchier alnifolia* | 119122776 | 3292686 | 2.8 | 1792 | 1652 | 1502 | 1424 | 1314 | 1180 | 0 | 82 | 290 | 765 | 738 | 0 | 0 |
| *Aruncus dioicus* | 48068974 | 2525245 | 5.3 | 1800 | 1734 | 1669 | 1653 | 1625 | 1586 | 0 | 7 | 11 | 1471 | 198 | 0 | 0 |
| *Chamaebatiaria millefolium* | 221453344 | 11665973 | 5.3 | 1806 | 1799 | 1784 | 1774 | 1753 | 1689 | 1 | 22 | 72 | 1250 | 534 | 0 | 0 |
| *Coleogyne ramosissima* | 157688268 | 9250175 | 5.9 | 1799 | 1783 | 1714 | 1673 | 1617 | 1543 | 0 | 11 | 43 | 1114 | 600 | 0 | 0 |
| *Crataegus cuneata* | 27080646 | 1704805 | 6.3 | 1805 | 1743 | 1673 | 1628 | 1585 | 1526 | 4 | 304 | 459 | 1311 | 362 | 0 | 0 |
| *Exochorda racemosa* | 24947790 | 1689427 | 6.8 | 1805 | 1750 | 1678 | 1634 | 1596 | 1549 | 0 | 15 | 25 | 1461 | 217 | 0 | 0 |
| *Gillenia stipulata* | 88956446 | 629988 | 0.7 | 1806 | 1794 | 1790 | 1769 | 1636 | 1171 | 0 | 0 | 7 | 173 | 1617 | 0 | 0 |
| *Holodiscus discolor* | 26342326 | 1599691 | 6.1 | 1803 | 1754 | 1674 | 1635 | 1598 | 1537 | 4 | 259 | 369 | 1353 | 321 | 0 | 0 |
| *Kageneckia oblonga* | 201338796 | 1714355 | 0.9 | 1806 | 1803 | 1800 | 1799 | 1793 | 1752 | 5 | 333 | 798 | 1002 | 798 | 0 | 0 |
| *Kerria japonica* | 21498478 | 1116663 | 5.2 | 1780 | 1501 | 1496 | 1454 | 1338 | 1107 | 0 | 1 | 2 | 822 | 674 | 0 | 0 |
| *Lyonothamnus floribundus* | 36615026 | 1686752 | 4.6 | 1798 | 1580 | 1355 | 1262 | 1173 | 1106 | 3 | 240 | 342 | 994 | 361 | 0 | 0 |
| *Malacomeles denticulata* | 28368454 | 1643154 | 5.8 | 1802 | 1697 | 1573 | 1529 | 1498 | 1454 | 5 | 267 | 457 | 1222 | 351 | 0 | 0 |
| *Malus asiatica* | 602407378 | 5217301 | 0.9 | 1806 | 1806 | 1805 | 1805 | 1800 | 1745 | 5 | 216 | 880 | 579 | 1226 | 0 | 0 |
| *Malus baccata* | 71999602 | 1147822 | 1.6 | 1806 | 1789 | 1787 | 1785 | 1770 | 1624 | 3 | 97 | 563 | 472 | 1315 | 0 | 0 |
| *Neillia incisa* | 41003194 | 683057 | 1.7 | 1805 | 1801 | 1800 | 1800 | 1795 | 1756 | 2 | 19 | 52 | 847 | 953 | 0 | 0 |
| *Neillia tanakae* | 25738654 | 1408789 | 5.5 | 1795 | 1693 | 1598 | 1555 | 1519 | 1481 | 0 | 12 | 18 | 1431 | 167 | 0 | 0 |
| *Neviusia alabamensis* | 169749288 | 7437911 | 4.4 | 1804 | 1795 | 1768 | 1736 | 1696 | 1626 | 0 | 21 | 51 | 1285 | 484 | 0 | 0 |
| *Oemleria cerasiformis* | 49307926 | 1917053 | 3.9 | 1797 | 1641 | 1565 | 1508 | 1464 | 1400 | 0 | 5 | 12 | 1326 | 240 | 0 | 0 |
| *Petrophytum caespitosum* | 27175908 | 1339493 | 4.9 | 1805 | 1737 | 1638 | 1602 | 1558 | 1478 | 1 | 4 | 5 | 1439 | 199 | 0 | 0 |
| *Photinia* sp. | 64078640 | 1640704 | 2.6 | 1803 | 1597 | 1438 | 1350 | 1198 | 865 | 1 | 45 | 145 | 930 | 508 | 0 | 0 |
| *Physocarpus amurensis* | 165838140 | 3157354 | 1.9 | 1806 | 1803 | 1803 | 1803 | 1802 | 1787 | 1 | 37 | 57 | 1338 | 465 | 0 | 0 |
| *Prinsepia uniflora* | 165828450 | 833181 | 0.5 | 1806 | 1767 | 1761 | 1752 | 1694 | 1469 | 2 | 76 | 244 | 674 | 1087 | 0 | 0 |
| *Prunus* × *subhirtella* | 389710634 | 6653273 | 1.7 | 1806 | 1806 | 1803 | 1803 | 1803 | 1803 | 1 | 20 | 64 | 1051 | 752 | 0 | 0 |
| *Prunus* × *yedoensis* | 389344286 | 6632641 | 1.7 | 1806 | 1806 | 1803 | 1803 | 1803 | 1803 | 2 | 14 | 68 | 729 | 1074 | 0 | 0 |
| *Prunus arborea* var. *montana* | 165076678 | 3221010 | 2 | 1806 | 1805 | 1804 | 1804 | 1803 | 1801 | 11 | 861 | 1391 | 1250 | 554 | 0 | 0 |
| *Prunus avium* | 437085744 | 6056219 | 1.4 | 1806 | 1806 | 1806 | 1806 | 1806 | 1806 | 2 | 20 | 60 | 1013 | 793 | 0 | 0 |
| *Prunus brachypoda* | 364695530 | 6744094 | 1.8 | 1806 | 1806 | 1803 | 1803 | 1803 | 1803 | 3 | 75 | 590 | 295 | 1508 | 0 | 0 |
| *Prunus buergeriana* | 26339348 | 2223704 | 8.4 | 1806 | 1769 | 1704 | 1666 | 1638 | 1606 | 1 | 197 | 469 | 1101 | 603 | 0 | 0 |
| *Prunus caroliniana* | 690495694 | 9813890 | 1.4 | 1806 | 1804 | 1801 | 1801 | 1799 | 1796 | 5 | 408 | 1250 | 663 | 1138 | 0 | 0 |
| *Prunus cerasoides* | 610420224 | 10635140 | 1.7 | 1806 | 1806 | 1804 | 1803 | 1802 | 1802 | 2 | 17 | 58 | 1060 | 744 | 0 | 0 |
| *Prunus clarofolia* | 433687596 | 5883535 | 1.4 | 1806 | 1806 | 1806 | 1806 | 1806 | 1805 | 1 | 18 | 56 | 1001 | 805 | 0 | 0 |
| *Prunus cyclamina* | 428033760 | 7035109 | 1.6 | 1806 | 1806 | 1803 | 1803 | 1803 | 1802 | 1 | 16 | 64 | 968 | 835 | 0 | 0 |
| *Prunus dasycarpa* | 535138870 | 10142426 | 1.9 | 1806 | 1806 | 1804 | 1804 | 1804 | 1803 | 1 | 16 | 81 | 640 | 1164 | 0 | 0 |
| *Prunus davidiana* | 88527308 | 2209965 | 2.5 | 1806 | 1806 | 1803 | 1803 | 1803 | 1802 | 1 | 28 | 74 | 1028 | 775 | 0 | 0 |
| *Prunus dielsiana* | 453256438 | 7247975 | 1.6 | 1806 | 1806 | 1806 | 1806 | 1806 | 1806 | 1 | 19 | 68 | 988 | 818 | 0 | 0 |
| *Prunus discadenia* | 65331840 | 1708206 | 2.6 | 1806 | 1806 | 1802 | 1802 | 1802 | 1801 | 1 | 21 | 73 | 1190 | 612 | 0 | 0 |
| *Prunus ferganensis* | 383347226 | 7448237 | 1.9 | 1806 | 1806 | 1805 | 1805 | 1805 | 1805 | 1 | 26 | 62 | 1121 | 684 | 0 | 0 |
| *Prunus fujianensis* 1 | 166886168 | 3788165 | 2.3 | 1806 | 1806 | 1805 | 1805 | 1805 | 1805 | 8 | 586 | 1362 | 1023 | 782 | 0 | 0 |
| *Prunus fujianensis* 2 | 163167226 | 3640650 | 2.2 | 1806 | 1806 | 1803 | 1803 | 1803 | 1802 | 11 | 585 | 1377 | 1004 | 799 | 0 | 0 |
| *Prunus fujianensis* 3 | 83808988 | 2035664 | 2.4 | 1806 | 1806 | 1803 | 1803 | 1803 | 1800 | 6 | 237 | 1086 | 560 | 1243 | 0 | 0 |
| *Prunus fujianensis* 4 | 121863964 | 3025154 | 2.5 | 1806 | 1806 | 1804 | 1804 | 1803 | 1802 | 6 | 356 | 1284 | 624 | 1180 | 0 | 0 |
| *Prunus fujianensis* 5 | 89685070 | 2140100 | 2.4 | 1806 | 1806 | 1805 | 1805 | 1804 | 1801 | 4 | 273 | 1174 | 562 | 1243 | 0 | 0 |
| *Prunus fujianensis* 6 | 97483868 | 2337116 | 2.4 | 1806 | 1806 | 1802 | 1802 | 1802 | 1801 | 6 | 397 | 1212 | 779 | 1023 | 0 | 0 |
| *Prunus fujianensis* 7 | 132721316 | 3408507 | 2.6 | 1806 | 1806 | 1802 | 1802 | 1801 | 1799 | 5 | 402 | 1294 | 635 | 1167 | 0 | 0 |
| *Prunus glandulosa* | 455209710 | 8945784 | 2 | 1806 | 1806 | 1803 | 1803 | 1803 | 1803 | 1 | 27 | 55 | 1171 | 632 | 0 | 0 |
| *Prunus gongshanensis* 1 | 70939130 | 1550849 | 2.2 | 1806 | 1806 | 1802 | 1802 | 1802 | 1801 | 3 | 372 | 1219 | 733 | 1069 | 0 | 0 |
| *Prunus gongshanensis* 2 | 67170668 | 1453247 | 2.2 | 1806 | 1806 | 1803 | 1803 | 1803 | 1802 | 6 | 344 | 1156 | 707 | 1096 | 0 | 0 |
| *Prunus gongshanensis* 3 | 149826116 | 3765141 | 2.5 | 1806 | 1806 | 1804 | 1804 | 1803 | 1802 | 5 | 345 | 1308 | 583 | 1221 | 0 | 0 |
| *Prunus gongshanensis* 4 | 182209552 | 5374799 | 2.9 | 1806 | 1806 | 1802 | 1802 | 1801 | 1799 | 4 | 170 | 1111 | 291 | 1511 | 0 | 0 |
| *Prunus gongshanensis* 5 | 158371348 | 3365815 | 2.1 | 1806 | 1805 | 1804 | 1801 | 1746 | 1427 | 0 | 0 | 2 | 60 | 1744 | 0 | 0 |
| *Prunus gongshanensis* 6 | 174192216 | 4845150 | 2.8 | 1806 | 1806 | 1802 | 1802 | 1802 | 1801 | 3 | 190 | 1155 | 357 | 1445 | 0 | 0 |
| *Prunus gongshanensis* 7 | 481515052 | 10033778 | 2.1 | 1806 | 1806 | 1803 | 1803 | 1802 | 1801 | 5 | 563 | 1409 | 821 | 983 | 0 | 0 |
| *Prunus grisea* | 204088462 | 3477806 | 1.7 | 1806 | 1805 | 1803 | 1803 | 1800 | 1799 | 9 | 783 | 1386 | 1104 | 699 | 0 | 0 |
| *Prunus himalayana* 1 | 128548808 | 3431901 | 2.7 | 1806 | 1806 | 1804 | 1804 | 1802 | 1790 | 1 | 17 | 417 | 169 | 1635 | 0 | 0 |
| *Prunus himalayana* 2 | 245486784 | 7373599 | 3 | 1806 | 1806 | 1802 | 1802 | 1800 | 1799 | 4 | 248 | 1217 | 381 | 1422 | 0 | 0 |
| *Prunus himalayana* 3 | 261661096 | 7844323 | 3 | 1806 | 1806 | 1802 | 1802 | 1802 | 1799 | 5 | 235 | 1205 | 359 | 1444 | 0 | 0 |
| *Prunus himalayana* 4 | 127215638 | 2724144 | 2.1 | 1806 | 1806 | 1804 | 1804 | 1803 | 1800 | 5 | 494 | 1312 | 922 | 882 | 0 | 0 |
| *Prunus himalayana* 5 | 178337194 | 3452972 | 1.9 | 1806 | 1806 | 1805 | 1805 | 1804 | 1804 | 6 | 594 | 1384 | 1012 | 793 | 0 | 0 |
| *Prunus holosericea* | 434259132 | 8445475 | 1.9 | 1806 | 1806 | 1804 | 1804 | 1804 | 1804 | 1 | 17 | 52 | 1074 | 730 | 0 | 0 |
| *Prunus hypoleuca* 1 | 175659460 | 3948123 | 2.2 | 1806 | 1806 | 1803 | 1803 | 1802 | 1801 | 6 | 597 | 1390 | 1016 | 787 | 0 | 0 |
| *Prunus hypoleuca* 10 | 119578966 | 2718152 | 2.3 | 1806 | 1806 | 1805 | 1804 | 1804 | 1801 | 3 | 408 | 1366 | 638 | 1167 | 0 | 0 |
| *Prunus hypoleuca* 11 | 129489286 | 3041047 | 2.3 | 1806 | 1806 | 1801 | 1801 | 1800 | 1799 | 6 | 464 | 1387 | 699 | 1102 | 0 | 0 |
| *Prunus hypoleuca* 12 | 119262494 | 3325824 | 2.8 | 1806 | 1806 | 1801 | 1801 | 1801 | 1801 | 4 | 361 | 1305 | 605 | 1196 | 0 | 0 |
| *Prunus hypoleuca* 13 | 97336956 | 2962435 | 3 | 1806 | 1806 | 1800 | 1799 | 1799 | 1799 | 7 | 224 | 1124 | 451 | 1349 | 0 | 0 |
| *Prunus hypoleuca* 14 | 121072690 | 2768579 | 2.3 | 1806 | 1806 | 1800 | 1799 | 1799 | 1797 | 8 | 478 | 1353 | 766 | 1034 | 0 | 0 |
| *Prunus hypoleuca* 15 | 113839006 | 2777340 | 2.4 | 1806 | 1806 | 1801 | 1801 | 1800 | 1797 | 5 | 438 | 1352 | 686 | 1115 | 0 | 0 |
| *Prunus hypoleuca* 16 | 120258138 | 2797293 | 2.3 | 1806 | 1806 | 1802 | 1802 | 1802 | 1802 | 7 | 401 | 1334 | 666 | 1136 | 0 | 0 |
| *Prunus hypoleuca* 17 | 160106950 | 3577427 | 2.2 | 1806 | 1806 | 1805 | 1805 | 1804 | 1803 | 6 | 537 | 1338 | 946 | 859 | 0 | 0 |
| *Prunus hypoleuca* 2 | 176922850 | 4135784 | 2.3 | 1806 | 1806 | 1805 | 1805 | 1804 | 1803 | 3 | 595 | 1395 | 979 | 826 | 0 | 0 |
| *Prunus hypoleuca* 3 | 163680606 | 4562670 | 2.8 | 1806 | 1806 | 1800 | 1800 | 1799 | 1796 | 2 | 58 | 712 | 201 | 1599 | 0 | 0 |
| *Prunus hypoleuca* 4 | 438944908 | 10497166 | 2.4 | 1806 | 1806 | 1803 | 1803 | 1803 | 1803 | 6 | 667 | 1424 | 985 | 818 | 0 | 0 |
| *Prunus hypoleuca* 5 | 125411946 | 1000135 | 0.8 | 1806 | 1777 | 1773 | 1756 | 1716 | 1537 | 0 | 4 | 44 | 455 | 1318 | 0 | 0 |
| *Prunus hypoleuca* 6 | 79566196 | 3796594 | 4.8 | 1806 | 1806 | 1801 | 1801 | 1799 | 1782 | 1 | 44 | 447 | 270 | 1531 | 0 | 0 |
| *Prunus hypoleuca* 7 | 79805998 | 2312383 | 2.9 | 1806 | 1806 | 1802 | 1802 | 1799 | 1783 | 1 | 32 | 410 | 299 | 1503 | 0 | 0 |
| *Prunus hypoleuca* 8 | 116234522 | 3312318 | 2.8 | 1806 | 1806 | 1802 | 1802 | 1801 | 1799 | 5 | 293 | 1235 | 491 | 1312 | 0 | 0 |
| *Prunus hypoleuca* 9 | 161338728 | 3773566 | 2.3 | 1806 | 1806 | 1802 | 1802 | 1801 | 1800 | 5 | 560 | 1357 | 955 | 847 | 0 | 0 |
| *Prunus hypoxantha* 1 | 159805816 | 3722409 | 2.3 | 1806 | 1806 | 1803 | 1803 | 1803 | 1802 | 3 | 574 | 1380 | 953 | 850 | 0 | 0 |
| *Prunus hypoxantha* 2 | 118128168 | 2842726 | 2.4 | 1806 | 1806 | 1803 | 1803 | 1803 | 1802 | 5 | 422 | 1363 | 664 | 1139 | 0 | 0 |
| *Prunus hypoxantha* 3 | 143871780 | 3275523 | 2.3 | 1806 | 1806 | 1804 | 1804 | 1803 | 1803 | 5 | 568 | 1359 | 963 | 841 | 0 | 0 |
| *Prunus hypoxantha* 4 | 112851682 | 3512217 | 3.1 | 1806 | 1806 | 1800 | 1800 | 1799 | 1798 | 4 | 251 | 1209 | 460 | 1340 | 0 | 0 |
| *Prunus hypoxantha* 5 | 87363340 | 3033001 | 3.5 | 1806 | 1806 | 1803 | 1803 | 1802 | 1800 | 5 | 106 | 814 | 361 | 1442 | 0 | 0 |
| *Prunus itosakura* | 633874790 | 12338425 | 1.9 | 1806 | 1806 | 1802 | 1802 | 1802 | 1802 | 1 | 19 | 63 | 1071 | 731 | 0 | 0 |
| *Prunus japonica* var. *nakaii* | 188504662 | 4743403 | 2.5 | 1806 | 1806 | 1805 | 1805 | 1805 | 1805 | 1 | 32 | 71 | 1197 | 608 | 0 | 0 |
| *Prunus laurocerasus* | 202726318 | 4186312 | 2.1 | 1806 | 1806 | 1797 | 1797 | 1796 | 1796 | 2 | 194 | 1120 | 483 | 1314 | 0 | 0 |
| *Prunus maackii* | 63808228 | 4925392 | 7.7 | 1805 | 1784 | 1753 | 1731 | 1694 | 1665 | 1 | 23 | 62 | 1458 | 295 | 0 | 0 |
| *Prunus mandshurica* | 554555640 | 10575841 | 1.9 | 1806 | 1806 | 1803 | 1803 | 1803 | 1802 | 1 | 19 | 54 | 1023 | 780 | 0 | 0 |
| *Prunus maximowiczii* | 614396216 | 12319693 | 2 | 1806 | 1806 | 1802 | 1802 | 1802 | 1802 | 1 | 14 | 66 | 941 | 861 | 0 | 0 |
| *Prunus mira* | 556074844 | 9583668 | 1.7 | 1806 | 1806 | 1805 | 1805 | 1804 | 1804 | 1 | 23 | 60 | 1195 | 610 | 0 | 0 |
| *Prunus mume* | 44548438 | 1153455 | 2.6 | 1806 | 1806 | 1804 | 1804 | 1803 | 1802 | 1 | 22 | 63 | 1204 | 600 | 0 | 0 |
| *Prunus obtusata* | 450742030 | 8340917 | 1.9 | 1806 | 1806 | 1805 | 1805 | 1805 | 1805 | 3 | 90 | 603 | 300 | 1505 | 0 | 0 |
| *Prunus padus* | 219361394 | 5291681 | 2.4 | 1806 | 1806 | 1799 | 1799 | 1798 | 1798 | 5 | 512 | 1387 | 854 | 946 | 0 | 0 |
| *Prunus persica* | 590282888 | 10951287 | 1.9 | 1806 | 1806 | 1805 | 1805 | 1805 | 1805 | 1 | 29 | 65 | 1211 | 594 | 0 | 0 |
| *Prunus phaeosticta* | 58342256 | 1202500 | 2.1 | 1806 | 1806 | 1804 | 1804 | 1803 | 1802 | 7 | 570 | 1289 | 962 | 842 | 0 | 0 |
| *Prunus pseudocerasus* | 477165342 | 7720132 | 1.6 | 1806 | 1806 | 1804 | 1804 | 1803 | 1803 | 1 | 13 | 57 | 962 | 842 | 0 | 0 |
| *Prunus salicina* | 649426630 | 12062149 | 1.9 | 1806 | 1806 | 1805 | 1805 | 1805 | 1805 | 2 | 18 | 57 | 969 | 836 | 0 | 0 |
| *Prunus serotina* | 182664540 | 4527574 | 2.5 | 1806 | 1806 | 1804 | 1804 | 1804 | 1804 | 8 | 728 | 1503 | 1061 | 743 | 0 | 0 |
| *Prunus sibirica* | 57034096 | 2487852 | 4.4 | 1806 | 1806 | 1804 | 1804 | 1804 | 1804 | 1 | 33 | 88 | 1164 | 640 | 0 | 0 |
| *Prunus* sp.1 | 492634634 | 9735659 | 2 | 1806 | 1806 | 1800 | 1800 | 1800 | 1799 | 2 | 11 | 73 | 771 | 1029 | 0 | 0 |
| *Prunus* sp.2 | 576583146 | 9719111 | 1.7 | 1806 | 1806 | 1804 | 1804 | 1804 | 1804 | 1 | 20 | 59 | 1095 | 709 | 0 | 0 |
| *Prunus* sp.3 | 409723226 | 8781975 | 2.1 | 1806 | 1806 | 1803 | 1803 | 1803 | 1803 | 1 | 21 | 55 | 1074 | 729 | 0 | 0 |
| *Prunus spinulosa* | 23611766 | 1406518 | 6 | 1802 | 1693 | 1541 | 1478 | 1433 | 1388 | 3 | 202 | 391 | 1099 | 442 | 0 | 0 |
| *Prunus tianshanica* | 594585840 | 12287403 | 2.1 | 1806 | 1806 | 1802 | 1802 | 1802 | 1802 | 1 | 18 | 61 | 985 | 817 | 0 | 0 |
| *Prunus tomentosa* | 207378648 | 5442940 | 2.6 | 1806 | 1806 | 1804 | 1804 | 1804 | 1804 | 1 | 25 | 70 | 1082 | 722 | 0 | 0 |
| *Prunus yunnanensis* | 533773116 | 9861645 | 1.8 | 1806 | 1806 | 1802 | 1802 | 1802 | 1802 | 1 | 12 | 77 | 747 | 1055 | 0 | 0 |
| *Pyrus betulifolia* | 69785618 | 600650 | 0.9 | 1805 | 1714 | 1712 | 1639 | 1382 | 870 | 1 | 3 | 14 | 327 | 1385 | 0 | 0 |
| *Rhaphiolepis umbellata* | 79238440 | 4855418 | 6.1 | 1802 | 1792 | 1742 | 1720 | 1700 | 1633 | 8 | 397 | 657 | 1221 | 521 | 0 | 0 |
| *Rhodotypos scandens* | 217413144 | 9036128 | 4.2 | 1803 | 1796 | 1760 | 1728 | 1681 | 1605 | 0 | 11 | 31 | 1294 | 466 | 0 | 0 |
| *Rosa rugosa* | 670455110 | 4906973 | 0.7 | 1806 | 1800 | 1795 | 1787 | 1752 | 1560 | 1 | 10 | 46 | 602 | 1193 | 0 | 0 |
| *Sibiraea laevigata* | 21529898 | 1063634 | 4.9 | 1801 | 1663 | 1541 | 1502 | 1446 | 1348 | 0 | 7 | 12 | 1386 | 155 | 0 | 0 |
| *Sorbaria sorbifolia* | 39522674 | 2694667 | 6.8 | 1805 | 1717 | 1564 | 1525 | 1492 | 1435 | 1 | 43 | 215 | 982 | 582 | 0 | 0 |
| *Sorbus aucuparia* | 182441736 | 3942377 | 2.2 | 1806 | 1806 | 1802 | 1800 | 1792 | 1730 | 4 | 170 | 866 | 404 | 1398 | 0 | 0 |
| *Spiraea thunbergii* | 46992320 | 629889 | 1.3 | 1805 | 1802 | 1799 | 1795 | 1780 | 1663 | 1 | 19 | 62 | 875 | 924 | 0 | 0 |
| *Vauquelinia californica* | 141008986 | 1163198 | 0.8 | 1806 | 1804 | 1799 | 1766 | 1593 | 1083 | 0 | 0 | 46 | 90 | 1709 | 0 | 0 |
