## Supplemental Table 3 for "Unveiling allopolyploidization-driven genome duplications through progressive analysis of deep genome skimming data"

| Node | Anchor name | Type | Geological period | Assigned data (Mya) | Reference(s) |
|---|---|---|---|---|---|
| F1 | *Prunus hirsutipetala* | fossil | Late Eocene | 37.2-33.9 | Sokoloff DD, Ignatov MS, Remizowa MV, Nuraliev MS, Blagoderov V, Garbout A, Perkovsky EE. 2018. Staminate flower of *Prunus* s. l. (Rosaceae) from Eocene Rovno amber (Ukraine). Journal of Plant Research 131: 925–943. |
| F2 | *Prunus wutuensis* | fossil | Early Eocene | 55 | Li Y, Smith T, Liu C-J, Awasthi N, Yang J, Wang Y-F, Li C-S. 2011. Endocarps of *Prunus* (Rosaceae: Prunoideae) from the early Eocene of Wutu, Shandong Province, China. TAXON 60: 555–564.<br>Mclntosh WC, Chapin CE, Cather SM. 2004. Geochronology of the Central Colorado volcanic field. New Mexico Bureau of Geology and Mineral Resources Bulletin: 205–237. |
| F3 | *Neviusia* | fossil | Early Eocene | 50-49 | Mathews WH. 1964. Potassium-Argon Age Determinations of Cenozoic Volcanic Rocks from British Columbia. GSA Bulletin 75: 465–468. |
| F4 | *Oemleria janhartfordae* | fossil | Early Eocene | 49.96-48.88 | Benedict JC, DeVore ML, Pigg KB. 2011. *Prunus* and *Oemleria* (Rosaceae) Flowers from the Late Early Eocene Republic Flora of Northeastern Washington State, U.S.A. International Journal of Plant Sciences 172: 948–958. |
| F5 | *Amelanchier peritula*<br>*Amelanchier scudderi* | fossil | Late Eocene | 33.9 | Cockerell TDA, Rohwer SA, Rohwer G, Cockerell WP (Wilmatte P), Rush W, Duce T. 1911. Fossil insects from Florissant, Colorado. Bulletin of the AMNH 30: 71–82.<br>MacGinitie HD. 1953. Fossil Plants of the Florissant Beds, Colorado. Carn Inst Wash, Contribut Paleon. 559: 1–198. |
| F6 | *Vauquelinia comptonifolia* | fossil | Middle Eocene | 46.2-40.4 | MacGinitie HD. 1969. The Eocene Green River flora of northwestern Colorado and northeastern Utah. University of California publications in geological sciences 83: 1–140. |
| F7 | *Spiraea* sp. | fossil | Early Eocene | 50-49 | Wehr Wesley C, Hopkins Donald Q. 1994. The Eocene orchards and gardens of Republic, Washington. Washington Geology 22: 27–34. |
| F8 | *Holodiscus lisii* | fossil | Late Eocene | 34 | Schorn HE. 1998. *Holodiscus lisii* (Rosaceae): a new species of ocean spray from the late Eocene Florissant Formation, Colorado, USA. PaleoBios 18: 21–24. |
| C1 | *Amygdaloideae* stem | secondary calibration | Cretaceous | 96.36-94.46 | Zhang SD, Jin JJ, Chen SY, Chase MW, Soltis DE, Li H-T, Yang J-B, Li D-Z, Yi T-S. 2017. Diversification of Rosaceae since the Late Cretaceous based on plastid phylogenomics. New Phytologist 214: 1355–1367. |
